# Genomic Analysis of Progenitors in Viral Infection Implicates Glucocorticoids as Suppressors of Plasmacytoid Dendritic Cell Generation

**DOI:** 10.1101/2024.10.28.620771

**Authors:** Yeara Jo, Trever T. Greene, Kai Zhang, Carolina Chiale, Ziyan Fang, Simone Dallari, Nuha Marooki, Wei Wang, Elina I. Zuniga

**Author notes:** These authors contributed equally (Trever T. Greene, Kai Zhang). **Author Contributions:** Y.J. designed, performed, analyzed, and interpreted all experiments and wrote the manuscript. T.T.G, C.C. and Z.F. helped perform and interpret some experiments and contributed to writing the manuscript. K.Z. and W.W. designed the pipeline and provided advice for processing bioinformatics data. S.D. and N.M. obtained and processed samples needed for some experiments. E.I.Z. conceived and supervised the project, designed and interpreted experiments, and wrote the manuscript. **Competing Interest Statement:** E.I.Z. is in the Scientific Advisory Board of Primmune and AGS Therapeutics. Other authors declare no competing interests. **Classification:** Biological Sciences, Immunology and Inflammation.

## Abstract

Plasmacytoid Dendritic cells (pDCs) are the most potent producers of interferons, which are critical antiviral cytokines. pDC development is, however, compromised following a viral infection, and this phenomenon, as well as its relationship to conventional (c)DC development is still incompletely understood. By using lymphocytic choriomeningitis virus (LCMV) infection in mice as a model system, we observed that DC progenitors skewed away from pDC and towards cDC development during *in vivo* viral infection. Subsequent characterization of the transcriptional and epigenetic landscape of fms-like tyrosine kinase 3^+^ (Flt3^+^) DC progenitors and follow-up studies revealed increased apoptosis and reduced proliferation in different individual DC-progenitors as well as a profound IFN-I-dependent ablation of pre-pDCs, but not pre-DC precursor, after both acute and chronic LCMV infections. In addition, integrated genomic analysis identified altered activity of 34 transcription factors in Flt3^+^ DC progenitors from infected mice, including two regulators of Glucocorticoid (GC) responses. Subsequent studies demonstrated that addition of GCs to DC progenitors led to downregulated pDC-primed-genes while upregulating cDC-primed-genes, and that endogenous GCs selectively decreased pDC, but not cDC, numbers upon *in-vivo* LCMV infection. These findings demonstrate a significant ablation of pre-pDCs in infected mice and identify GCs as suppressors of pDC generation from early progenitors. This provides an explanation for the impaired pDC development following viral infection and links pDC generation to the hypothalamic-pituitary-adrenal axis.

**Significance Statement:** Plasmacytoid dendritic cells (pDCs) play critical roles in antiviral responses. However, adaptations of DC progenitors lead to compromised pDC generation after viral infection. Here, we characterized the transcriptional and epigenetic landscapes of DC progenitors after infection. We observed widespread changes in gene expression and chromatin accessibility, reflecting shifts in proliferation, apoptosis, and differentiation potential into various DC subsets. Notably, we identified alterations in the predicted activity of 34 transcription factors, including two regulators of glucocorticoid responses. Our data demonstrate that glucocorticoids inhibit pDC generation by reprogramming DC progenitors. These findings establish a molecular framework for understanding how DC progenitors adapt to infection and highlight the role of glucocorticoid signaling in this process.

## Introduction

Immunosuppression is a common feature of sustained viral infections, which are a massive burden on human health, impacting millions of lives each year ^1–3^. While this immunosuppression is detrimental for host defense against secondary pathogens, it also represents a strategy from the host to avoid overt pathology, choosing coexistence with the pathogen over mutual destruction. Indeed, genetic ablation of key immunosuppressive molecules is highly detrimental, and in many cases lethal, to hosts challenged with what would otherwise be a protracted viral infection ^2; 3^. In contrast, transient interruption of these host adaptations has been highly successful as a therapeutic strategy in the treatment of chronic infections ^3; 4^ or other long-term diseases such as cancer ^5^.

Much of the work on immune adaptation during persistent infections focuses directly on changes to the adaptive immune system (e.g. T and B cells). However, innate immune responses are also impacted by sustained infections. Particularly, dendritic cells (DCs), a crucial bridge between innate and adaptive immunity which shepherd T and B cell responses throughout infection ^6^, also adapt their function to sustained insults ^7; 8^. Furthermore, we have recently shown that some of these adaptations act to mitigate infection-associated pathology ^9^. However, unlike T and B cells, DCs are short-lived ^6^ and therefore stable DC populations rely on continuous development from progenitors ^6^. It is therefore likely, that some DC changes after infection are coordinated with DC progenitors’ adaptations. Indeed, DC progenitors from mice infected with lymphocytic choriomeningitis virus (LCMV) show compromised generation of pDCs both *ex vivo* and after *in vivo* cell transfer ^10^. Still, how DC progenitor populations change during persistent infection, and how this impacts the development of distinct DC types is not yet well understood.

Immediate clonogenic precursor populations that derive only one type of DC have been found for most DC types (e.g., conventional (c)DC1 ^11; 12^, cDC2A ^13^, cDC2B ^13^, plasmacytoid (p)DC ^14–16^. However, the preceding developmental stages are not as well defined. For example, common dendritic cell progenitors (CDP) give rise to cDC1, cDC2, and pDC ^17^. However, pDC development has also been described to originate from common lymphoid progenitors (CLP), leading to arguments about their classification as DCs or innate lymphocytes ^18–20^. Furthermore, while CDP can be divided into subsets with different skews toward cDC1 or cDC2 development, even these populations have drastically varying potential for DC subtype development when clonally isolated ^14^.

Finally, there is evidence that cDC1 and pDC, but not cDC2 share a clonogenic precursor^16^. Altogether, this has led some to favor a more diffuse and continuous model for early DC development (^21^, Fig. 1A), in which a larger bounding population with DC potential Lin^-^c-Kit^int/lo^Flt3^+^ population contains a spectrum of cells with individually varying developmental potential that does not necessarily divide cleanly along available surface marker definitions.

**Figure 1.**
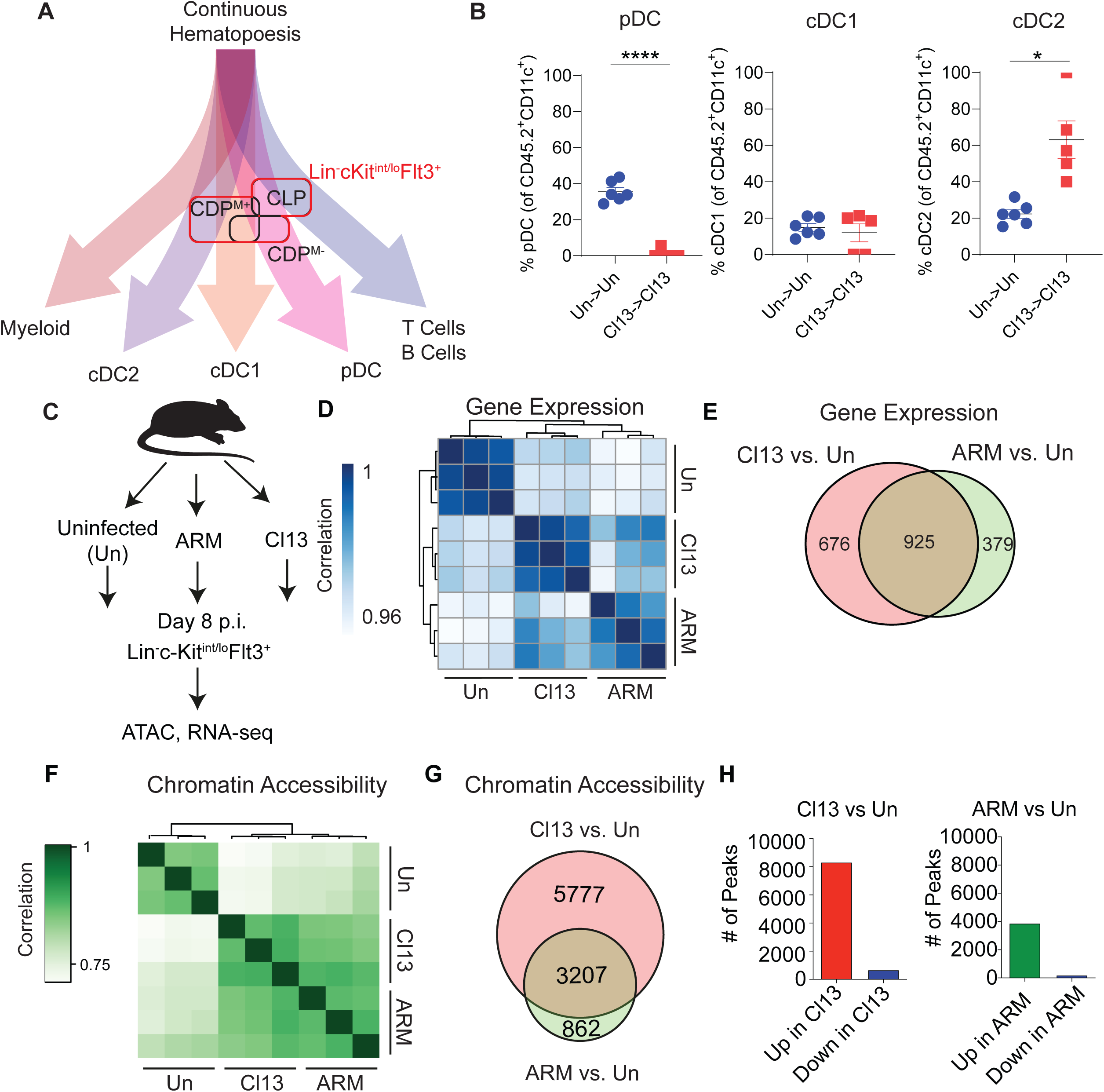
Acute and chronic LCMV infections induced overlapping changes in transcriptomes and chromatin landscapes. (A) Conceptual and non-proportional diagram of the continuous model of DC hematopoiesis. Lin^−^c-kit^int/lo^Flt3^+^ progenitors are outlined in red. Classically defined progenitor populations (CD115+ CDP (CDP^M+^), CD115-CDP (CDP^M-^), CLP, black boxes) partially overlap in potential. (B) Lin^−^c-kit^int/lo^Flt3^+^ progenitors were isolated from CD45.2^+^ mice infected with LCMV Cl13 at day 8 p.i., or mice that were left uninfected (Un) and transferred into infection matched or Un controls, respectively. Eight days after transfer, spleens were harvested, and DC development assessed by flow cytometry. (C-H) C57BL6/J mice were infected with LCMV ARM or Cl13 or left uninfected (Un), sacrificed at day 8 p.i., and Lin^−^c-kit^int/lo^Flt3^+^ progenitors were FACS-purified from BM for RNA-seq (D-E) and ATAC-seq (F-H) analyses. Data are representative of three independent repeats, each with 3-5 mice pooled per group. (D) Hierarchical clustering of RNA-seq profiles on the whole transcriptome by using Pearson correlation. (E) Venn diagram showing overlap between differentially expressed genes (DEGs) from Cl13 vs. Un and ARM vs. Un comparisons. (F) Hierarchical clustering of ATAC-seq profiles by using Pearson correlation. (G) Venn diagram showing overlap between differentially accessible (DA) chromatin regions from Cl13 vs. Un and ARM vs. Un comparisons. (H) Number of differential peaks that opened (left, red; right, green) or closed (blue) during infection. Graphs depict means ± SEM and symbols represent individual mice. (B) Data are pooled from two independent experiments with 2– 4 mice/group. (D-H) Data are representative of three independent repeats, each with 3-5 mice pooled per group. * p < 0.05, ****p < 0.0001. Statistical significance was determined by unpaired student’s t-test (B).

We previously found that Lin^-^c-Kit^int/lo^Flt3^+^ progenitors are numerically reduced and have reduced capacity to generate pDCs after infection ^10^. Importantly, we found that this was true after both acute and persistent viral infection, though this effect was longer lasting in the chronic setting. The mechanisms underlying such DC progenitor adaptation, however, remain poorly understood.

A growing body of evidence supports an active role of chromatin remodeling, in addition to gene expression differences, in regulating multiple aspects of immune responses, including function and development of immune cells ^22; 23^. Chromatin remodeling enables transcription factors (TFs) to bind to accessible chromatin regions and drive differential expression of their target genes ^24^. Thus, integration of TF levels, target gene expression, and chromatin accessibility represents a powerful approach to identify regulators of immune responses. To generate hypotheses on the drivers of infection induced Flt3^+^ progenitor adaptations that result in reduction in their numbers and defects in their capacity to generate DCs, we determined the transcriptional and chromatin landscapes of the Flt3^+^ progenitor population isolated from the bone marrow (BM) of LCMV-infected mice. We found that, in contrast to T cells ^25; 26^, infection with acute (Armstrong 53b, ARM) or persistent (Clone 13, Cl13) LCMV variants induced largely overlapping transcriptomes and chromatin landscapes in Lin^−^c-kit^int/lo^Flt3^+^ progenitors. Follow-up studies elucidated the differential involvement of altered proliferation and apoptosis in individual progenitor sub-populations (i.e. CLP, CDP CD115^+^ and CDP CD115^-^) included in our genomics analysis, providing an explanation for their reduced numbers ^10; 27^. We also revealed that pre-pDCs, which are pDC-committed early progenitors^14; 15^, were depleted in an IFN-I-dependent manner after infection, consistent with the IFN-I-dependent impairment in pDC development that we previously reported^10^. We then used an algorithm that integrates transcriptome and chromatin accessibility analysis ^28^ to identify potential regulators of DC development during infection. Altogether, we identified 34 TFs with altered predicted activity in progenitors from infected mice, including 23 TFs with no previous connection to DC biology. Among them, we found several TFs that regulate responses to glucocorticoids (GC), including GC Receptor (GR, encoded by Nr3c1) itself. Subsequent *in vitro* investigation revealed that a short-term spike of the GC corticosterone was sufficient to downregulate pDC-primed genes in early progenitors and reduce pDC development, biasing DC progenitors to the cDC2 fate similar to the bias observed after *in vivo* transfer. Consistently, we found that mice deficient in GC production (adrenalectomized), which had similar pDC numbers in uninfected conditions, exhibited greater numbers of pDCs (but not cDC) following infection. Together, our results revealed important mechanistic insights on the adaptations of DCs and their progenitors after infection, including: i) increased apoptosis or decreased proliferation underlying the reduction of individual progenitors with potential to generate DCs, ii) identification of several TFs with potential to regulate DC development, iii) a new effect of GC in downregulating pDC-primed-genes in early DC progenitors, iv) a selective GC-mediated effect on pDC generation *in vitro* and pDCs number upon *in vivo* infection, linking DC progenitor adaptations with the hypothalamic-pituitary-adrenal (HPA) axis.

## Results

### Acute and chronic LCMV infections drove largely overlapping changes in transcriptomes and chromatin accessibility of Flt3 ^+^ progenitors

All the precursors with potential to generate DCs are encompassed within the Flt3^+^ progenitor population ^29^. Specifically, Flt3^+^ lineage^-^ progenitors incorporate different subpopulations that have been previously demonstrated to contribute to the generation of DCs, including common lymphoid progenitors (CLPs) which are progenitors of lymphocytes, but when considering DC potential primarily produce pDCs although they can also produce cDC1 and cDC2 cells ^14; 15; 30–32^, CD115^-^ common dendritic cell progenitors (CDPs) which produce pDC, cDC1, and cDC2 but are also biased towards pDC generation ^17; 33; 34^, and CD115^+^ CDPs which produce pDC, cDC1, and cDC2 but skewed toward cDC development ^17; 33; 34^. Importantly, we and others previously reported that Flt3^+^ lineage^-^ progenitors are reduced in numbers ^10; 27^ and exhibit a profoundly compromised capacity to generate pDCs after adoptive cell transfer at the chronic phase (i.e. day 30 post-infection (p.i.)) of LCMV Cl13 infection ^10^. To investigate how these progenitors change their pDC developmental capacity at an earlier time point, during the acute phase of LCMV Cl13 infection, and how this compares to generation of cDC1 and cDC2 subsets, we isolated Lin^−^c-kit^int/lo^Flt3^+^ progenitors from uninfected mice (Un) or LCMV Cl13 (Cl13) infected mice at 8 day p.i., and transferred them into congenically distinct (CD45.1) uninfected or infection matched recipients, respectively. In line with our previous observations at day 30 p.i., we found that transferred Lin^−^c-kit^int/lo^Flt3^+^ progenitors were severely deficient in their capacity to develop into pDCs at day 8 p.i. (Fig. S1A, Fig. 1B). Intriguingly, there was a relative increase in the derived cDC2, but not cDC1, percentages with the donor-derived DC population (Fig. S1A, Fig. 1B). Given the importance of the cytokine Flt3 ligand (Flt3L) in the development of pDCs and cDCs (cDC1s in particular) from Lin^−^c-kit^int/lo^Flt3^+^ progenitors^35–37^, we asked if altered DC development after infection is caused by changes in the systemic levels of Flt3L during infection. At day 8 p.i., serum levels of Flt3L measured by ELISA were significantly higher in Cl13-infected, but not ARM-infected, mice compared to uninfected mice (Fig. S1B). We also measured the systemic levels of Leukemia Inhibitory Factor (LIF), which has been recently reported to inhibit pDC development^38^, by ELISA. However, at day 8 p.i., LIF was not detectable in serum from any of the mice examined, regardless of infection status (data not shown). These data suggested that changes in DC development after LCMV infection are mediated by mechanisms other than shortage of Flt3L or increase in LIF.

Next, as an initial approach to generate hypotheses on the mechanisms underlying changes in DC development after LCMV infection, we chose to compare the transcriptomes and the chromatin landscapes of Lin^−^c-kit^int/lo^Flt3^+^ progenitors from mice infected with acute (ARM) or chronic (Cl13) LCMV variants to their counterparts from uninfected mice (Fig. 1C). Lin^−^c-kit^int/lo^Flt3^+^ progenitor cells, isolated with > 95% purity as shown in Fig. S1C, were analyzed at day 8 p.i. when the progenitors from both ARM- and Cl13-infected mice are defective for pDC generation and when their phenotypes deviate the most from their counterparts from uninfected mice ^10^.

Principal component analysis (PCA) and hierarchical clustering of RNA-seq profiles between samples revealed that Lin^−^c-kit^int/lo^Flt3^+^ progenitors from uninfected mice had distinct gene expression, whereas their counterparts from ARM- and Cl13-infected mice were more closely related to each other (Fig. 1D and Fig. S1D). Approximately 71% (925) of the differentially expressed genes (DEGs) between progenitors from ARM-infected and uninfected mice were also differentially expressed upon Cl13 infection (Fig. 1E and Table S1& S2), supporting the distinct but highly overlapping transcriptomic changes of Lin^−^c-kit^int/lo^Flt3^+^ progenitors in response to the two infections. In agreement with our clustering analysis, we observed fewer DEGs and more modest differences in expression levels when comparing ARM to Cl13 infection relative to our comparisons of ARM- or Cl13-infected mice to their uninfected counterparts (Fig. S1E).

Consistently, evaluation of open chromatin regions via assay for transposase-accessible chromatin using sequencing (ATAC-seq) ^39^ followed by PCA and hierarchical clustering revealed that the chromatin landscapes of Lin^−^c-kit^int/lo^Flt3^+^ progenitors from ARM- and Cl13-infected mice were more closely related to each other vs. their counterparts from uninfected mice (Fig. 1F and Fig. S1F). Indeed, approximately 79% (3207) of the differentially accessible (DA) chromatin regions in progenitors from ARM-infected vs. uninfected mice were also DA in progenitors from Cl13-infected vs. uninfected mice (Fig. 1G). Furthermore, we found no significantly DA region when progenitors from ARM- and Cl13-infected mice were directly compared (not shown), suggesting a high degree of similarity in chromatin remodeling of Lin^−^c-kit^int/lo^Flt3^+^ progenitors after both acute and chronic LCMV infections. Notably, over 90% of DA regions became more accessible upon either ARM or Cl13 infection, indicating that Lin^−^c-kit^int/lo^Flt3^+^ progenitors underwent large-scale and mostly overlapping gains, rather than losses, in chromatin accessibility in response to both acute and chronic infection (Fig. 1H). Furthermore, regions that became DA in progenitors from ARM- or Cl13-infected mice were mostly found distal (>1kb) to transcription start sites (Fig. S1G), suggesting that opening of distal regulatory elements were primarily involved in shaping the adaptations in progenitors’ chromatin landscapes upon infection.

Taken together these data demonstrated that LCMV ARM and Cl13 infections drove distinct but highly overlapping gene expression changes in Lin^−^c-kit^int/lo^Flt3^+^ progenitors and induced large-scale gains in their chromatin accessibility, which were not significantly different between the two infections. Although such transcriptional and chromatin changes may result from an altered composition of the individual subpopulations encompassed within the Lin^−^c-kit^int/lo^Flt3^+^ progenitors, our data suggest that, if such, those subpopulation changes are expected to be quite similar after acute and chronic LCMV infections.

### Progenitors with potential to generate DCs exhibited differential alterations in proliferation and apoptosis after LCMV infection

Given that we previously observed similarly reduced Lin^−^c-kit^int/lo^Flt3^+^ progenitors at early stages of both acute and chronic LCMV infections ^10^, we sought to gain insight on biological pathways that may underlie their numerical reduction.

For this, we subjected genes and chromatin regions that were similarly regulated upon both ARM and Cl13 infections to over-represented pathway analysis (ORA) in WebGestalt using the GO Biological Processes database ^40^ and Genomic Regions Enrichment of Annotations Tool (GREAT) ^41^, respectively. We found that, in addition to the expected alterations in pathways related to immune responses, there were alterations in cell cycle and apoptosis related pathways in the progenitors from infected vs. uninfected mice (Fig. S1H and I). This led us to hypothesize that changes in proliferation and/or apoptosis induced upon infection may drive the numerical reduction that we previously reported in the individual DC progenitors encompassed within Lin^−^c-kit^int/lo^Flt3^+^ population ^10^. To test this hypothesis, we measured BrdU incorporation and active caspase 3 in CLPs, CD115^+^ and CD115^-^ CDPs from uninfected, ARM-infected, and Cl13-infected mice. We observed that CLPs derived from ARM- and Cl13-infected mice exhibited reduced proliferation compared to their counterparts from uninfected mice, as indicated by lower frequency of cells that incorporated BrdU *in vivo* (Fig. 2A), while their apoptosis levels were not significantly changed (Fig. 2B). On the other hand, we noted enhanced proliferation in CD115^-^ CDPs derived from Cl13-infected, but not ARM-infected, mice (Fig. 2C), while apoptosis levels in these cells were increased after both ARM and Cl13 infections (Fig. 2D). Interestingly, we did not observe any significant difference in proliferation or apoptosis between CD115^+^ CDPs derived from ARM- or Cl13-infected mice versus their counterparts from uninfected mice (Fig. 2E and F). As impaired retention of DC progenitors in the BM can also contribute to their numerical reduction, we investigated regulation of CXCR4, a chemokine receptor that mediates the retention of CDPs in the BM^42^, in Lin^−^c-kit^int/lo^Flt3^+^ progenitors from infected mice. CXCR4 protein levels were increased in progenitors from ARM-infected and, to a larger extent, from Cl13-infected mice (Fig. S1J), suggesting that defective retention of BM progenitors may not be a major cause of their numerical reduction after LCMV infection.

**Figure 2.**
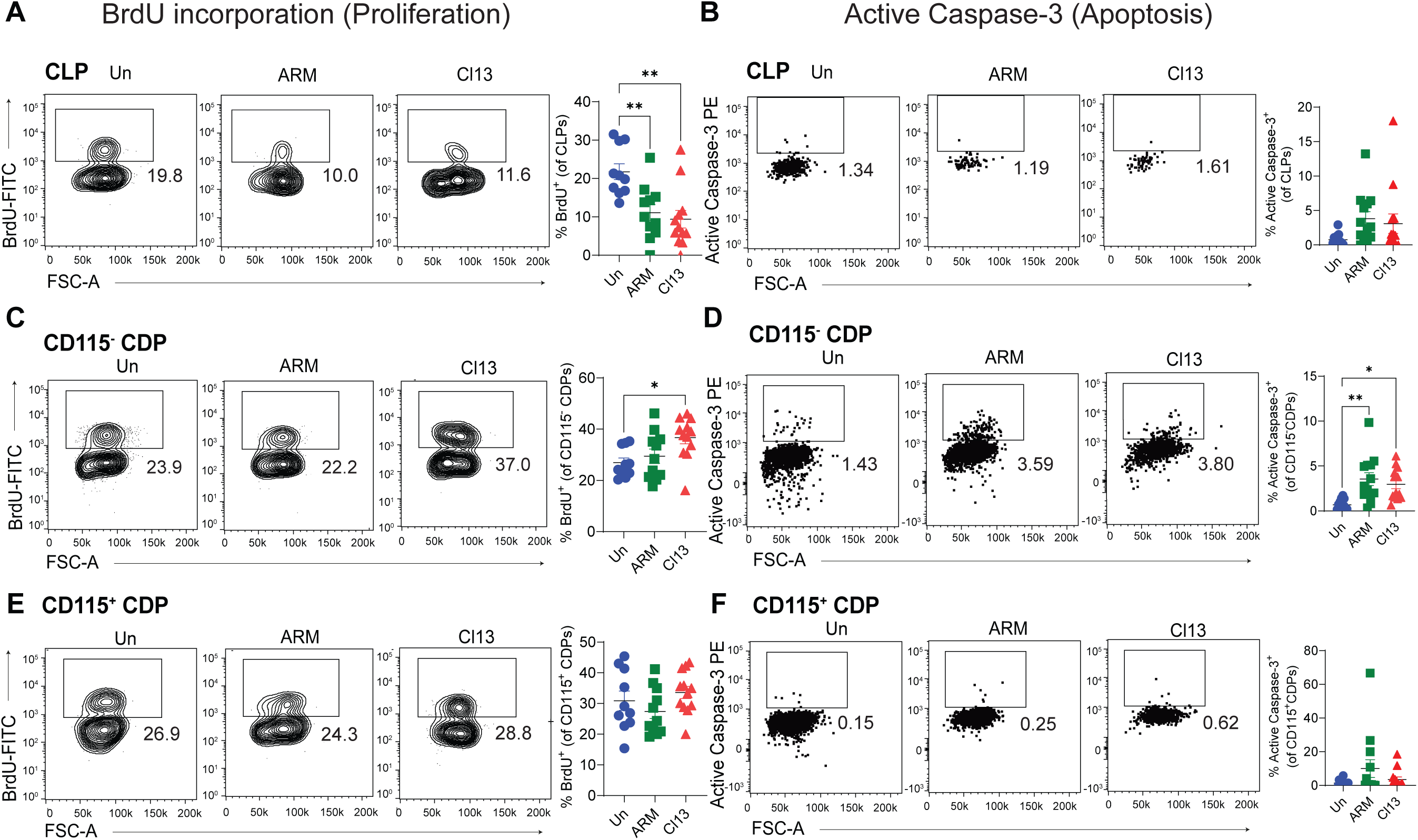
DC progenitors showed changes in proliferation and apoptosis following infection. C57BL6/J mice were infected with LCMV ARM or Cl13 or left uninfected (Un) for 8 days. CLPs (A-B), CD115^-^ CDPs (C-D), and CD115^+^ CDPs (E-F) from uninfected (blue). ARM-infected (green) and Cl13-infected (red) mice were analyzed for *in vivo* BrdU incorporation (A, C, and E) and exhibition of active caspase 3 (B, D, and F). Representative flow cytometry plots are shown. Graphs depict means ± SEM and symbols represent individual mice. Data are pooled from three independent experiments with 4–5 mice/group. ** p < 0.01, ***p < 0.001. Statistical significance was determined by one-way ANOVA with Tukey’s multiple comparisons test (A-F).

Taken together, these results provide a mechanistic explanation for the numerical reduction of CLPs and CD115^-^ CDPs that we previously reported after LCMV infection ^10^, indicating that reduced proliferation at least partially accounted for the CLP reduction while increased apoptosis explained the decrease in the number of CD115^-^ CDPs. Finally, our data suggest that the numerical reduction of CD115^+^ CDPs previously reported in LCMV-infected mice ^10^ is probably mediated by mechanisms unrelated to changes in proliferation, apoptosis, or CXCR4-mediated retention in the BM.

### LCMV infected mice exhibited depletion of pre-pDCs and bias towards pre-DCs

Given the identification of pre-pDC and pre-DC progenitors as fully committed pDC and cDC lineage precursors, respectively ^14; 15^, we sought to analyze the aforementioned genomic dataset in the light of these findings. We therefore compared the genes that were differentially expressed in Lin^−^c-kit^int/lo^Flt3^+^ progenitors from ARM and Cl13 infected vs. uninfected mice to the gene signatures reported for pre-pDC or pre-DC priming ^14^. We observed that both infections drove a dramatic bias against pre-pDC-primed signature genes, with downregulation of 117 (96.7%) out of the 121 differentially regulated pre-pDC-primed genes in Lin^−^c-kit^int/lo^Flt3^+^ progenitors from infected vs. uninfected mice (Fig. 3A, left). In contrast, we found that 80 of the 85 (94.1%) pre-DC-primed signature genes differentially regulated in Lin^−^c-kit^int/lo^Flt3^+^ progenitors were significantly upregulated upon infection (Fig. 3A, right). Subsequent gene set enrichment analysis (GSEA) validated these findings by showing that genes that were significantly downregulated in Lin^−^c-kit^int/lo^Flt3^+^ progenitors from infected mice were enriched for the pre-pDC-primed signature (Fig. 3B, top), while upregulated genes were enriched for the pre-DC-primed signature (Fig. 3B, bottom).

**Figure 3.**
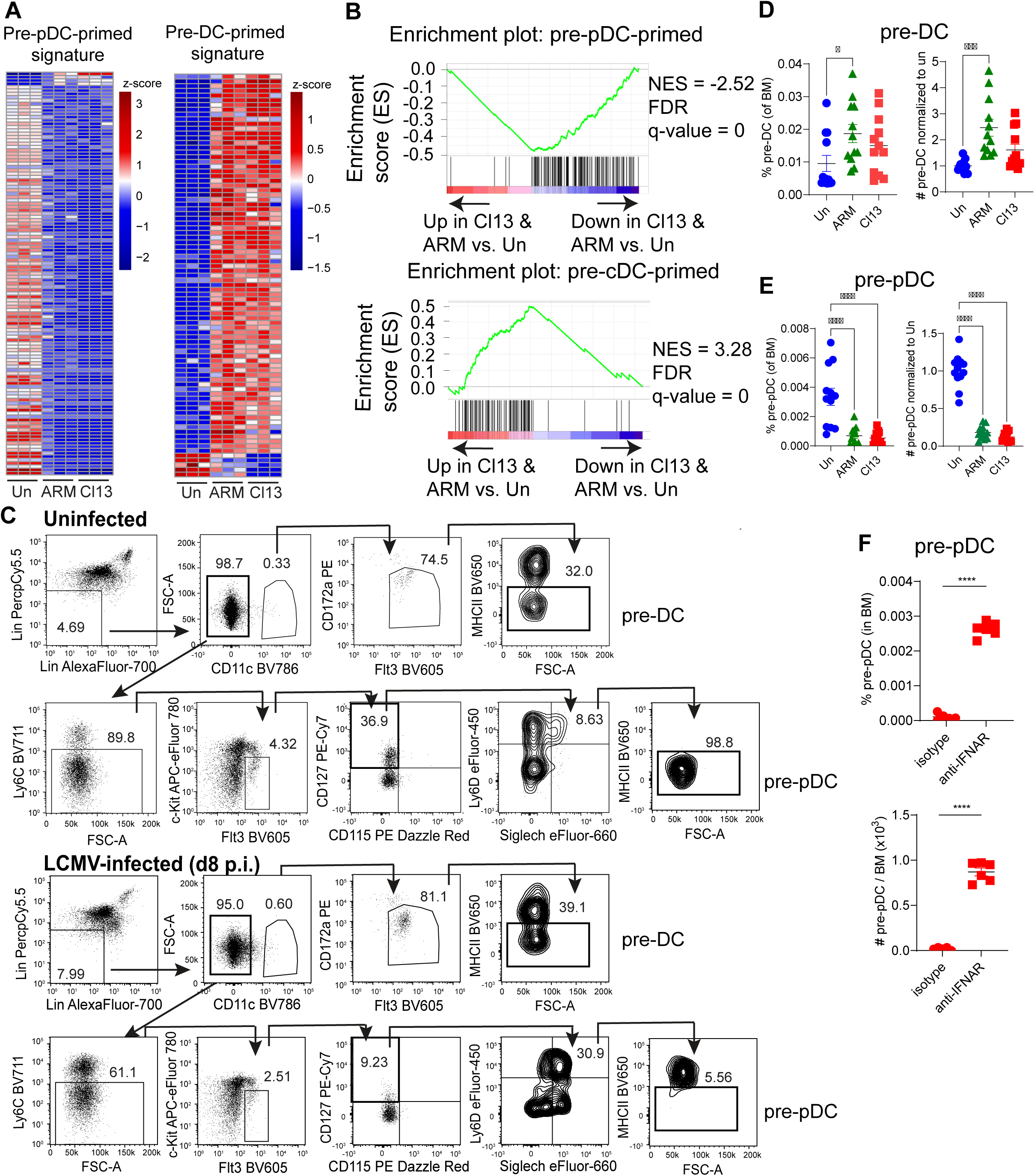
LCMV infected mice exhibited depletion of pre-pDCs and enrichment of pre-DCs. C57BL6/J mice were infected with LCMV ARM or Cl13 or left uninfected (Un) for 8 days. (A-B) RNA was isolated from FACS-purified Lin^−^c-kit^int/lo^Flt3^+^ progenitors and subjected to RNA-seq analyses. Data are representative of three independent repeats, each with 3-5 mice pooled per group. (A) Heatmaps for relative expression of DEGs in both ARM and Cl13 infected vs. uninfected mice that were defined as pre-pDC-primed (left) or pre-DC-primed (right) signature in Dress, et al. 2019. (B) Gene set enrichment analysis (GSEA) and normalized enrichment score (NES) of infection-specific DEGs for pre-pDC-primed (top) or pre-DC-primed (bottom) signature in Dress, et al. 2019. (C) Gating strategy and representative flow cytometric plots for pre-pDCs and pre-DCs. (D-E) Frequency (left) and numbers (right) of pre-DCs (D) and pre-pDCs (E) in BM. Numbers were normalized to the average of numbers in uninfected mice in the same experiment. (F) Mice were treated with isotype or anti-IFNAR blocking antibody beginning 1 day before and throughout LCMV Cl13 infection. Frequency (top) and numbers (bottom) of pre-pDCs were quantified. (D-F) Graphs depict mean ± SEM and symbols represent individual mice. Data are representative of 2 independent experiments (F) or pooled from 3 (C-E) independent experiments. *<0.05, ***p < 0.001, ****<0.0001. Statistical significance was determined by one-way ANOVA with Tukey’s multiple comparisons test (D, E) or unpaired student’s t-test (F).

The aforementioned analysis raised the hypothesis that pre-pDCs (but not pre-DCs) might be suppressed after LCMV infection. To directly test this hypothesis, we quantified the pre-pDCs and pre-DCs from uninfected, ARM-infected, and Cl13-infected mice via flow cytometry, as previously described ^12; 14; 15^ (Fig. 3C). Consistent with our transcriptomic analysis, we observed that while the percentage and numbers of pre-DCs were significantly enhanced after ARM infection and exhibited an increasing trend after Cl13 infection (Fig. 3D), the frequency and absolute number of pre-pDCs were significantly reduced and barely detectable after both ARM and Cl13 infections (Fig. 3E). Interestingly, IFN-I, which suppresses pDC development after infection ^10^, contributed to the reduced numbers of pre-pDCs, as treatment of mice with anti IFN-I receptor (IFNAR) blocking antibody throughout the Cl13 infection restored the frequency and numbers of pre-pDCs (Fig. 3F).

Overall, these results indicated that, while the cDC progenitor program might be favored, there is a clear and profound IFN-I dependent suppression of pDC-committed progenitors after viral infection. Taken together, our data revealed a profound loss of committed pDC progenitors (i.e. pre-pDCs) with a concomitant trend towards increased committed cDC progenitors.

### Integrated transcriptomic and chromatin analysis in DC progenitors highlighted both known and novel TF candidate regulators after acute and chronic LCMV infection

We then sought to identify potential regulators that might contribute to the suppression of pDC generation and/or the bias toward cDC generation after viral infection. In this regard, we previously reported that the expression of TFs that promote pDC generation, including Tcf4 ^43^, SpiB ^44^, and Bcl11a ^45^, are reduced in Lin^−^c-kit^int/lo^Flt3^+^ progenitors from LCMV Cl13-infected mice compared to those from uninfected mice ^10^. Additionally, progenitors from both ARM- and Cl13-infected mice exhibited reduced expression of Runx2, a TF that also promotes pDC generation^46^ (Fig. S2A). In contrast, consistent with bias towards cDC development after infection, expression of Batf2, a TF that can promote cDC1 development by compensating for Batf3^47^, was significantly upregulated in DC progenitors from both ARM- and Cl13-infected mice (Fig. S2A). While differential expression of TFs and their target genes can provide insights into alterations in the development and function of cells during infection, the expression level of a TF does not always correlate with its activity ^48^. Therefore, in order to identify TFs with differential activity upon infection that could contribute to the suppressed pDC development after infection, we used Taiji, a software package that integrates ATAC-seq and RNA-seq data and performs personalized PageRank analysis to assess the global importance of TFs in a defined biological condition ^28^. Taiji has been successfully used to find novel TF regulators in adaptive immune cells such as CD8^+^ T cells ^49; 50^ and B cells ^51^. By implementing Taiji for the first time in DC progenitors, we identified 34 TFs with predicted altered activities in Lin^−^c-kit^int/lo^Flt3^+^ progenitors from both ARM- and Cl13-infected mice compared to their counterparts from uninfected mice (more than 2-fold change in rank and above the minimum expression level threshold (Transcripts Per Million (TPM) > 1)) (Table 1). Out of these 34 TFs, there were 12 TFs with aligned changes in expression and activity in progenitors from both ARM- and Cl13-infected vs. uninfected mice (Fig. 4A, S2B and C, blue dots). Remarkably, 21 out of the 34 (61.8%) TFs with differential activity upon infection did not show any variation in expression upon infection or were differentially expressed upon only one type of infection (Fig. 4A, S2B and C, black dots), while 1 TF with modest differential expression showed predicted activity changes in the opposite direction (Fig. 4A, S2B and C, magenta dot). Interestingly, 11 out of the aforementioned 34 (32.4%) TFs with differential predicted activity upon infection have been previously associated with DC development and/or function ^43; 52–64^ (Table 1 and named TFs in Fig. 4A, S2B and C), while the remaining 23 TFs have not yet been associated with DC biology.

**Figure 4.**
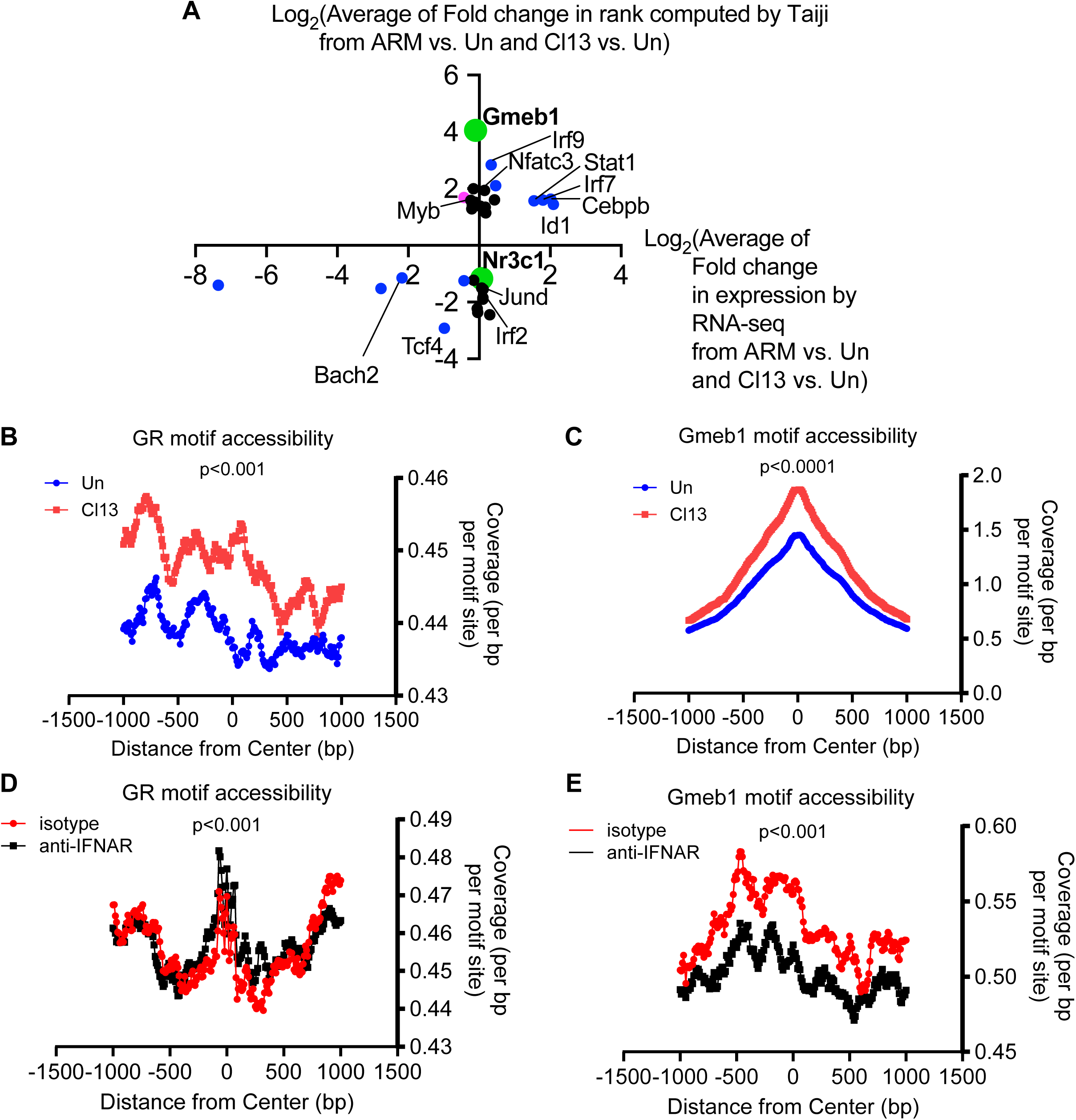
Infection altered predicted activity of multiple TFs in DC progenitors from infected mice, among which GR and GMEB1 exhibited enhanced chromatin accessibility. C57BL6/J mice were infected with LCMV ARM or Cl13 or left uninfected (Un) and sacrificed at day 8 p.i. (A) Lin^−^c-kit^int/lo^Flt3^+^ progenitors were isolated from BM by flow cytometry for RNA-seq and ATAC-seq analyses, which were subsequently used in Taiji analysis. Scatter plot showing correlation between fold change in expression quantified by RNA-seq and fold change in rank computed by Taiji of TFs predicted by Taiji to have altered activities in progenitors from ARM- and Cl13-infected vs. uninfected mice. Average of fold changes in ARM vs. uninfected and Cl13 vs. uninfected comparisons was log2-transformed to generate the plot. Differential expression was defined by DESeq2 with the threshold of FDR < 0.05. Data are representative of three independent repeats, each with 3-5 mice pooled per group. (B-E) Predicted GR (B and D) and GMEB1 (C and E) motif sites were curated by scanning the GR and GMEB1 position matrix from the MEME suite across the genome independently of Taiji with p-value cutoff of 1E-5. Coverages of these sites by sequence tags identified from ATAC-seq in uninfected vs. Cl13-infected mice (B, C) or Cl13-infected mice treated with isotype controls or anti-IFNAR nAb (D, E) are plotted. Statistical significance was Wilcoxon signed rank test (B-E).

**Table 1.** Table of key TFs with higher or lower activity in BM DC progenitors from LCMV ARM- and Cl13-infected (day 8 p.i.) compared to uninfected mice. Key TFs were filtered using three criteria: 1) Average rank of all conditions is greater than 0.0001. 2) The fold change between progenitors from LCMV-infected and uninfected mice is over 2 in both ARM vs. uninfected and Cl13 vs. uninfected comparisons. 3) The expression level of TF passes minimum expression threshold (average Transcripts per Million (TPM) across 3 replicates >1 in progenitors from uninfected mice).* denotes previous link with DC development or function in the cited references.

| TF regulators with higher activity in ARM and CI13 vs. uninfected | Fold change (rank ARM/rank uninfected) | Fold change (rank CI13/rank uninfected) | Average(rank ARM/rank uninfected, rank CI13/rank uninfected) | TF regulators with lower activity in ARM and CI13 vs. uninfected | Fold change (rank ARM/rank uninfected) | Fold change (rank CI13/rank uninfected) | Average(rank ARM/rank uninfected, rank CI13/rank uninfected) |
| --- | --- | --- | --- | --- | --- | --- | --- |
| Gmeb1 | 25.32617709 | 7.904396518 | 16.6152868 | Tcf4*<br>*Cisse, et al. 2008*<br>*Nagasawa, et al. 2008* | 0.123733291 | 0.140109039 | 0.131921165 |
| Irf9<br>*Huber, et al. 2017* | 3.405555657 | 10.85500811 | 7.130281881 | Zfp524 | 0.173637009 | 0.192936942 | 0.183286976 |
| Atf7 | 3.003552737 | 5.573331714 | 4.288442225 | Mbd1 | 0.254206244 | 0.134761543 | 0.194483894 |
| Nfatc3*<br>*Bao, et al. 2016* | 4.916336111 | 3.061921114 | 3.989128613 | Cic | 0.194181542 | 0.233615841 | 0.213898691 |
| Cebpg | 4.753076932 | 2.917606929 | 3.83534193 | Mecp2 | 0.11088892 | 0.426478628 | 0.268683774 |
| Elk3 | 2.773475208 | 3.651880771 | 3.212677989 | Irf2*<br>*Ichikawa, et al. 2004*<br>*Honda, et al. 2004* | 0.07139777 | 0.484175682 | 0.277786726 |
| Cebpb*<br>*Bornstein, et al. 2014*<br>*Scholz, et al. 2020* | 3.148771052 | 3.052562014 | 3.100666533 | E2f4 | 0.256447927 | 0.430092364 | 0.343270146 |
| Mtf1 | 3.049726935 | 3.042644903 | 3.046185919 | Glis2 | 0.315493832 | 0.379911586 | 0.347702709 |
| Irf7*<br>*Owens, et al. 2012*<br>*Bao, et al. 2016* | 2.74321694 | 3.305625578 | 3.024421259 | Jund*<br>*Hara, et al. 2013* | 0.253922584 | 0.452430827 | 0.353176705 |
| Myb*<br>*Papathanasiou, et al. 2017* | 2.269600826 | 3.734257407 | 3.001929117 | Hnf4a | 0.391732642 | 0.365768461 | 0.378750552 |
| Stat1*<br>*Jackson, et al. 2004*<br>*Longman, et al. 2007* | 2.004484922 | 3.927104101 | 2.965794512 | Zkscan1 | 0.364152907 | 0.477091913 | 0.42062241 |
| Zbtb6 | 3.504869419 | 2.241278505 | 2.873073962 | Hoxa3 | 0.48039334 | 0.370950259 | 0.4256718 |
| Nr2f6 | 2.986104742 | 2.482938076 | 2.734521409 | Nr3c1 | 0.397732449 | 0.489861082 | 0.443796765 |
| Id1*<br>*Papaspyridonos, et al.*<br>2015 | 2.830149937 | 2.604014927 | 2.717082432 | Tfdp1 | 0.491835963 | 0.40071066 | 0.446273311 |
| Hbp1 | 3.075706126 | 2.264685563 | 2.670195844 | Bach2*<br>*Kurotaki, et al. 2018* | 0.414919227 | 0.490872432 | 0.45289583 |
| Hmbox1 | 2.483435981 | 2.721642917 | 2.602539449 |  |  |  |  |
| Mlxip | 2.456693653 | 2.637820801 | 2.547257227 |  |  |  |  |
| Mybl1 | 2.071956447 | 2.819078857 | 2.445517652 |  |  |  |  |
| Zfp691 | 2.261171987 | 2.15661673 | 2.208894358 |  |  |  |  |

Overall, the integrated analysis of gene expression and chromatin accessibility that we performed here not only suggested potential roles of known DC modulators in regulating changes in DC progenitors during infection but also highlighted 23 novel TFs as candidate regulators of DC development and/or function.

### GCs selectively suppressed pDC development and pDC-primed-signature expression in BM-Flt3L cultures

We posited that a biological pathway that is coordinately regulated by more than one TF among the aforementioned candidate TF regulators might be highly relevant in suppressing pDC generation and/or the bias toward cDC generation after viral infection. In this regard, two TFs that are known to regulate responses to glucocorticoids (GCs), Glucocorticoid Receptor (GR; encoded by *Nr3c1*) and Glucocorticoid Modulatory Element Binding Protein (GMEB1; encoded by *Gmeb1* gene) ^65^, exhibited discrete predicted differential activity between progenitors from infected versus uninfected mice (Fig. 4A, S2B and C, green dots), raising the possibility that GCs may regulate DC development.

While multiple circumstances including co-factor availability or post-transcriptional modification can change TF activity, several reports associated changes in TF accessibility to its binding sites with increased TF binding ^66; 67^ and altered target gene expression ^68^. Because Taiji takes the number of TF binding motifs within open chromatin regions as one of the determinants to rank TF activity, the greater the number of open chromatin regions containing the binding motif for a particular TF, the higher the predicted activity of such TF ^28^. To evaluate whether the altered GR and GMEB1 activity in progenitors from infected mice could result from changes in chromatin accessibility at these TFs’ binding sites, we collected all GR and GMEB1 binding motifs across the genomes and evaluated their ATAC-seq coverage in Lin^−^c-kit^int/lo^Flt3^+^ progenitors from uninfected vs. infected mice. Notably, the number of sequencing reads corresponding to the open chromatin regions at and near the sites that contain GR and GMEB1 binding motif was higher in progenitors from Cl13-infected mice compared to their counterparts from uninfected mice (Fig. 4B and C). Interestingly, the enhanced Gmeb1 (but not GR) accessibility to its binding sites in progenitors from infected mice was decreased in mice that received treatment with anti-IFNAR blocking Ab (Fig. 4D and E), which prevented the impairment of pDC development ^10^ and depletion of pre-pDCs after infection (Fig. 3F). These data indicated that both GR and GMEB1 binding sites across the genome exhibited higher degree of openness in progenitors from infected vs. uninfected mice and that the enhanced GMEB1 chromatin accessibility after infection was at least partially dependent on IFN-I signaling.

Given that Taiji predicted GR to exhibit lower activity in Lin^−^c-kit^int/lo^Flt3^+^ progenitors from infected mice (Fig. 4A) despite increased chromatin accessibility at GR binding sites (Fig. 4B), we investigated alternative factors that led to this Taiji prediction. Taiji utilizes the differential regulation of putative target genes as one of the determinants to assess the importance of a given TF^28^. Therefore, we first treated BM-Flt3L cultures derived from Cl13-infected mice with vehicle or 50nM of corticosterone for 3.5 days and investigated effects on gene expression in Pro-DCs, a DC committed progenitor with potential for all DC types that emerges in Flt3L culture^33^. RNA-seq analysis of Pro-DCs revealed significant changes in 1503 genes, among which 549 genes were upregulated and 954 genes were downregulated upon exposure to GCs (Table S3, Fig. 5A). Genes with significantly increased expression upon corticosterone treatment included previously reported GC-responsive genes such as Fn1 ^69^, Fabp4 ^70^, Tsc22d3 (a.k.a. GILZ) ^71^, IL6ra ^72^, and Fkbp5 ^73^ (Fig. 5A, labeled genes), confirming that treatment with corticosterone was effective. In addition, consistent with widely known anti-inflammatory effects of GCs ^74^, genes with decreased expression upon corticosterone treatment were significantly enriched for multiple pathways associated with immune activation (Fig. 5A and 5B), though notably we also observed a decrease in genes associated with suppression of immune responses. This could be representative of forms of negative regulation that are induced by immune activation (e.g. negative feedback), and therefore also inhibited when activation is suppressed.

**Figure 5.**
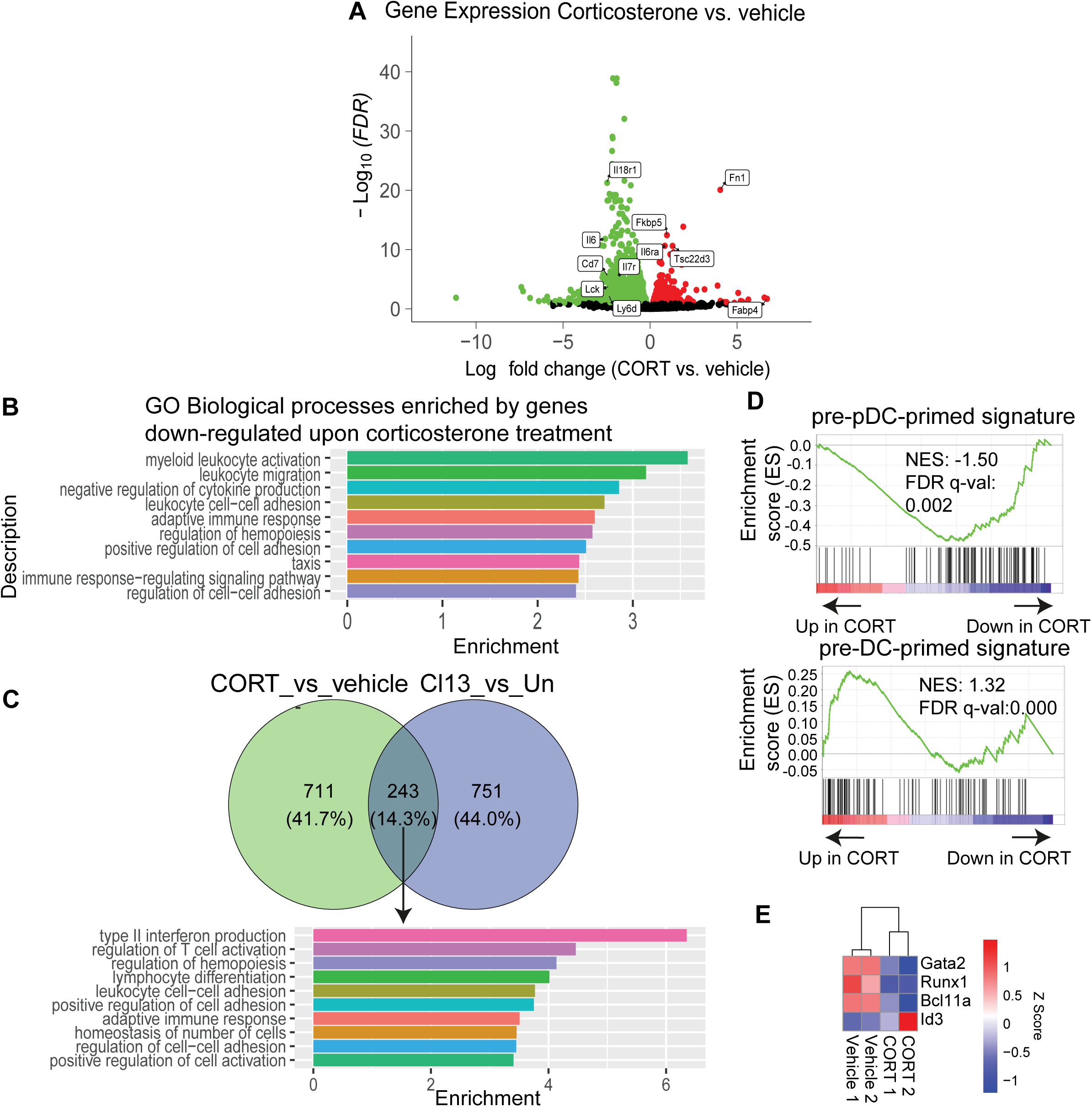
GC downregulated pre-pDC-primed signature. BM-Flt3L cultures from day 8 LCMV Cl13-infected mice were treated with vehicle or 50nM corticosterone (CORT) for 3.5 days. Pro-DCs were isolated from culture by flow cytometry for RNA-seq analyses. (A) Volcano plot of RNA-seq comparing CORT vs vehicle. Colored dots indicate DEGs (FDR < 0.1). Red and green dots indicate genes with higher or lower expression in pro-DCs from CORT-treated vs. vehicle-treated culture, respectively. (B) Top 10 Gene Ontology (GO) Biological Processes enriched by genes that were significantly downregulated in pro-DCs upon CORT treatment. (C) Venn diagram showing overlap between genes that were downregulated in pro-DCs upon CORT treatment and genes that were downregulated in Lin^−^c-kit^int/lo^Flt3^+^ progenitors after Cl13 infection based on RNA-seq performed in Fig. 1 (top) and Top 10 GO Biological Processes overrepresented by these genes (bottom). (D) GSEA and NES of DEGs for pre-pDC-primed and pre-DC-primed signature in Dress, et al. 2019. (E) Heatmaps of selected differentially expressed TFs. Data are representative of two independent repeats, each with 3-5 mice pooled per group.

We next sought to investigate the GC-regulated genes among the genes with significantly altered expression after infection. Approximately 5% of the genes upregulated in Lin^−^c-kit^int/lo^Flt3^+^ progenitors after Cl13 infection were also upregulated by the corticosterone treatment (Fig. S2D). Some of these genes were involved in the regulation of cytokine production, in line with the anti-inflammatory effects of GCs^74^ (Fig. S2D, right). Notably, ∼14% of the genes downregulated in Lin^−^c-kit^int/lo^Flt3^+^ progenitors after Cl13 infection were also downregulated by the corticosterone treatment (Fig. 5C). The excess of GC-downregulated over GC-upregulated genes among genes that are co-regulated in progenitors from Cl13-infected mice may explain the predicted lower activity of GR despite higher chromatin accessibility in GR-binding sites after Cl13 infection.

We observed that multiple pathways related to immune cell development and differentiation were overrepresented by the genes that were commonly reduced after Cl13 infection and GC treatment (Fig. 5C), raising the possibility that changes in pDC and cDC developmental capacity of Lin^−^c-kit^int/lo^Flt3^+^ progenitors are at least partially attributable to GC signaling.

Remarkably, GSEA analysis indicated that the pre-pDC-primed signature was reduced in corticosterone treatment, while the pre-DC-primed signature was increased (Fig. 5D, ^14^), suggesting a negative role for GCs in pDC development, and a possible positive role in cDC development. Some notable genes with altered expression upon corticosterone treatment included genes encoding critical TFs that have been previously shown to be required for pDC development at the progenitor stage, including GATA-2 ^75^, Runx1 ^76^, and Bcl11a ^45; 77^ (Fig. 5E). In contrast, Id3, which impairs pDC, but not cDC, development through antagonizing E-protein activity ^78^, showed significantly higher expression in GC-treated vs. vehicle control-treated pro-DCs (Fig. 5E). Collectively, these results indicate that GC treatment shifts pro-DC transcriptional state away from the pre-pDC primed state and toward the pre-DC primed state.

Of note, GCs have been previously shown to reduce the number of pDCs when added into BM-Flt3L cultures ^79^, but given that in these studies GCs were left in culture throughout the experiment, it is unclear whether this is the result of changes in DC progenitors or previously reported pro-apoptotic effects of GCs on differentiated pDCs ^80–82^. Given the significant reduction in pre-pDC-primed signature genes and the increase in pre-DC primed signature genes in pro-DCs treated with GCs (Fig. 5D), we next investigated the GC effects on DC progenitors’ generation of DCs, independently of GC effects on differentiated pDCs. For that, we first treated uninfected BM-Flt3L cultures with corticosterone from day 3.5 to day 4.5 p.c., a time when DC progenitors are enriched while most irrelevant cells have died and no mature DCs have yet developed ^33^. Strikingly, in response to this short corticosterone spike, pDC generation quantified at day 8 p.c. was profoundly and selectively compromised (Fig. 6A), while the relative development of cDC subsets (CD24^high^ cDC1, and CD11b^high^ cDC2) was slightly enhanced (Fig. 6B). Notably, when total yield was considered, this resulted in only a small increase in absolute number of cDC2 and no change in cDC1. Importantly, a similarly short corticosterone spike applied from day 6 to 7 p.c. – when most progenitors have already differentiated into DCs ^33^ – did not cause any significant effect on pDC frequency or numbers (Fig. S3A), suggesting that GC suppressed pDC generation by acting in DC progenitors rather than differentiated pDCs. In contrast, this same late treatment caused severe reduction in cDC1, and a relative increase in cDC2 among DCs (Fig. S3B), though notably, in terms of total yield, cDC2s were also reduced. These data suggest that the effects of corticosterone on DC progenitors and mature DCs are distinct, and in some cases oppositional. Specifically, a short spike of corticosterone treatment from day 3.5 p.c. to 4.5 p.c. reduced pDC development from progenitors, but treatment with corticosterone in mature pDCs from day 6 to 7 did not impact pDC numbers. In contrast, the same early treatment increased relative development of cDCs, but treatment of corticosterone from day 6 to 7 reduced both cDC1 and cDC2 with a more extreme effect on cDC1.

**Figure 6.**
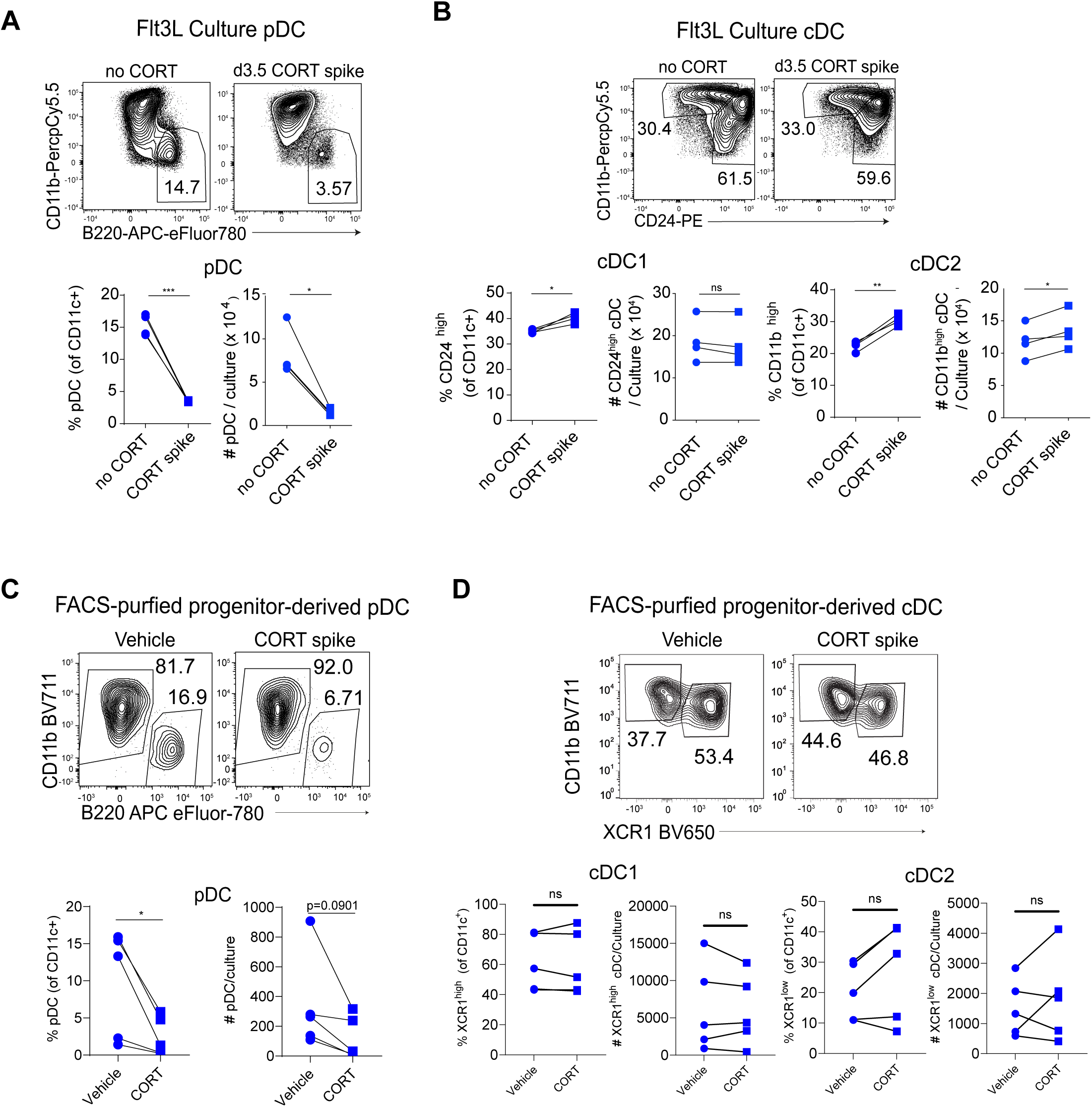
Early transient treatment of Lin^−^c-kit^int/lo^Flt3^+^ progenitors with GCs suppressed pDC development in BM Flt3L cultures. (A-B) BM cells from uninfected mice were cultured with Flt3L for 8 days. Cultures were given a spike of 1uM corticosterone or vehicle for 1 day from day 3.5 p.c. and media replaced thereafter. pDCs (A), or cDC (B), were analyzed for their frequency within DCs (CD11c^+^) (left) and absolute numbers (right) at day 8 p.c. (C-D) FACS-purified Lin^-^c-kit^int/lo^Flt3^+^ progenitors from uninfected mice were cultured with Flt3L for 8 days. Culture was given a spike of 1uM corticosterone for 1 day at Day 0 p.c. pDCs (C), cDC1s (XCR1^high^CD11b^low^) or cDC2s (XCR1^low^CD11b^high^) (D) were analyzed for their frequency within DCs (CD11c^+^) (left) or absolute numbers (right) at day 8 p.c. Data are representative of 2-3 experiments. *<0.05, **<0.01, ***<0.001, ****<0.0001. Statistical significance was determined by paired t-test (A-D).

To exclude the effects of GCs on other cells present at day 3.5 in BM-Flt3L culture that could be affected by GC and indirectly affect DC development, we cultured FACS-purified Lin^−^c-kit^int/lo^Flt3^+^ progenitors from uninfected mice with Flt3L for 8 days and treated these cells with a high dose of corticosterone from day 0 to day 1 p.c. Similar to what we observed in total BM-Flt3L cultures, the pDC number and frequency within differentiated DCs that developed from Lin^−^c-kit^int/lo^Flt3^+^ progenitors was decreased in response to an early and brief exposure to GCs (Fig. 6C), while the proportions of cDC subsets within DCs were unchanged in either proportion or absolute numbers (Fig. 6D). Altogether, these data show that GC selectively suppresses pDC development, and may bias progenitors toward cDC development in non-infectious conditions.

### Endogenous GCs selectively decreased pDC numbers during infection

Given the effect of exogenous GC on pDC development, we next sought to evaluate whether endogenous GC may also regulate pDCs *in vivo* and whether any difference was observed in infectious vs. noninfectious conditions. For that, we compared pDCs and their progenitors from uninfected vs. LCMV Cl13 infected adrenalectomized (ADX) mice (Fig. 7A), in which a major source of systemic GC is absent, using sham-operated mice as a control. Because most ADX mice died 5 days after LCMV Cl13 infection (Fig. 7B and ^83^), we were constrained to investigate pDC development in ADX vs. sham-operated mice at day 4 p.i. We observed a significant increase in the number of both BM and splenic pDCs from ADX vs. sham-operated infected (but not uninfected) mice (Fig. 7C and D). Notably, we observed no significant increase in pDC proliferation or decrease in apoptosis from ADX vs. sham-operated Cl13-infected mice (Fig. S4A-D), ruling out that the pDC numerical increase could have resulted from GC regulation of pDC proliferation or death rate. Intriguingly, the numbers of splenic cDC1s or cDC2s were not significantly altered in ADX uninfected or LCMV Cl13-infected mice (Fig. 7E), indicating a selective suppression of pDC but not cDC numbers by endogenous GC from infected mice at this timepoint.

**Figure 7.**
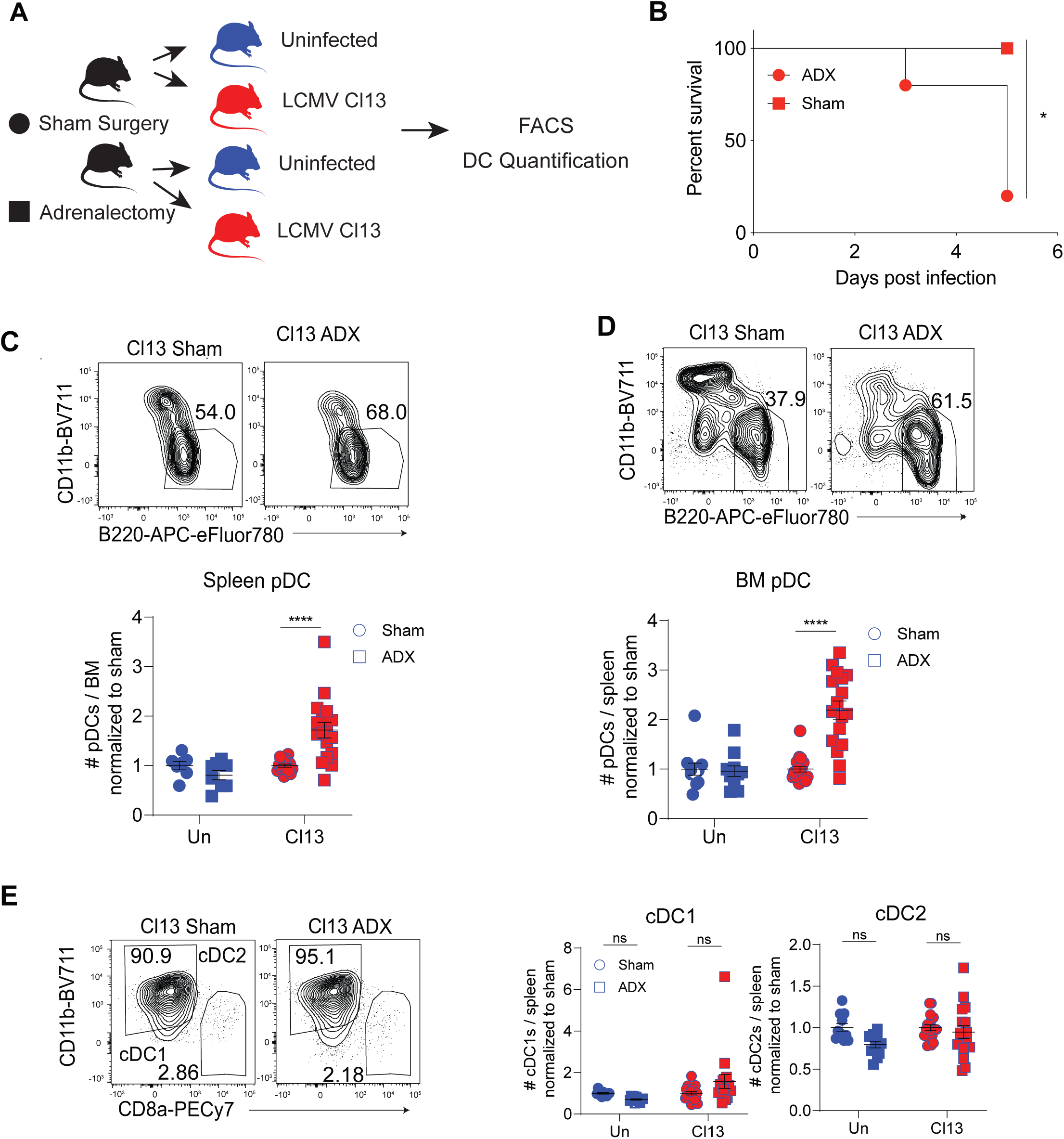
GCs suppressed pDC development and reduced the number of pDCs during *in vivo* LCMV infection. (A-E) Sham-operated (Sham) or adrenalectomized (ADX) C57BL6/J mice were left uninfected (Un) or infected with LCMV Cl13. (A) Diagram of adrenalectomized (ADX) mouse experiments. Sham surgery (Sham, circles) or ADX (squares) were either left uninfected (Un) or infected with LCMV Cl13. (B) Mouse survival following infection with LCMV Cl13. (C-E). At day 4 p.i., Spleen (C, E) or BM (D) was analyzed for number of pDCs (C, D) or cDCs (E). Graphs depict mean ± SEM, and symbols represent individual mice. Data are pooled from 2-4 experiments (C-E). *<0.05, ****<0.0001. Statistical significance was determined by Mantel-cox test (B), or 2-way ANOVA with Sidak’s multiple comparisons test (C-E).

Taken together, these data indicated that the numbers of splenic and BM pDCs, but not other DC subsets, were selectively compromised by GCs in the context of early LCMV Cl13 infection but not in uninfected conditions.

## Discussion

It is now well established that DCs have the capacity to adapt in response to infections ^3; 8; 84^. However, because mature DCs are short-lived, it is not surprising that these adaptations are accompanied with changes in the progenitor populations that have the potential to perpetually replenish DC compartments ^7; 10; 85^. Still, essential knowledge of what these adaptations are and how they impact mature DC populations is lacking. We have previously established that a specific, albeit heterogeneous, population of Lin^−^c-kit^int/lo^Flt3^+^ progenitors is adapted early after acute and chronic LCMV infections, becoming unable to generate pDCs ^10^. Through profiling of this progenitor population, we now identified convergent genomic changes early after acute and chronic infections, multiple mechanisms underlying the reduced number of progenitors with capacity to generate pDCs, as well as GCs as novel regulators of pDC development, connecting pDC progenitor adaptations to the HPA axis.

Integrated RNA and ATAC sequencing analysis of CD8^+^ T cells shows a clear divergence in acute and chronic infections ^25; 26^. Given the well-described differences in the environment created by these infections it might be expected the same would be true for other hematopoietic compartments. However, our analysis in Lin^−^c-kit^int/lo^Flt3^+^ progenitors with potential to generate DCs revealed that there was a convergent transcriptional and epigenetic state in these two distinct infection types. Although the transcriptional and epigenetic differences observed in these heterogeneous DC progenitors after infection may have partly resulted from changes in population composition, they are still contrasting to the diverging adaptation of T cells in acute and chronic infection and suggest the existence of a unitary default adaptation program in DC progenitors responding to viral infections. This program must be robust to override differences in viral titer ^86^, cytokine levels ^87–89^, T cell responses ^90; 91^ and systemic metabolites^92; 93^ that have been described to differ between these two infections at the time-point studied. On the other hand, the synchronicity of this program reinforces our previous observations that many of the pDC and progenitor adaptations previously described in chronic LCMV infection also emerge transiently in acute LCMV infection ^10^, making our study relevant for a multitude of infectious diseases regardless of pathogen clearance or persistence.

Having described the unique large-scale changes of DC progenitors, we then addressed the question of how these changes in Lin^−^c-kit^int/lo^Flt3^+^ progenitor transcriptome and epigenome connected to the impairment in pDC development that we previously described ^10^. We first focused on the well-established subsets of progenitors with potential to generate pDCs: CLP ^14; 15; 30–32^, CD115^-^ CDP ^34^, and CD115^+^ CDP ^17; 33^. Aided by our RNA-seq and ATAC-seq pathway analysis we identified infection-induced defects in CLP proliferation, which neatly explain our previously described decrease in their abundance^10^. We also detected increased proliferation, accompanied by increased apoptosis in the CD115^-^ CDP population after infection. As we have also previously described an overall decrease in this progenitor population ^10^, we speculate that increased apoptosis in this compartment must be the dominant effect. Concomitantly, we observed no changes in either proliferation or apoptosis in the CD115^+^ CDP compartment despite their decreased abundance^10^. Given these observations and the upregulation of Cxcr4, which is associated with BM retention^42^, in the DC progenitors from infected mice, we speculate that accelerated differentiation could be responsible for the numerical reduction of CD115^+^ CDP.

We then extended our investigation into the regulation of pre-DC and pre-pDC committed progenitors ^14; 15^. In line with a reduction in pre-pDC committed signature within Lin^−^c-kit^int/lo^Flt3^+^ progenitors, we observed a concurrent overall retraction in pre-pDC abundance. We also demonstrated that the depletion of pDC-committed pre-pDCs was dependent on IFN-I. This dovetails with our previous work indicating that the impaired pDC development and depletion of Lin^−^c-kit^int/lo^Flt3^+^ cells during infection occur in an IFN-I dependent manner ^10^. On the contrary, infection seemed to enhance the cDC-committed pre-DC signature and abundance, suggesting that this regulatory loop differentially affects both pDCs and cDC lineages. This is particularly intriguing as two previous studies ^7; 27^ described inhibition of cDC development during chronic LCMV infection. Our results would thereby suggest that such defect in cDC development is not due to a reduction in cDC-committed pre-DC, and could instead be a result of a depletion of earlier progenitors, inhibition of pre-DC differentiation into mature cDCs or compromised survival of the newly generated cDCs. Additionally, our results differ from a previous report suggesting a positive role of IFN-I signaling on pDC generation in uninfected conditions^94^. This discrepancy may be explained by differences in levels or timing of IFN-I signaling as well as potential cell-intrinsic versus extrinsic effects of these cytokines on DC progenitors.

Our study also leveraged our ATAC-seq and RNA-seq data via the Taiji algorithm to identify 34 TFs expected to have differential activity in DC progenitors during viral infection. The validity of this list is reinforced by previous studies using Taiji in other cell types ^49–51^ and the fact that 11 of these TFs have previously described roles in DC biology ^43; 52–64^. Importantly, by analyzing expression of the TFs’ putative target genes and accessibility of TFs at their binding sites, this analysis was able to identify candidate regulators of DC progenitor adaptation that are not themselves differentially expressed.

Taiji predicted infection-induced changes in activities of two TFs previously associated with responses to GCs, GR (encoded by *Nr3c1*) and GMEB1 (encoded by *Gmeb1*). Notably, we found that IFN-I signaling during *in vivo* infection enhances chromatin accessibility at Gmeb1 binding sites, providing a molecular explanation for both the enhanced Gmeb1 activity in Lin^−^c-kit^int/lo^Flt3^+^ progenitors from infected mice and the aforementioned role of IFN-I in the suppression of pDC development ^10^.

This subsequently prompted us to investigate a role for GCs in the regulation of DC development. Notably, this is conceptually in line with a large body of literature describing a cell-intrinsic apoptotic role for GC on mature pDC populations ^80–82^. However, this is the first evidence that GC may selectively modulate pDC numbers through the regulation of early lineage negative DC progenitors, affecting pDC development rather than differentiated pDCs. Furthermore, through our analysis of adrenalectomized mice we demonstrated that the GC suppression of pDCs is relevant *in vivo* as endogenous GC significantly reduced pDC numbers during and ongoing infection. It is important to emphasize that the levels of serum corticosterone in uninfected vs. infected mice were equivalent at the time point when adrenalectomized mice were analyzed (Fig. S4E). Thus, the selective effect of GC in pDC numbers after (but not before) infection may be explained by the infection-induced enhancement in DC-progenitors’ Gmeb1 activity, as Gmeb1 is known to increase sensitivity to GCs ^65^. Given that GC pro-apoptotic effects in mature pDCs can be counteracted by sustained Toll-like receptor (TLR) signaling^74; 82; 95; 96^, it is tempting to speculate that mature pDCs from Cl13-infected mice may experience similar protection through ongoing TLR stimulation by viral RNA. Enhancement in GMEB1 activity in progenitors could have evolved as a safeguard mechanism to keep the developmental output of pDCs at reduced levels after infection.

Notably, we also observed a slight but significant bias toward cDC development when GC treatment was performed *in vitro*, that interestingly did not extend to changes in mature cDC populations in mice depleted of GC *in vivo*. This discrepancy may be attributed to the limitation of using adrenalectomized mice, which are known to succumb early after Cl13 infection^83^, thus restricting our analysis to day 4 p.i.. Alternatively, it could reflect inherent differences between *in vitro* culture conditions and *in vivo* DC development. Future studies with animal models with GR- and/or GMEB1-deficiency restricted to DC progenitors, that may not have survival defects upon Cl13 infection, would allow for examination of GC effects on DC development at later time points p.i..

Manipulation of DC progenitors may offer not only the opportunity to provide a source of DCs that is self-perpetuating, but also access to the specialized processes provided by different DC subsets. Our study provides a characterization of the adaptations acquired by progenitors with potential to generate DCs in the context of a viral infection. We revealed a deep conservation of adaptation early after both a relatively mild as well as an aggressive fast-spreading infection context. This is a promising signal that manipulations of DC progenitors for therapeutic purposes may be robust across a spectrum of insults. We have also identified 23 novel targets whose impact is likely to be diverse across DC lineages. Ultimately, this work provides a foundation for both a deeper understanding of the logic behind DC progenitor adaptation to infection, as well as the molecular and organismal systems that regulate them, a necessary step toward the therapeutic use of DC progenitors in the treatment of human diseases.

## Materials and Methods

### Mice

C57BL6/J mice, CD45.1^+^ mice, sham-operated and ADX mice (7-8 weeks old) were purchased from The Jackson Laboratory. ADX mice were maintained on drinking water containing 1% NaCl per the manufacturer’s instructions. All mice were housed under specific-pathogen-free conditions at the University of California, San Diego (UCSD). Mice were bred and maintained in a closed breeding facility, and mouse handling and experiments conformed to the requirements of the National Institute of Health and the Institutional Animal Care and Use Guidelines of UCSD. Unless stated otherwise, experiments were initiated in mice (female and male) at 7-8 weeks of age.

### Virus strains

Mice were infected with 2 × 10^6^ plaque-forming units (pfu) of LCMV Armstrong (ARM) or LCMV clone 13 (Cl13) intravenously (i.v.) via tail vein. Viruses were propagated on BHK cells and quantified by plaque assay performed on Vero cells ^97^. Briefly, Vero cell monolayers were infected with 500μL of serially diluted viral stock and incubated for 60 minutes at 37°C in 5% CO_2_ with gentle shaking every 15 minutes. Agarose overlay was added to infected cells, and the cells were placed in an incubator for 6 days at 37°C in 5% CO_2_. Cells were fixed with formaldehyde and stained with crystal violet for 5 minutes at room temperature, and plaques were counted.

### Cell lines

HEK293T cells were cultured in DMEM containing 10% FBS, penicillin-streptomycin, and supplements of L-glutamine. 1.5×10^6^ cells were passaged every 3 days. Cells were washed in PBS pH 7.4 then lifted from culture flasks with 0.05% Trypsin-EDTA. Trypsin was deactivated by addition of serum containing media.

### Primary cell culture

BM DCs were generated at the concentration of 2 × 10^6^ cells/ml for 8 days in 5 ml of DC medium (RPMI-1640 supplemented with 10% (vol/vol) fetal bovine serum, L-glutamine, penicillin-streptomycin, and HEPES pH 7.2) supplemented with 100 ng/mL Flt3L (provided by CellDex Therapeutics) and 50 µM β-mercaptoethanol. Half of the medium was replaced after 5 days with medium with fresh Flt3L added. For corticosterone treatment, culture was replaced with fresh medium containing 50nM corticosterone every 2.5 days. For the short term high-dose corticosterone experiments (Fig. 6 and S3), culture was replaced with fresh medium containing 1µM corticosterone at indicated time points and was replaced again with fresh medium with no additional corticosterone after 24 hours.

### Mouse Cell Processing and Purification

Spleens were incubated with 1mg/mL collagenase D for 20 min at 37°C and passed through a 100µm strainer to achieve a single cell suspension. Bone marrow (BM) cells were prepared as single cell suspension by flushing the femurs and tibias of mice with DC medium. Lin^-^c-Kit^int/lo^Flt3^+^ progenitors were enriched by using an EasySep™ Mouse Streptavidin RapidSpheres™ Isolation Kit (Stemcell Technologies), biotin-conjugated anti-Thy1.2 (53-2.1), anti-B220 (RA3-6B2), anti-CD11b (M1/70), anti-CD4 (RM4-5), anti-CD19 (6D5) and anti-CD8 (536.7). Fractions unbound to streptavidin beads were stained with PI and FACS-purified using a BD ARIA II (BD) or ARIA Fusion (BD) for progenitors (c-Kit^int/lo^Flt3^+^) after exclusion of dead cells (stained with PI) and cells stained with lineage markers as follows: B cells (stained with CD19, B220), T cells (stained with Thy1.2, CD3, CD4, and CD8), NK cells (stained with NK1.1), erythroid cell (stained with Ter119), granulocyte (stained with Gr-1), monocyte (stained with CD11b), cDCs (stained with CD11c, CD11b and CD8) and pDCs (stained with CD11c, B220 and Gr-1). Note that the Gr-1 Ab used (RB6-8C5) binds Ly6C and Ly6G.

### Flow Cytometry

Single cell suspensions prepared from murine BM or spleens were stained with antibodies specific for the following markers: Thy 1.2 (53-2.1 or 30-H12), CD19 (eBio1D3 or 6D5), NK 1.1 (PK136), Gr-1 (RB6-8C5), CD11c (N418), CD11b (M1/70), B220 (RA3-6B2), BST2 (eBio927), CD45.2 (104), CD4 (RM4-5), CD8a (53-6.7), Ter119 (TER-119), CD3e (145-2c11), CD45.1 (A20), MHC class II/I-Ab (AF6-120.1), Ki67 (SolA15), c-kit (2B8), CD115 (AFS98), Flt3 (A2F10), CD127 (A7R34), Ly6d (49-H4), Siglech (eBio440c), Ly6G (1A8), CD138 (281-2), CD24 (M1/69), XCR1 (ZET), CD172α (P84), Clec9a (10B4), Ly6c (HK1.4). Unless stated otherwise, staining for surface markers was performed as previously described ^10^. Propidium Iodide (PI), Ghost dye (Tonbo Biosciences, San Diego, CA), or eBioscience™ Fixable Viability Dye eFluor™ 455UV was used to exclude dead cells. Cells were pre-incubated with CD16/CD32 Fc block (BD Pharmingen) for 10 minutes prior to surface staining. To measure *in vivo* BrdU incorporation in progenitors, mice were injected with 2mg BrdU (Sigma Aldrich). 16 hours later, BM cells were harvested and stained using the BrdU Flow kit (BD Biosciences) following the manufacturer’s instructions. Staining for Ki67 (SolA15) was performed using the Foxp3 / Transcription Factor Staining kit (Thermo Fisher Scientific) following the manufacturer’s instructions. Active caspase-3 staining protocol was performed following the manufacturer’s instructions of the PE active caspase-3 apoptosis kit (BD Biosciences). Cells were acquired with a LSRII flow cytometer (BD Biosciences) or ZE5 analyzer (Biorad) and data were analyzed using FlowJo software 9.9.6 or 10 (Treestar, Inc.).

### RNA Sequencing

For RNA-seq studies in Lin^-^c-Kit^int/lo^Flt3^+^ progenitors, cells were collected from three independent experiments in which BM cells from 3-5 mice per group were pooled. For RNA-seq studies in pro-DCs, cells were collected from two independent experiments in which BM Flt3L cultures from 3-5 mice per group were pooled. Cells from each independent experiment were kept at −80C in RLT buffer from RNeasy Micro kits (Qiagen) containing β-mercaptoethanol until all the independent experiments were completed. Library preparation for all the independent experimental repeats was performed simultaneously. Total RNA was prepared from 2 × 10^4^ to 1 × 10^5^ cells, and subsequently processed by the UCSD IGM Genomics center to generate an mRNA-seq library using a TruSeq RNA Library Prep Kit v2 (Illumina). The library for Lin^-^c-Kit^int/lo^Flt3^+^ progenitors was sequenced using Hiseq 4000 for single-end 100bp sequencing. The library for pro-DCs treated with vehicle or 50nM corticosterone was sequenced using NOVA-seq for paired-end 50bp sequencing. Raw sequencing data were processed with CASAVA 1.8.2 (Illumina) to generate FastQ files. Raw sequencing data were aligned to mouse genome mm10 (Genome Reference Consortium GRCm38) by using Spliced Transcripts Alignment to a Reference (STAR) aligner ^98^ with the following parameters: “--outFilterType BySJout --outFilterMultimapNmax 20 --alignSJoverhangMin 8 --alignSJDBoverhangMin 1 -- outFilterMismatchNmax 999 --outFilterMismatchNoverReadLmax 0.04 –alignIntronMin 20 --alignIntronMax 1000000 --alignMatesGapMax 1000000”.

Transcript data from STAR were subsequently analyzed using RSEM ^99^ for gene and transcript level quantification. Downstream analyses and heatmaps were performed with R 3.5.3 ^100^ and Bioconductor 3.8 ^101^. For differential gene expression analysis, we filtered out genes with low expression by setting a threshold for minimum expression level such that the sum of Transcript per Million (TPM) for 9 samples (3 conditions, 3 biological replicates for each condition) is greater than or equal to 9. Differentially expressed genes (DEGs) were determined by DEseq2 using a 2-fold change with false discovery rate (FDR) < 0.05 (for Lin^-^c-Kit^int/lo^Flt3^+^ progenitors) or FDR < 0.1 (for pro-DCs) as the threshold ^102^. Pathway analyses were done with WebGestalt ^40^ using genes with altered expression with FDR < 0.05 as threshold and gene set enrichment analysis through GSEA ^103^ software using DEGs.

### ATAC Sequencing

For ATAC-seq studies in Lin^-^c-Kit^int/lo^Flt3^+^ progenitors from uninfected or LCMV-infected mice, cells were collected from three independent experiments in which BM cells from 3-5 mice per group were pooled. For ATAC-seq studies in Lin^-^c-Kit^int/lo^Flt3^+^ progenitors from Cl13-infected mice treated with anti-IFNAR antibody or isotype control, cells were collected from one experiment with 10 mice per group and BM cells were pooled from 3-4 mice per group resulting in three samples per group. ATAC-seq was performed according to a published protocol ^39^ with minor modification. Briefly, cells were sorted (2.5 × 10^4^), resuspended in lysis buffer and centrifuged at 600g for 30 min at 4°C. Nuclei pellet was resuspended into transposition reaction mixture containing Tn5 transposase from Nextera DNA Sample Prep Kit (Illumina) and incubated for 30 min with gentle shaking at 37°C. Following purification of transposed DNA, cells from each independent experiment were kept at −20°C until all the independent repeats were completed. Library preparation for all the independent experimental repeats was performed simultaneously. We first amplified the full libraries for 5 cycles and the additional number of cycles needed for the amplification of the remaining 45 μL reaction was determined as published in the original protocol ^39^. The libraries were purified using a Qiagen QIAquick PCR purification kit in 20 μL. Libraries were amplified for a total of 10–12 cycles. The library was pooled and size-selected to range between 100bp and 600bp by using PippinHT (Sage Sciences). Finally, the libraries were sequenced using Hiseq 4000 for single-end 100bp sequencing. The sequencing reads were aligned to a mouse genome (mm10; Genome Reference Consortium GRCm38) using BWA ^104^ with the following parameters: “bwa mem -M -k 32”. Low quality and unmapped reads were removed using samtools^105^ with the following parameters: “-F 0×70c -q 30”. After the removal of duplicated reads, peaks for each individual replicate were identified using MACS2 ^106^ with q value cutoff of 0.01. Downstream analyses and heatmaps were generated with R 3.5.3 ^100^ and Bioconductor 3.8^101^. The differential chromatin accessibility was determined by Diffbind ^107^ using DESeq2-based statistical method and FDR < 0.05 as the threshold. Pathway analyses were done with Genomic Regions Enrichment of Annotations Tool (GREAT) ^41^.

### Transcription Regulatory Network Analysis and TF Ranking

The network analysis was carried out using the Taiji software ^28^. Briefly, TF binding sites were identified within open chromatin regions and assigned to the nearest genes. Next, the gene expression levels were used to assign weights to the edges and nodes in the network. Lastly, personalized PageRank algorithm was used to measure the global influence of TFs in the network, defined by the PageRank scores. Let **s** be the vector containing node weights and **W** be the edge weight matrix. The personalized PageRank score vector **v** was calculated by solving a system of linear equations **v** = (1 − *d*)**s** + *d***Wv**, where *d* is the damping factor (default to 0.85), s is the vector containing node weights, and W is the edge weight matrix ^28^. Key TFs in Table 1 were filtered using three criteria: 1) Average rank of all conditions is greater than 0.0001. 2) The fold change between progenitors from LCMV-infected mice and uninfected mice is over 2 or below 0.5 in both LCMV ARM vs. uninfected and LCMV Cl13 vs. uninfected comparisons. 3) The expression level of TF passes the minimum expression threshold (TPM > 1).

### Progenitor Transfer Assay

For transfer of Lin^-^c-Kit^int/lo^Flt3^+^ progenitors, 1.5-4 × 10^4^ Lin^-^c-Kit^int/lo^Flt3^+^ cells were FACS-purified as indicated above from the BM of CD45.2^+^ uninfected or Cl13-infected donor mice at day 8 p.i., and injected i.v. into CD45.1^+^ non-irradiated uninfected or infection-matched recipient mice, respectively. Mice were sacrificed at day 8 post transfer, and spleen cells were analyzed by flow cytometry after splenocytes were enriched for donor-derived cells by using EasySep™ Mouse Streptavidin RapidSpheres™ Isolation Kit (Stemcell Technologies) and biotin-conjugated anti-CD45.1 (A20).

### Corticosterone, Flt3L, and LIF Measurements

Corticosterone in media used for BM-Flt3L culture and culture supernatants was measured by Corticosterone Competitive ELISA Kit (Thermo Fisher Scientific), following the manufacturer’s instructions. Flt3L in serum from uninfected, ARM-infected, and Cl13-infected mice was measured by Mouse/Rat Flt-3 Ligand/FLT3L Quantikine ELISA Kit (R&D Systems), following the manufacturer’s instructions. LIF in serum from uninfected, ARM-infected, and Cl13-infected mice was measured by LEGEND MAX™ Mouse LIF ELISA Kit (BioLegend).

### *In vivo* IFNAR Blockade

C57BL6/J mice were intraperitoneally (i.p.) injected with neutralizing antibody (nAb, 500μg/mouse) against IFNAR (clone MAR1-5A3; BioXCell) or Anti-Mouse IgG1 Isotype Control (clone MOPC-21; BioXCell) on days −1 and 0 post-LCMV Cl13-infection, as done previously ^108^. Mice were then treated with 250μg/mouse anti-IFNAR nAb or isotype control i.p. on days 2, 4, and 6 post-LCMV Cl13 infection as previously described_108._

### Quantification and Statistical Analysis

All statistical parameters are described in figure legends. Student’s t tests (two-tailed, unpaired, or, where indicated, paired), one-way Anova, two-way Anova, Mantel-cox test and multiple comparisons were performed using GraphPad Prism 8.0 (GraphPad). Significance was defined as p ≤ 0.05. In all figures, error bars indicate S.E.M.

## Acknowledgments

We would like to thank all members of Zuniga lab for helpful discussion and feedback. Sample quality control, size selection, and sequencing for RNA-seq and ATAC-seq, RNA-seq library generation, and size selection for ATAC-seq were conducted at the IGM Genomics Center, University of California, San Diego, La Jolla, CA. This study was supported by a research grant from San Diego Center for Precision Immunotherapy and National Institute of Health grants AI132122, AI145314, AI081923 and AI113923.

