## Supplemental Figures for "Genomic Analysis of Progenitors in Viral Infection Implicates Glucocorticoids as Suppressors of Plasmacytoid Dendritic Cell Generation"

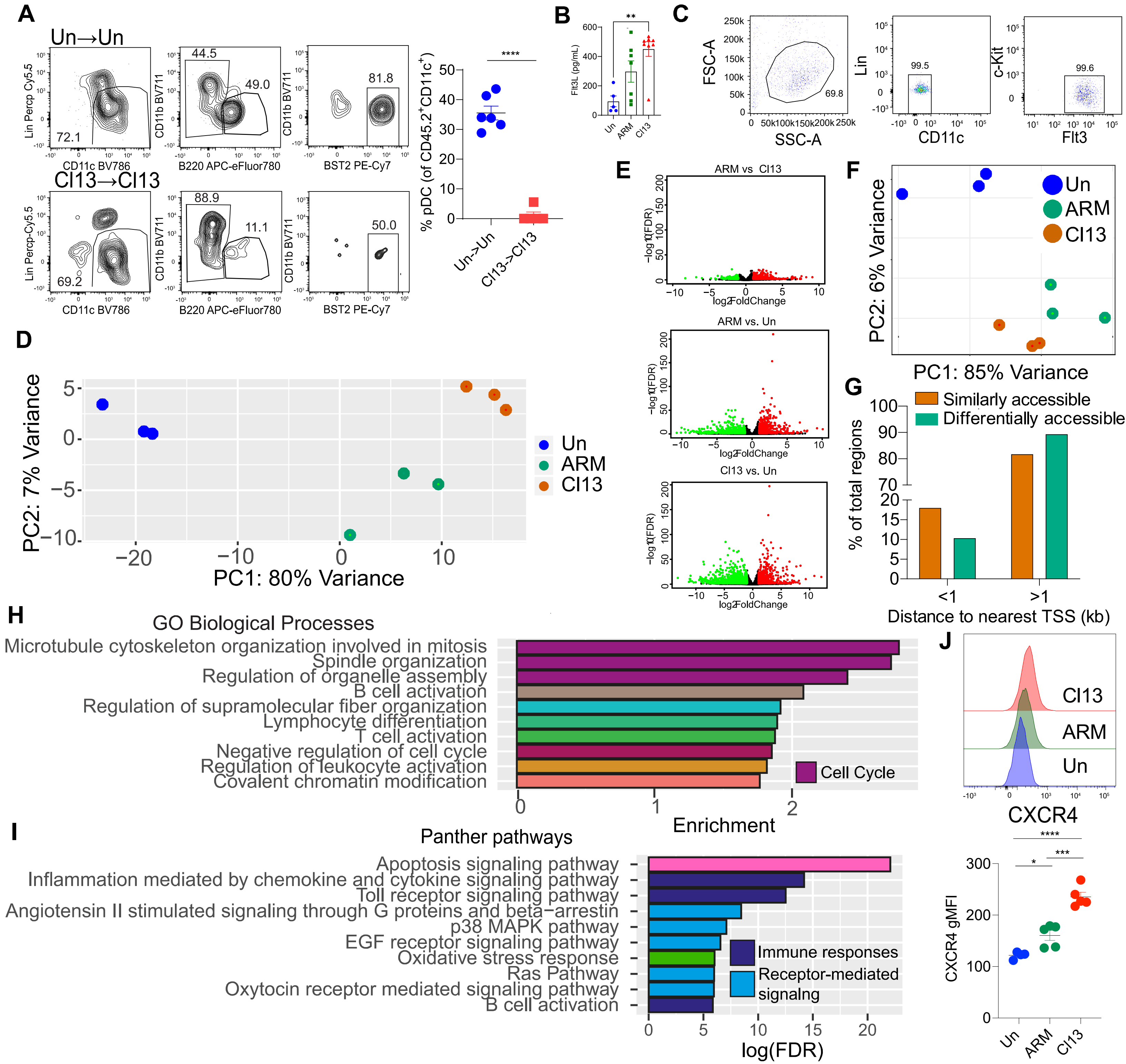


**Figure S1. Acute and chronic LCMV infections induced closely related transcriptomes and chromatin landscapes in DC progenitors.** C57BL6/J mice were infected with LCMV ARM or Cl13 or left uninfected (Un) for 8 days. (A) FACS-purified Lin$^-$c-kit$^{int/lo}$Flt3$^+$ progenitors (1-4 x $10^4$ cells) from CD45.2$^+$ uninfected or Cl13-infected donor mice were injected intravenously into CD45.1$^+$ non-irradiated uninfected or infection-matched recipients, respectively. Mice were sacrificed on day 8 after transplantation (day 16 p.i.), and spleen cells were analyzed by flow cytometry. Gating strategy and representative flow cytometry plots for spleen pDCs within donor- derived cells are shown. (B) Flt3L levels in serum were measured by ELISA. (C-I) Lin$^-$c-kit$^{int/lo}$Flt3$^+$ progenitors were isolated by flow cytometry for RNA-seq (D-E, H) and ATAC-seq analyses (F, G, and I). (C) Purity of Lin$^-$c-kit$^{int/lo}$Flt3$^+$ progenitors. (D) Principal component analysis (PCA) of RNA-seq profiles using the top 500 variable genes. Replicates are represented as separate data points and color- coded by presence/type of infection. (E) Volcano plots of RNA-seq profiles for the indicated group comparisons. Colored dots indicate DEGs. Red dots indicate genes with fold change > 2 and green dots indicate genes with fold change < -2. (F) PCA of ATAC-seq profiles. Replicates are represented as separate data points and color-coded by presence/type of infection. (G) Distance to nearest transcription start sites (TSS) for all accessible regions that were similarly (orange) and differentially (green) accessible in progenitors from ARM-infected or Cl13-infected mice compared to those from uninfected mice. (H) Top 10 Gene Ontology (GO) Biological Processes enriched by genes that were downregulated (FDR < 0.05) in progenitors during both ARM and Cl13 infections. (I) Panther pathways enriched in regions that become more accessible in progenitors during both ARM and Cl13 infections (FDR < 0.05). (J) CXCR4 expression in progenitors analyzed by flow cytometry. ARM vs. Un comparison reached statistical significance in 2 out of 3 experiments. Data are pooled from 2 experiments (B) or representative of 2 (A) or 3 independent experiments (C-J). *<0.05, **<0.01, ***<0.001, ****<0.0001. Statistical significance was determined by student's t-test (A) or one-way ANOVA with Tukey's multiple comparisons test (B, J).

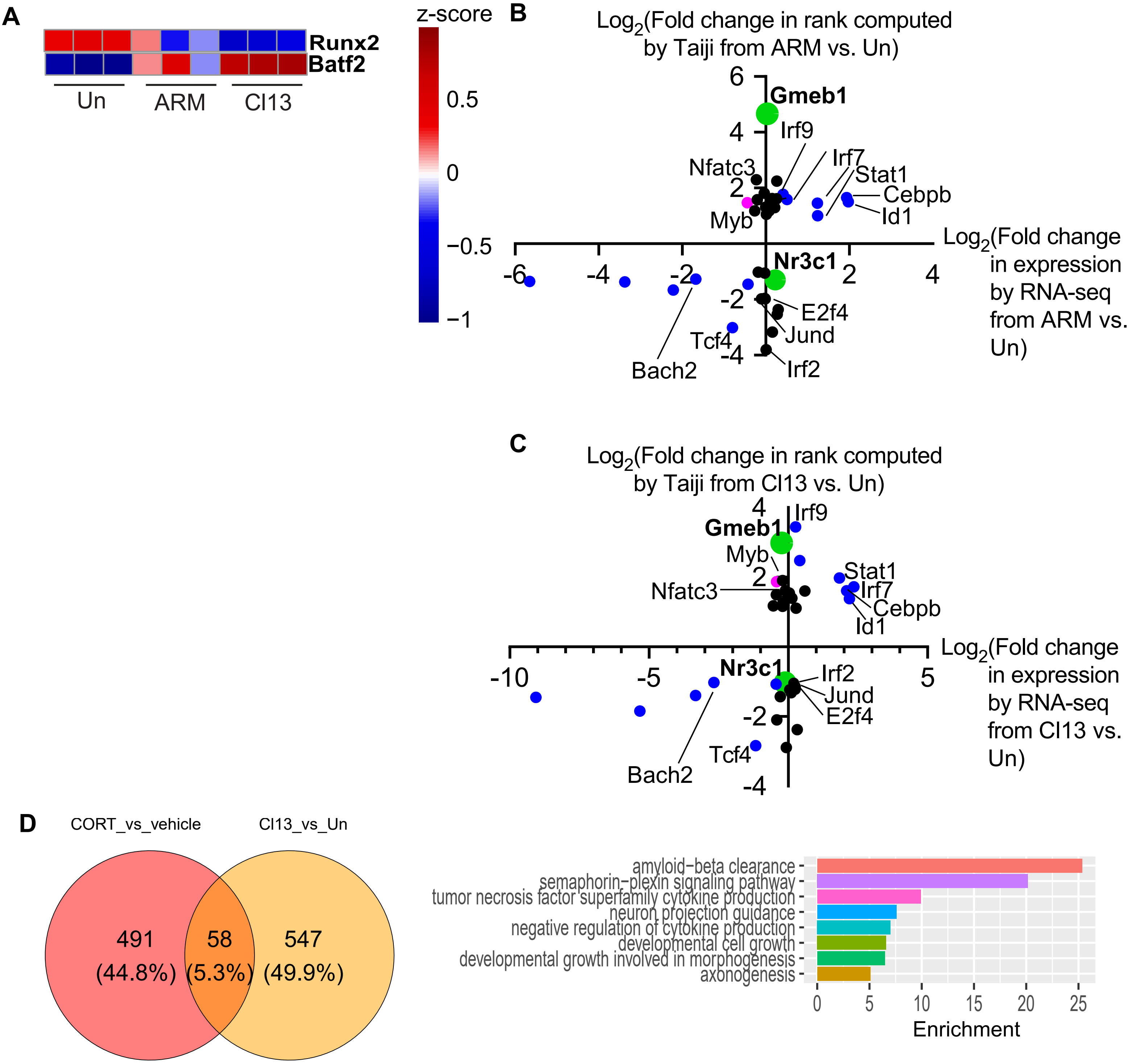


**Figure S2. Infection altered expression and predicted activity of multiple TFs in DC progenitors from infected mice and GC downregulated pre-pDC-primed signature.** (A) Heatmaps of selected differentially expressed (DE) TFs with known roles in DC development that were significantly differentially expressed in progenitors from both ARM- and Cl13-infected mice compared to their counterparts from uninfected mice are shown. (B and C) Lin$^-$c-kit$^{int/lo}$Flt3$^+$ progenitors were isolated from BM by flow cytometry for RNA-seq and ATAC-seq analyses, which were subsequently used in Taiji analysis. Scatter plot shows correlation between fold change in expression quantified by RNA-seq and fold change in rank computed by Taiji of TFs predicted by Taiji to have altered activities in progenitors from both ARM- and Cl13-infected vs. uninfected mice. Plots for comparison of progenitors from ARM-infected (B) or Cl13-infected (C) vs. uninfected mice are shown. Fold changes in ARM vs. uninfected and Cl13 vs. uninfected comparisons was log2-transformed to generate the plot. Differential expression was defined by DESeq2 with the threshold of FDR < 0.05. (D) Venn diagram showing overlap between genes that were upregulated in pro-DCs upon CORT treatment and genes that were upregulated in Lin$^-$c-kit$^{int/lo}$Flt3$^+$ progenitors after Cl13 infection based on RNA-seq performed in Fig. 1 (top) and top 10 GO Biological Processes overrepresented by these genes (bottom). Data are from 3 (A-C) or 2 (D-E) independent repeats, each with 3-5 mice pooled per group.

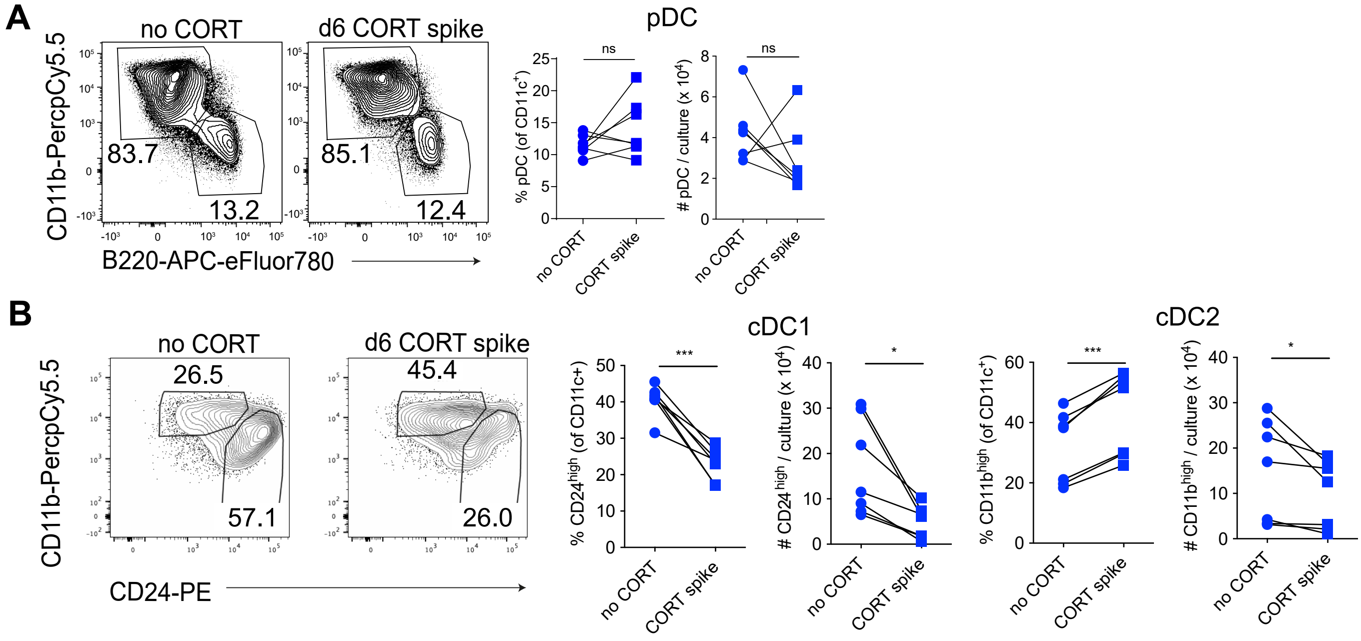


**Figure S3. GC spike after DC differentiation did not change pDC recovery from Flt3L culture.** BM cells from uninfected mice were cultured with Flt3L for 8 days and given a spike of 1uM corticosterone or vehicle for 1 day from day 5 to 6 p.c. and were replaced with fresh medium. pDCs (A) and CD24$^{high}$ and CD11b$^{high}$ cDCs (B) were analyzed for their frequency within DCs (left) and absolute numbers in culture at day 8 p.c. (right). Representative flow cytometry plots are shown.

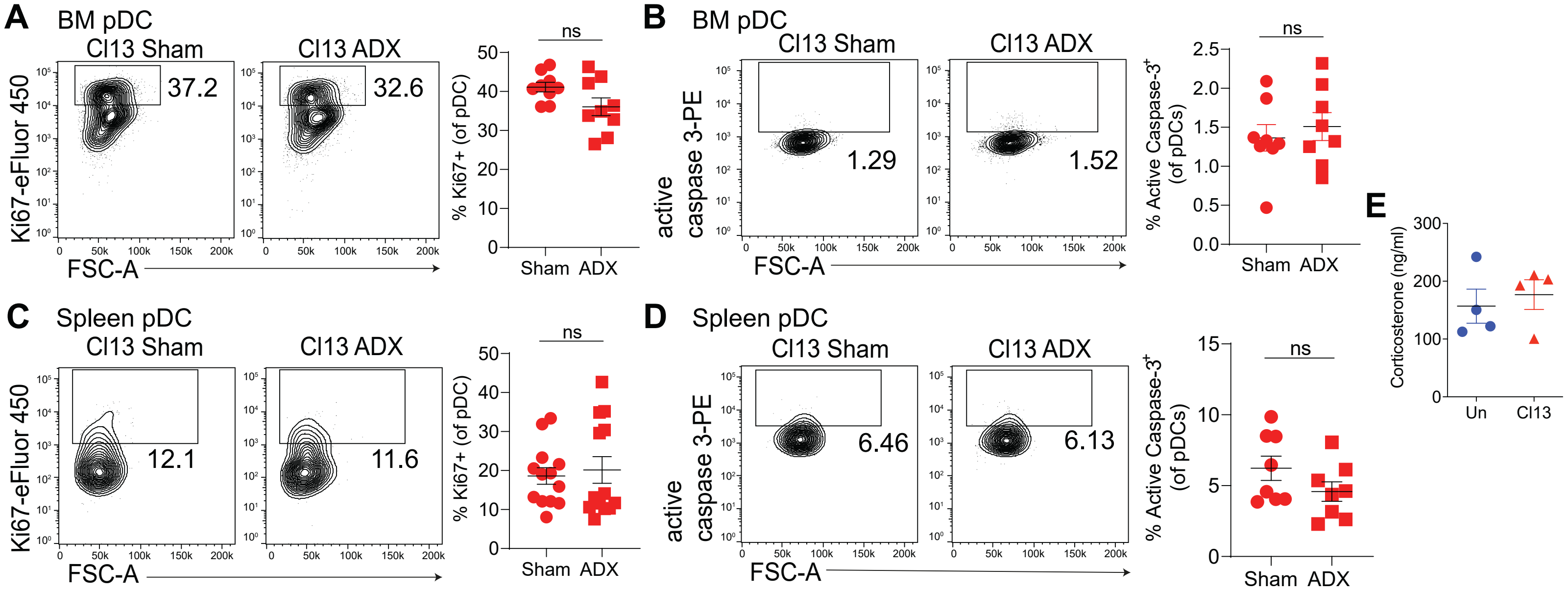


**Figure S4. Adrenalectomized mice did not show differences in pDC proliferation or apoptosis after LCMV infection.** (A-D) Sham-operated (Sham) or adrenalectomized (ADX) C57BL6/J mice were infected with LCMV Cl13. At day 4 p.i., BM (A, B) and spleen (C, D) were analyzed for pDC Ki67 expression (A, C) and Caspase 3 activity (B, D). (E) C57BL6/J mice were infected with LCMV Cl13 or left uninfected (Un) for 4 days. Corticosterone levels in serum were measured by ELISA. Data are pooled from 2-4 independent experiments (A-D) or representative of 2 experiments (E). ns, not significant. Statistical significance was determined by unpaired, two-tailed t-test.
