## Supplemental Table 1 for "Genomic Analysis of Progenitors in Viral Infection Implicates Glucocorticoids as Suppressors of Plasmacytoid Dendritic Cell Generation"

Supplementary Table 1. List of genes that are differentially expressed in Lin$^{-}$c-kit$^{int/lo}$Flt3$^{+}$ progenitors from day 8 p.i. ARM-infected vs. uninfected mice by RNA-seq.

| **Symbol** | **log2FoldChange (ARM vs. Un)** | **FDR** |
|---|---|---|
| Adgrb1 | 10.07782 | 1.23E-10 |
| Amotl2 | 9.263451 | 1.41E-12 |
| Dnmt3l | 7.868989839 | 1.02E-07 |
| Klra2 | 7.775527 | 4.60E-06 |
| Cd200r4 | 7.454751 | 1.90E-07 |
| Agmo | 6.995119 | 0.000934 |
| Ccl24 | 6.973663034 | 7.38E-05 |
| Gm4951 | 6.812208 | 8.54E-25 |
| Ppef2 | 6.588256 | 0.003537 |
| Ablim2 | 6.533229 | 0.01303 |
| Hsd17b14 | 6.429614 | 6.71E-05 |
| Fignl2 | 6.373503 | 0.025648 |
| Vnn3 | 5.914098138 | 0.01805 |
| 9430069I0 | 5.900618 | 0.001925 |
| H2-Bl | 5.894765 | 3.70E-06 |
| Mak | 5.744286179 | 0.024012 |
| 1520401A0 | 5.741605 | 0.029207 |
| Trim69 | 5.729114 | 0.038215 |
| Efemp2 | 5.604816 | 0.027233 |
| Ear2 | 5.538079 | 0.01093 |
| Spon1 | 5.340609 | 1.79E-17 |
| Insyn2b | 5.232163 | 0.01065 |
| Spocd1 | 5.216781 | 0.012171 |
| Shisa3 | 5.183465 | 0.010804 |
| Chst1 | 5.089965 | 0.002996 |
| Ace | 5.080728993 | 0.002627 |
| F830016B0 | 5.079698 | 1.30E-12 |
| C4b | 5.068912 | 8.25E-09 |
| Cbr2 | 5.053093 | 0.002122 |
| Serpi3f | 5.028479 | 5.07E-10 |
| Skida1 | 5.017655 | 0.037602 |
| Sectm1a | 5.016421 | 1.38E-06 |
| Art2b | 4.985318 | 0.046559 |
| Nexn | 4.955934 | 0.011057 |
| Fam83a | 4.92515 | 0.000175 |
| Batf2 | 4.884463 | 7.48E-20 |
| Clec4a4 | 4.85876 | 6.73E-06 |
| Glb1l2 | 4.854696 | 0.000111 |
| Spink2 | 4.703579 | 0.000615 |
| Clec4a2 | 4.700564 | 1.33E-13 |
| Ly6a | 4.66398 | 5.11E-24 |
| Penk | 4.650019 | 0.03064 |
| Clec4a3 | 4.559465 | 2.80E-17 |

| | | |
|---|---|---|
| Arhgef37 | 4.547767 | 4.01E-13 |
| Ccl19 | 4.486544 | 6.82E-10 |
| Rhou | 4.430932 | 1.96E-11 |
| Gm35549 | 4.365549 | 0.040084 |
| Inpp5j | 4.354964 | 0.033933 |
| Hoga1 | 4.273072 | 0.011257 |
| Prkn | 4.212476 | 0.024092 |
| Cfb | 4.173132 | 0.004316 |
| Mmp25 | 4.130059 | 0.000447 |
| Gpnmb | 4.127084 | 9.00E-11 |
| Porcn | 4.067862 | 0.000548 |
| Slfn1 | 4.056875 | 2.14E-06 |
| Scarf1 | 4.029432 | 4.40E-44 |
| Iigp1 | 4.012515 | 2.53E-16 |
| Aplnr | 4.011717 | 0.002317 |
| Htr2b | 4.001709 | 0.007292 |
| Gbp2 | 3.973826 | 1.50E-47 |
| Tchh | 3.87412 | 2.67E-15 |
| Ly6i | 3.870208 | 1.31E-18 |
| LOC11548[illegible] | 3.766121 | 0.031598 |
| Slc13a3 | 3.752926395 | 2.36E-05 |
| Hpgd | 3.746869 | 4.19E-18 |
| Lin28a | 3.630162 | 0.016036 |
| Apoc2 | 3.603412968 | 1.20E-06 |
| Pilrb2 | 3.585526 | 0.009698 |
| Ifi213 | 3.570367 | 2.64E-07 |
| Rse2a | 3.545238 | 0.026495 |
| Gm44805 | 3.545003 | 1.48E-05 |
| Nod2 | 3.532572 | 2.78E-12 |
| Gda | 3.457151 | 3.63E-35 |
| Abca8b | 3.388991181 | 0.039001 |
| Plekhg4 | 3.376504245 | 0.000395 |
| Il27 | 3.372261 | 0.000159 |
| Serpi3g | 3.350578 | 3.21E-11 |
| Trim30c | 3.336757 | 0.009349 |
| Cd74 | 3.321595 | 5.06E-35 |
| H2-Eb1 | 3.304971 | 4.18E-19 |
| Cp | 3.273167595 | 2.92E-23 |
| Cxcl9 | 3.23299 | 0.035549 |
| H2-Q2 | 3.220607 | 0.034979 |
| H2-Aa | 3.215585 | 3.43E-24 |
| Ifi204 | 3.204116 | 1.16E-58 |
| Mxra7 | 3.193058908 | 0.003001 |
| Saa3 | 3.173812 | 4.75E-05 |
| Msr1 | 3.171607 | 0.005433 |
| Ly6a2 | 3.103527 | 4.65E-38 |
| Serpi3i | 3.091629 | 3.50E-11 |

| | | |
|---|---|---|
| Cux2 | 3.018685 | 0.015324 |
| H2-Q7 | 3.01191 | 7.94E-211 |
| Gm13710 | 2.997078 | 0.021524 |
| Fcgr1 | 2.996940227 | 6.11E-18 |
| Bvht | 2.983925 | 0.023734 |
| Zbp1 | 2.982938 | 1.51E-17 |
| Myof | 2.97485 | 6.49E-34 |
| Acpp | 2.966639 | 0.000135 |
| Tlr13 | 2.946324 | 6.81E-06 |
| Lyz2 | 2.935735 | 9.74E-13 |
| Rspo1 | 2.926015 | 3.96E-05 |
| Adam3 | 2.905092 | 0.007245 |
| Rem1 | 2.885035387 | 9.21E-07 |
| H2-Q6 | 2.861402 | 5.27E-154 |
| Prdm1 | 2.848334 | 0.015881 |
| H2-Q5 | 2.808707 | 2.81E-76 |
| Mag | 2.797939 | 4.28E-05 |
| Ptges | 2.782076 | 0.033341 |
| Lcn2 | 2.770414 | 0.032104 |
| Lyz1 | 2.748171 | 7.05E-16 |
| Clec4b1 | 2.746469 | 1.83E-06 |
| Gm10825 | 2.743522 | 0.038192 |
| Gpx3 | 2.734176916 | 2.19E-24 |
| Cd40 | 2.680648261 | 8.11E-19 |
| Nxpe4 | 2.672514 | 1.33E-15 |
| Ptprr | 2.671680567 | 1.43E-06 |
| Slamf8 | 2.671448 | 1.19E-11 |
| Ciita | 2.653331 | 0.001321 |
| Mcemp1 | 2.651007286 | 1.25E-13 |
| Clec4a1 | 2.649292 | 0.006828 |
| AB124611 | 2.633285 | 6.26E-63 |
| Plekhg1 | 2.627403 | 1.06E-11 |
| C3 | 2.615382 | 1.08E-40 |
| Filip1 | 2.615256 | 0.00058 |
| Tlr4 | 2.613954 | 2.20E-05 |
| Sult1a1 | 2.613887 | 0.00065 |
| Glt1d1 | 2.612391 | 6.48E-06 |
| H2-Ab1 | 2.60286 | 6.03E-22 |
| Trpm2 | 2.602813931 | 1.57E-33 |
| Ifitm6 | 2.593799 | 3.78E-37 |
| Xdh | 2.579626 | 1.53E-40 |
| Gm15972 | 2.573967 | 0.013526 |
| Neurl1a | 2.573822962 | 0.022783 |
| Tnfrsf8 | 2.569257 | 0.00121 |
| Pgap1 | 2.561024 | 0.00029 |
| Mfsd7a | 2.558437 | 2.81E-11 |
| F10 | 2.525141 | 2.46E-49 |

| | | |
|---|---|---|
| Hal | 2.521915103 | 4.73E-06 |
| Hopx | 2.511882 | 1.10E-16 |
| Gpr85 | 2.480739 | 0.014778 |
| Fcgr3 | 2.461734 | 5.57E-20 |
| Parp12 | 2.450318 | 4.18E-17 |
| Gm12185 | 2.446588 | 1.36E-08 |
| Gm9733 | 2.445386 | 0.003328 |
| Tmem178 | 2.444929 | 8.30E-40 |
| Fn1 | 2.443507 | 8.16E-05 |
| F7 | 2.417207 | 8.20E-06 |
| Shtn1 | 2.415604 | 2.91E-16 |
| Abca13 | 2.410811508 | 0.000507 |
| Osgin1 | 2.410525 | 0.000609 |
| Mgst1 | 2.408810845 | 2.13E-26 |
| 1600010M | 2.398277 | 0.000112 |
| Nlrp1c-ps | 2.396076 | 3.56E-05 |
| Gm4841 | 2.395397 | 0.000675 |
| Ms4a4c | 2.372636 | 1.18E-51 |
| Cybb | 2.371391876 | 5.62E-33 |
| Gpr15 | 2.35917 | 0.000783 |
| Oas1g | 2.358883 | 6.55E-17 |
| Adgre5 | 2.355596953 | 6.55E-17 |
| Cd177 | 2.350402 | 7.34E-06 |
| Phf11d | 2.34096 | 2.35E-06 |
| Mapk13 | 2.336264124 | 4.35E-06 |
| BB031773 | 2.285696 | 1.48E-10 |
| Gstm3 | 2.283272771 | 9.57E-06 |
| Bmx | 2.282451 | 2.31E-16 |
| Apol7d | 2.280534 | 0.000236 |
| Sgms2 | 2.278245 | 0.001549 |
| Ly6c1 | 2.267608 | 1.07E-44 |
| Mt1 | 2.256099 | 2.60E-28 |
| Tgfbi | 2.248899 | 2.51E-14 |
| Nlrp1b | 2.243521 | 4.35E-08 |
| H2-Q10 | 2.243174 | 8.52E-28 |
| Ly6c2 | 2.238153 | 1.17E-35 |
| Catsperg1 | 2.22967 | 6.92E-06 |
| Ifi27l2a | 2.22947 | 1.32E-14 |
| Mrc1 | 2.225488 | 1.13E-07 |
| Ccl2 | 2.222105 | 0.00263 |
| H2-Q4 | 2.215896 | 3.54E-47 |
| Aoah | 2.213202232 | 2.25E-25 |
| Gm21188 | 2.200538 | 3.01E-07 |
| Kif26b | 2.200531 | 0.027653 |
| Pilra | 2.188723 | 0.000239 |
| Rgmb | 2.184322 | 0.048223 |
| F830208F2 | 2.182816 | 6.86E-06 |

| | | |
|---|---|---|
| Ifitm1 | 2.177789 | 2.58E-25 |
| Tlr11 | 2.17518 | 0.000687 |
| Cxcr3 | 2.164604 | 0.00686 |
| Tmem106a | 2.161254 | 1.04E-24 |
| Axl | 2.1560818 | 7.41E-32 |
| Cysltr1 | 2.155484 | 1.92E-08 |
| Polm | 2.149361519 | 5.47E-20 |
| Rse4 | 2.148677404 | 2.59E-16 |
| Selp | 2.135138 | 1.99E-13 |
| Ms4a6d | 2.13392 | 1.56E-06 |
| Cxcl10 | 2.131502 | 3.96E-06 |
| Tlr8 | 2.131208 | 9.55E-05 |
| Gpr141 | 2.126983 | 1.34E-19 |
| Zfyve9 | 2.123027 | 1.70E-06 |
| Lpl | 2.118385519 | 2.65E-14 |
| Casp12 | 2.111087 | 1.24E-06 |
| Gm21188 | 2.100001 | 3.97E-08 |
| Gbp5 | 2.087313 | 1.32E-35 |
| Gm18852 | 2.08418 | 4.17E-09 |
| Fgd6 | 2.078454677 | 0.034814 |
| Ifi211 | 2.077773 | 6.01E-34 |
| Sycp3 | 2.07497638 | 0.040888 |
| Trem2 | 2.066763 | 1.10E-15 |
| Tctex1d1 | 2.061672 | 2.24E-11 |
| Bst1 | 2.0597 | 0.003511 |
| Gng11 | 2.056642 | 9.06E-06 |
| Oasl2 | 2.054017 | 2.43E-09 |
| Calhm6 | 2.053135 | 3.39E-21 |
| Mmp8 | 2.049376694 | 8.09E-09 |
| Fcgr4 | 2.048973 | 0.013379 |
| Ifitm3 | 2.045552 | 1.97E-50 |
| Grap | 2.044562235 | 1.11E-17 |
| Gbp6 | 2.041059 | 1.76E-05 |
| Lilrb4a | 2.039217 | 3.28E-11 |
| Gm18853 | 2.018766 | 1.89E-08 |
| Mocos | 2.016122 | 3.17E-06 |
| Mcub | 2.012098 | 5.38E-11 |
| Hotairm1 | 2.003157 | 0.016309 |
| Ifi207 | 2.002692 | 3.77E-18 |
| Gbp10 | 2.002296 | 1.13E-07 |
| Ccl6 | 1.997911177 | 2.63E-15 |
| Slc11a1 | 1.995497 | 0.000197 |
| Kcnk13 | 1.993653 | 0.00014 |
| Hp | 1.991274 | 2.41E-18 |
| Arhgef15 | 1.987025 | 0.018076 |
| Prkar1b | 1.980534 | 0.001554 |
| Id1 | 1.977467 | 9.24E-10 |

| | | |
|---|---|---|
| Ndrg2 | 1.975987805 | 0.000885 |
| Hmox1 | 1.963998633 | 0.002094 |
| Rapsn | 1.9634271 | 7.46E-05 |
| Asgr2 | 1.958412 | 0.045076 |
| Slfn5 | 1.947354 | 8.92E-07 |
| Qpct | 1.944894 | 0.001903 |
| Stra6l | 1.93923 | 0.034957 |
| Cebpb | 1.93874 | 1.45E-09 |
| Apoe | 1.937212162 | 4.26E-10 |
| Gstm1 | 1.936505 | 2.25E-13 |
| Lrp4 | 1.935986 | 0.001036 |
| Ms4a3 | 1.925636 | 3.82E-05 |
| Gm15417 | 1.925422 | 0.038214 |
| Mfsd6l | 1.925293 | 0.010038 |
| Fcnb | 1.916892 | 0.00365 |
| Gpr157 | 1.910623 | 2.66E-08 |
| A530040E1 | 1.898538 | 1.78E-12 |
| Pira6 | 1.897392 | 3.21E-10 |
| Selenom | 1.889984 | 8.56E-13 |
| Nrg1 | 1.889451 | 4.70E-24 |
| Aif1 | 1.885359 | 5.25E-33 |
| Gm15448 | 1.88485 | 5.38E-11 |
| Cd300lf | 1.875574 | 7.09E-07 |
| Cac1b | 1.870668139 | 5.13E-07 |
| Pstpip2 | 1.868742 | 8.15E-13 |
| Lgals3 | 1.867303 | 1.66E-14 |
| Glrp1 | 1.863997 | 0.013902 |
| Slamf7 | 1.859596 | 0.01515 |
| Pde7b | 1.856851318 | 6.49E-05 |
| App | 1.854841 | 7.70E-26 |
| Slc39a4 | 1.85337 | 0.024262 |
| Chd7 | 1.848614 | 6.44E-18 |
| Papss2 | 1.843022 | 8.46E-27 |
| Fgl2 | 1.839048 | 1.88E-22 |
| Pde8b | 1.838184012 | 0.005504 |
| Sh2d6 | 1.836859 | 0.00198 |
| Pirb | 1.836663 | 1.39E-11 |
| Ccr1 | 1.835434 | 2.44E-05 |
| Il13ra1 | 1.832817851 | 1.64E-06 |
| Fcor | 1.831861 | 0.042601 |
| cc2 | 1.829345 | 1.05E-07 |
| Apobec1 | 1.814848 | 5.96E-28 |
| Slc22a18 | 1.810373653 | 2.70E-05 |
| Nrp2 | 1.807334 | 3.75E-32 |
| Cd14 | 1.801431 | 0.000295 |
| Itga1 | 1.79652 | 1.20E-17 |
| Tnfsf8 | 1.795296 | 1.52E-08 |

| | | |
|---|---|---|
| Myo18b | 1.788022 | 0.005959 |
| Slfn2 | 1.787971 | 4.27E-12 |
| Hsd11b1 | 1.786388201 | 0.001243 |
| C4a | 1.785749143 | 2.76E-06 |
| Fas | 1.785129 | 0.008024 |
| aa | 1.7812 | 7.61E-13 |
| H2-M3 | 1.780620766 | 1.06E-34 |
| Oasl1 | 1.776556 | 1.25E-05 |
| A530010L1 | 1.773098 | 0.028111 |
| Eps8 | 1.767727785 | 2.16E-12 |
| P2ry6 | 1.765067 | 0.00127 |
| Plppr2 | 1.763257 | 0.021409 |
| Copz2 | 1.759944686 | 9.45E-05 |
| Usp35 | 1.755299 | 0.005985 |
| Tnfsf13 | 1.74498 | 2.04E-10 |
| Csf2ra | 1.74389 | 1.14E-16 |
| ip1 | 1.73998705 | 3.07E-06 |
| Smpdl3a | 1.738784154 | 5.44E-12 |
| Rnf217 | 1.738602 | 7.56E-11 |
| Mok | 1.737312 | 0.036323 |
| Clec5a | 1.733953 | 9.11E-23 |
| Cd302 | 1.733686 | 5.71E-08 |
| Prg3 | 1.730554 | 0.010574 |
| Kcnq3 | 1.729937 | 0.003071 |
| H2-T22 | 1.727852 | 4.50E-57 |
| H2-K1 | 1.72629 | 1.71E-86 |
| Gvin1 | 1.724475 | 7.27E-38 |
| Pyroxd2 | 1.723007 | 0.00155 |
| Clec1b | 1.718889 | 0.027543 |
| Nupr1 | 1.718615 | 0.000168 |
| 1700012B0 | 1.711594 | 1.72E-05 |
| Methig1 | 1.706888 | 0.000143 |
| Nlrc5 | 1.685487 | 1.28E-20 |
| Ang | 1.682758 | 0.01419 |
| Cac1e | 1.682593394 | 0.004939 |
| Myo15 | 1.678118 | 0.018377 |
| Gm4070 | 1.675902 | 4.08E-30 |
| Stxbp6 | 1.674275 | 1.03E-18 |
| Usp44 | 1.672586632 | 0.000611 |
| Oas3 | 1.672537 | 0.034613 |
| ip2 | 1.667575 | 2.17E-21 |
| Hacd4 | 1.666525 | 1.56E-11 |
| Trim6 | 1.665318 | 0.003198 |
| Cyp39a1 | 1.664452 | 0.000245 |
| Nectin2 | 1.651338 | 2.13E-05 |
| Kif19a | 1.651056737 | 0.019908 |
| Ctsc | 1.650955 | 2.00E-16 |

| | | |
|---|---|---|
| 6430548M | 1.647105 | 8.32E-12 |
| Btla | 1.631491 | 4.92E-31 |
| Aldh1l1 | 1.626422 | 0.000218 |
| Klrk1 | 1.625926 | 4.70E-16 |
| Oas1a | 1.625351 | 1.75E-10 |
| AW112010 | 1.614224 | 0.000201 |
| Tgm1 | 1.614134704 | 0.002285 |
| Aatk | 1.606378 | 1.19E-12 |
| Dstn | 1.605452695 | 2.58E-11 |
| Per1 | 1.604402322 | 0.002208 |
| Samd9l | 1.603078 | 7.08E-07 |
| Nhsl2 | 1.602071 | 0.000188 |
| Tnfsfm13 | 1.594187074 | 1.04E-14 |
| H2-Q1 | 1.591573 | 3.48E-48 |
| Plxnb3 | 1.589965 | 0.000862 |
| Pira2 | 1.588358 | 1.31E-09 |
| Tmem38b | 1.581599 | 5.07E-14 |
| Slc15a3 | 1.580512 | 1.17E-05 |
| Tlr1 | 1.579371 | 1.09E-07 |
| Pira1 | 1.573817 | 3.42E-11 |
| Tnfsf15 | 1.571083 | 0.004754 |
| Casp4 | 1.567586 | 5.19E-15 |
| H2-D1 | 1.567234 | 1.35E-95 |
| Stox2 | 1.562328 | 1.26E-11 |
| Xaf1 | 1.55643 | 6.51E-08 |
| Cdiptos | 1.555434 | 0.002892 |
| Msrb1 | 1.549864 | 1.42E-17 |
| Sowahc | 1.549567 | 4.07E-13 |
| F630028O1 | 1.546536 | 2.36E-12 |
| Slc36a2 | 1.542861797 | 0.039948 |
| Pir | 1.540859 | 0.001253 |
| | 1.535413 | 3.09E-07 |
| Ahrr | 1.532840291 | 0.041532 |
| Gramd3 | 1.528893846 | 1.06E-10 |
| Zfp979 | 1.527886 | 0.007128 |
| Cldn15 | 1.525919116 | 4.45E-07 |
| Lrp1 | 1.517345 | 5.70E-13 |
| Dpep2 | 1.515117 | 8.05E-09 |
| Phf24 | 1.514394 | 0.021264 |
| Ccm2l | 1.509303 | 0.000974 |
| Cyp27a1 | 1.508481 | 9.65E-08 |
| F5 | 1.499779 | 0.00013 |
| Gsr | 1.49438 | 5.56E-11 |
| Mefv | 1.489526 | 8.42E-22 |
| Sqor | 1.485431269 | 1.60E-08 |
| Gm8909 | 1.485376 | 1.47E-24 |
| Homer1 | 1.484430382 | 3.49E-11 |

| | | |
|---|---|---|
| Ifi205 | 1.484219 | 8.41E-20 |
| Pld4 | 1.483642 | 3.41E-23 |
| Slfn4 | 1.480983795 | 0.039832 |
| Tbxas1 | 1.480463 | 2.86E-10 |
| Tfec | 1.474753 | 6.29E-17 |
| Galnt9 | 1.473069 | 0.029839 |
| Pira11 | 1.468083 | 2.86E-08 |
| Pygl | 1.467524421 | 5.28E-23 |
| Clec7a | 1.467067 | 1.24E-26 |
| Cnr2 | 1.465552 | 4.05E-06 |
| B4galnt4 | 1.462353 | 8.53E-07 |
| Tnfsf12 | 1.455213 | 1.98E-06 |
| Tsc22d3 | 1.454199 | 2.51E-08 |
| H2-T10 | 1.45286 | 5.43E-15 |
| Mcf2l | 1.450086 | 5.27E-14 |
| Acacb | 1.436915 | 0.000241 |
| Ptgir | 1.436783 | 0.012303 |
| Zbtb7b | 1.434982 | 1.57E-22 |
| Tlr6 | 1.431905 | 0.000303 |
| Ptafr | 1.428019 | 1.42E-06 |
| C1rb | 1.427596 | 0.043597 |
| Mettl7a2 | 1.421253 | 0.001603 |
| Gm807 | 1.416984 | 0.001825 |
| C130050O: | 1.416913 | 5.14E-08 |
| Muc13 | 1.416609 | 6.69E-49 |
| Aldh3b1 | 1.414866 | 1.50E-14 |
| Gm7030 | 1.410917 | 1.35E-27 |
| Slc31a2 | 1.408488 | 2.36E-18 |
| Gm11127 | 1.400809 | 3.32E-19 |
| Gpr35 | 1.400731 | 3.39E-08 |
| Igsf6 | 1.398181 | 3.60E-21 |
| Arhgap26 | 1.391059 | 1.50E-14 |
| Hck | 1.39051408 | 1.22E-13 |
| Dse | 1.38329 | 2.81E-07 |
| Arhgef10l | 1.38172 | 1.81E-14 |
| Camk2d | 1.381608 | 3.09E-11 |
| Adgre1 | 1.38014899 | 2.16E-09 |
| Tnni3 | 1.378825 | 0.00839 |
| Dio2 | 1.377198849 | 0.000648 |
| Ankmy1 | 1.376355 | 0.008189 |
| Mpzl3 | 1.372946 | 0.002027 |
| Elane | 1.364164686 | 0.001315 |
| Ctss | 1.363484 | 1.41E-30 |
| Arhgap24 | 1.360284 | 1.68E-05 |
| Tmem51 | 1.360176 | 3.88E-05 |
| Ttc39c | 1.360061 | 3.23E-07 |
| H2-K2 | 1.349002 | 1.09E-10 |

| | | |
|---|---|---|
| Mgst2 | 1.347924 | 1.36E-05 |
| Clec4e | 1.346841 | 0.002024 |
| ip6 | 1.345721 | 6.05E-07 |
| Mtus1 | 1.343187 | 1.34E-20 |
| Il12rb1 | 1.339841194 | 0.000143 |
| Trem3 | 1.338495 | 0.000101 |
| Il15ra | 1.337863 | 4.44E-13 |
| Tgtp1 | 1.33702 | 3.04E-05 |
| Stx11 | 1.337006 | 2.94E-14 |
| Lair1 | 1.331552 | 2.94E-10 |
| Gca | 1.326675 | 0.000757 |
| Ttc21a | 1.325765 | 0.017106 |
| Pde1b | 1.323142 | 0.000311 |
| Ugt1a7c | 1.320682 | 4.71E-14 |
| ip5 | 1.320279 | 4.73E-06 |
| B4galt6 | 1.318108 | 7.47E-10 |
| B430010I2 | 1.317827 | 0.00023 |
| Ms4a6b | 1.317707 | 7.83E-34 |
| Lrrc25 | 1.316861 | 9.66E-12 |
| Galm | 1.316256 | 0.037005 |
| Ugt1a10 | 1.313183 | 4.13E-09 |
| Igtp | 1.312987 | 1.03E-08 |
| Ugt1a6a | 1.312944 | 3.79E-11 |
| Slc7a7 | 1.303704731 | 2.18E-10 |
| Chpt1 | 1.30215 | 5.41E-05 |
| Atp11a | 1.29807 | 6.78E-06 |
| Lbp | 1.295416104 | 9.39E-13 |
| Hnmt | 1.294539 | 0.001378 |
| Crybg1 | 1.293099702 | 1.18E-05 |
| B230217C1 | 1.290627 | 1.31E-07 |
| 1700034H1 | 1.289524 | 0.045747 |
| Klf9 | 1.289292 | 0.010438 |
| Fcgrt | 1.288806638 | 7.44E-16 |
| Tmem71 | 1.288663 | 0.000742 |
| Gas2l1 | 1.2865 | 0.004646 |
| Ccl9 | 1.286468533 | 2.34E-12 |
| Cyp4f18 | 1.285893455 | 1.19E-11 |
| Galc | 1.285213112 | 1.24E-06 |
| Psap | 1.277553139 | 4.98E-35 |
| Nod1 | 1.274937 | 9.36E-29 |
| Fam129a | 1.274771 | 1.09E-07 |
| C1galt1c1 | 1.274149 | 2.05E-15 |
| Oas2 | 1.27343 | 0.000272 |
| Pi16 | 1.273382 | 7.09E-07 |
| Cask | 1.272578 | 6.31E-21 |
| Irgm2 | 1.272481 | 2.74E-06 |
| Plaur | 1.272118 | 3.54E-12 |

| | | |
|---|---|---|
| Tnfrsf1b | 1.264295 | 5.76E-23 |
| Ifngr1 | 1.264116911 | 1.55E-17 |
| Pros1 | 1.263353 | 1.08E-06 |
| Arsg | 1.260704532 | 0.047667 |
| Gbp7 | 1.260684 | 4.75E-32 |
| Igf2bp3 | 1.260471 | 0.009313 |
| Krt80 | 1.259943 | 0.004888 |
| Cracr2b | 1.258035 | 0.01615 |
| Fcgr2b | 1.25556 | 5.76E-23 |
| Lilra6 | 1.255407 | 0.049426 |
| Ugt1a1 | 1.252948 | 1.02E-09 |
| Ugt1a6b | 1.250812 | 1.93E-10 |
| Olfr56 | 1.25023 | 0.013575 |
| Gas7 | 1.249956 | 4.38E-07 |
| Cd300c2 | 1.249932 | 0.000383 |
| Ugt1a5 | 1.249925 | 1.15E-09 |
| Plx1 | 1.249 | 0.00225 |
| Ugt1a2 | 1.248825 | 1.32E-09 |
| Ltb4r1 | 1.248758 | 0.015127 |
| Nectin4 | 1.24806135 | 0.003147 |
| v2 | 1.247959 | 3.10E-11 |
| C1qtnf6 | 1.246683503 | 2.60E-05 |
| Ugt1a9 | 1.245313 | 1.20E-09 |
| Slc22a4 | 1.244172527 | 8.72E-05 |
| Kif1b | 1.242108 | 6.32E-11 |
| Stat1 | 1.239253 | 8.15E-13 |
| Irf7 | 1.238449 | 3.50E-07 |
| Acvrl1 | 1.236137 | 2.25E-07 |
| Ldlrad3 | 1.233509 | 2.77E-08 |
| Cyp4v3 | 1.232845 | 2.39E-06 |
| Zfp287 | 1.229660901 | 0.000281 |
| Tspan17 | 1.229183 | 2.58E-05 |
| Nostrin | 1.228522 | 1.72E-12 |
| Hoxb3 | 1.225697 | 0.024366 |
| Scnn1a | 1.225372 | 2.72E-05 |
| Pitpnm1 | 1.222339 | 1.74E-13 |
| Tap1 | 1.219934 | 5.50E-28 |
| Samhd1 | 1.217451 | 7.50E-21 |
| Rwdd2a | 1.215345 | 0.024262 |
| Slc2a6 | 1.214862 | 5.14E-23 |
| Tgtp2 | 1.212863 | 9.16E-06 |
| Gm10693 | 1.212786 | 0.00096 |
| Padi4 | 1.211983 | 2.85E-07 |
| Cttnbp2nl | 1.211662 | 0.004349 |
| Ldlr | 1.209097 | 2.10E-10 |
| Ssc4d | 1.20638 | 9.32E-07 |
| Dram1 | 1.204438152 | 1.31E-11 |

| | | |
|---|---|---|
| Sdc3 | 1.202946 | 0.000418 |
| Kcnj2 | 1.199753 | 2.05E-05 |
| Sema6b | 1.196555407 | 1.43E-08 |
| Dmxl2 | 1.195606 | 3.60E-06 |
| Soat1 | 1.192741 | 8.69E-13 |
| Pik3r6 | 1.190622 | 3.40E-15 |
| Prdx5 | 1.189442 | 1.54E-19 |
| Spata13 | 1.186897345 | 1.46E-15 |
| Hpn | 1.184919967 | 2.36E-10 |
| B2m | 1.184144 | 1.70E-45 |
| Sulf2 | 1.183184471 | 4.02E-13 |
| 1-Mar | 1.181936 | 1.23E-07 |
| Ikbke | 1.179629 | 5.52E-13 |
| Plxdc1 | 1.179611348 | 4.45E-09 |
| Rras | 1.177927 | 0.001354 |
| Icosl | 1.174702795 | 6.49E-11 |
| Ifih1 | 1.173293 | 5.93E-06 |
| S100a6 | 1.172950296 | 8.68E-09 |
| C1ra | 1.172852 | 0.011039 |
| Plaat3 | 1.170359 | 1.82E-05 |
| Sat1 | 1.168518 | 2.27E-09 |
| Fgd4 | 1.158483 | 3.08E-07 |
| Lst1 | 1.157391 | 1.39E-09 |
| Fstl1 | 1.154327 | 0.039668 |
| Emilin2 | 1.148569 | 2.86E-11 |
| Pld1 | 1.145213 | 0.000153 |
| Lgals3bp | 1.139346 | 5.08E-06 |
| Sema4a | 1.137808 | 3.61E-16 |
| Lrrk2 | 1.136479 | 3.36E-09 |
| Rnf144b | 1.132355 | 1.67E-07 |
| Ccr2 | 1.131892 | 6.74E-20 |
| Glcci1 | 1.130797 | 0.000159 |
| Mgl2 | 1.130463 | 0.00049 |
| Khk | 1.12973 | 2.89E-11 |
| Gbp3 | 1.128128 | 1.37E-22 |
| Mical2 | 1.122338 | 0.000647 |
| Tmem154 | 1.120892 | 0.000174 |
| Fam241a | 1.120265 | 8.43E-07 |
| Cpeb2 | 1.119109 | 9.32E-06 |
| Osm | 1.118653 | 0.047269 |
| Vsir | 1.115894253 | 7.15E-10 |
| Grk5 | 1.115257137 | 0.012952 |
| Tent5a | 1.112456 | 1.35E-21 |
| Tmem63a | 1.109168 | 4.35E-08 |
| Loxl3 | 1.105215289 | 9.56E-06 |
| Fosl2 | 1.101884 | 9.23E-12 |
| Anxa5 | 1.100602 | 5.46E-19 |

| | | |
|---|---|---|
| Tbkbp1 | 1.098255 | 1.73E-22 |
| Ube2l6 | 1.097631 | 0.000589 |
| Gpr65 | 1.095818534 | 2.60E-05 |
| Olfm1 | 1.094087 | 9.73E-07 |
| Tlr9 | 1.093789 | 2.85E-07 |
| Phf11b | 1.09164 | 0.001637 |
| Pmaip1 | 1.089878 | 0.004578 |
| Ralb | 1.089137792 | 1.15E-19 |
| Susd3 | 1.08701673 | 9.57E-06 |
| Ephx1 | 1.085242 | 0.018522 |
| Idh1 | 1.079901 | 2.68E-21 |
| Map3k6 | 1.079737 | 2.06E-05 |
| H2-DMb1 | 1.076459 | 1.10E-15 |
| Abcd2 | 1.075216 | 3.95E-07 |
| Ptpro | 1.074885 | 8.06E-08 |
| Stard9 | 1.07403 | 1.12E-05 |
| Tom1 | 1.073557 | 1.18E-15 |
| Glrx | 1.073004274 | 5.47E-07 |
| Kalrn | 1.06448 | 1.70E-09 |
| Scpep1 | 1.063984937 | 7.72E-20 |
| Fbxl2 | 1.061853 | 0.015079 |
| Dusp22 | 1.061747 | 7.55E-11 |
| Itgb5 | 1.060956 | 5.86E-05 |
| Dok3 | 1.059816 | 4.68E-09 |
| Stom | 1.058904 | 1.84E-30 |
| Cd68 | 1.057503648 | 6.69E-23 |
| Usp18 | 1.054877 | 0.019319 |
| Klf4 | 1.05209779 | 1.97E-22 |
| Dgkg | 1.05157 | 0.001342 |
| Trafd1 | 1.04772 | 5.08E-13 |
| Phf11c | 1.047252 | 0.03111 |
| Parp14 | 1.043132 | 1.92E-12 |
| Mospd2 | 1.041232 | 2.59E-05 |
| Epsti1 | 1.041058975 | 3.04E-11 |
| Rassf4 | 1.036213 | 6.57E-13 |
| Stard8 | 1.035583 | 6.32E-06 |
| Nlrp3 | 1.033083 | 7.48E-12 |
| Anxa1 | 1.032626 | 3.39E-08 |
| Plcb1 | 1.030008 | 0.028213 |
| Cers6 | 1.024233 | 2.13E-07 |
| BC147527 | 1.023549 | 0.010756 |
| Adam19 | 1.020860441 | 0.006864 |
| Hpse | 1.018039 | 6.35E-15 |
| Ifi47 | 1.018021 | 7.41E-07 |
| Bcl3 | 1.017986 | 2.97E-07 |
| Mettl7a1 | 1.01679 | 1.31E-08 |
| Lcp2 | 1.016156592 | 8.94E-08 |

| | | |
|---|---|---|
| Cd52 | 1.015974985 | 1.54E-14 |
| Itgb7 | 1.015714267 | 2.65E-11 |
| Lgr4 | 1.014658 | 0.004917 |
| Dusp3 | 1.013839421 | 2.30E-17 |
| Slc8b1 | 1.011262 | 8.33E-16 |
| Mmp19 | 1.009034 | 0.000175 |
| Glis3 | 1.008213 | 0.009313 |
| Rap1gap2 | 1.004713 | 0.000173 |
| Mx1 | 1.004204311 | 4.35E-06 |
| Fkbp1b | 1.001925352 | 5.92E-09 |
| 10-Sep | 1.000493535 | 8.08E-05 |
| Tmem220 | 1.00021 | 0.02674 |
| A530032D1 | 1.000182 | 4.04E-05 |
| Fads2 | -1.000978 | 0.015334 |
| Paip2b | -1.002449 | 3.94E-06 |
| Flt3 | -1.005107 | 1.39E-09 |
| Hook1 | -1.00653 | 0.00043 |
| Gnb4 | -1.007888 | 0.041532 |
| Tsc22d1 | -1.010094435 | 1.47E-05 |
| Lrfn4 | -1.016903 | 0.000709 |
| Rnf141 | -1.023314 | 6.55E-06 |
| Spib | -1.02445974 | 0.047781 |
| Plekhh1 | -1.025782 | 0.000118 |
| Kif3a | -1.026933782 | 0.000262 |
| Cyfip2 | -1.02714726 | 1.08E-13 |
| Runx2 | -1.029755 | 4.96E-05 |
| Oscp1 | -1.033411 | 0.012723 |
| Repin1 | -1.033537 | 4.29E-07 |
| Cbx2 | -1.034874 | 0.001746 |
| Aldh7a1 | -1.039094 | 0.00318 |
| Kank2 | -1.04293 | 8.27E-05 |
| Tspyl4 | -1.04357 | 0.017273 |
| Zfp608 | -1.044819 | 1.82E-05 |
| Dcaf1 | -1.046068 | 2.13E-07 |
| 4930555A0 | -1.048227 | 0.047043 |
| Cep19 | -1.051304 | 3.16E-07 |
| Armcx2 | -1.05209 | 0.00213 |
| Egln3 | -1.054062 | 0.000647 |
| Fam234b | -1.055442 | 0.000772 |
| Adat1 | -1.055518 | 0.012567 |
| Zfhx2 | -1.064919 | 0.011033 |
| Cbarp | -1.06521 | 0.007422 |
| Tpk1 | -1.065901 | 0.010601 |
| Srl | -1.066593 | 0.000316 |
| Smo | -1.067645466 | 4.60E-09 |
| Prxl2a | -1.074637247 | 0.006089 |
| Erg | -1.079866 | 0.000399 |

| | | |
|---|---|---|
| Fam129c | -1.080504 | 0.007673 |
| Camk1d | -1.08238 | 8.68E-09 |
| 2500004C( | -1.08297 | 0.034979 |
| Zdhhc8 | -1.084212 | 9.37E-10 |
| Slc43a1 | -1.085031 | 5.36E-05 |
| Hsph1 | -1.085282 | 0.041241 |
| Ppp1r13l | -1.086027 | 0.002169 |
| Zfp566 | -1.089106 | 0.032281 |
| Ccnd2 | -1.091290107 | 3.72E-05 |
| Carhsp1 | -1.093462275 | 0.006837 |
| Atp2b4 | -1.095325 | 8.80E-16 |
| Ahi1 | -1.097687341 | 0.001316 |
| Strbp | -1.100165 | 1.78E-12 |
| Clstn1 | -1.100242 | 2.00E-05 |
| Cd320 | -1.101105957 | 0.001721 |
| Zfp773 | -1.103494 | 0.020001 |
| Thnsl1 | -1.108156 | 0.002503 |
| Slc14a1 | -1.110734 | 0.001696 |
| Inka1 | -1.113083 | 3.68E-06 |
| Hmces | -1.114284 | 3.98E-06 |
| Chst11 | -1.119212 | 2.25E-06 |
| 2810408A1 | -1.120299682 | 0.010649 |
| Tapt1 | -1.121374 | 5.99E-06 |
| Tgif2 | -1.122271 | 2.49E-05 |
| Kif7 | -1.124361 | 0.004709 |
| Dock9 | -1.127756 | 1.10E-06 |
| Pard3b | -1.128545 | 0.005439 |
| Stat4 | -1.1288 | 0.000142 |
| Slc25a23 | -1.129145 | 0.001569 |
| Gpsm1 | -1.13207 | 7.30E-05 |
| Wdr60 | -1.137439 | 0.011232 |
| Clcn2 | -1.138438 | 0.022004 |
| Rhoh | -1.13883 | 1.39E-12 |
| Nkg7 | -1.143830469 | 0.000228 |
| Pde7a | -1.145633 | 7.56E-08 |
| Lca5 | -1.146735 | 0.042335 |
| Ltb | -1.152064 | 0.02452 |
| Gpr171 | -1.152794 | 2.10E-07 |
| Rac3 | -1.154786151 | 0.030026 |
| Tet1 | -1.155223 | 0.005671 |
| Bcl9l | -1.155951 | 1.85E-06 |
| Fstl3 | -1.157667306 | 4.34E-05 |
| Msi2 | -1.158492 | 6.42E-06 |
| Prss57 | -1.159759758 | 6.38E-09 |
| Zdhhc15 | -1.162562 | 0.001532 |
| Flnb | -1.163595 | 2.81E-06 |
| Apcdd1 | -1.172937 | 0.00442 |

| | | |
|---|---|---|
| Tmem238 | -1.174187 | 0.025296 |
| Plcg1 | -1.174264892 | 1.08E-06 |
| Ddx58 | -1.175119 | 0.000793 |
| Slc25a37 | -1.177073 | 1.20E-07 |
| Tbc1d12 | -1.179619 | 0.029071 |
| Nos1ap | -1.179699 | 1.99E-07 |
| Rpgrip1 | -1.180202 | 3.44E-15 |
| Ccnd1 | -1.180604 | 2.03E-05 |
| Jcad | -1.184006 | 0.028275 |
| Slc5a3 | -1.186652 | 0.001381 |
| Pard6g | -1.187808 | 0.026675 |
| Hmgn1 | -1.190568 | 4.56E-18 |
| Prpf40b | -1.190721 | 3.06E-07 |
| Epb41l4b | -1.191974 | 2.01E-05 |
| Ap1s3 | -1.195145 | 0.000111 |
| Adgra3 | -1.201559 | 0.000886 |
| Sh2d5 | -1.206205 | 8.19E-09 |
| Btbd3 | -1.209065 | 5.45E-05 |
| Mllt6 | -1.209555 | 3.77E-18 |
| Retreg1 | -1.216825546 | 2.70E-07 |
| Actn1 | -1.218794291 | 0.003921 |
| Arvcf | -1.219544684 | 0.026776 |
| Slc27a1 | -1.230223 | 2.04E-09 |
| Otub2 | -1.23093516 | 0.008714 |
| Tdrkh | -1.231001 | 6.78E-05 |
| Tbc1d10c | -1.234837 | 2.56E-07 |
| Tfrc | -1.248169 | 1.38E-08 |
| Sox12 | -1.250949 | 0.000173 |
| Plx3 | -1.253391 | 0.033937 |
| Thsd1 | -1.254908 | 0.000326 |
| Ddah2 | -1.265491851 | 0.000202 |
| Gm2011 | -1.2662 | 0.023281 |
| Clstn3 | -1.267620119 | 0.005783 |
| Amt | -1.273868 | 0.048606 |
| Cacnb2 | -1.274251 | 0.028571 |
| Ppp1r16b | -1.278891 | 0.00061 |
| Paqr8 | -1.281489 | 0.00265 |
| Ripor1 | -1.28608 | 6.98E-14 |
| Ppp3cc | -1.292179229 | 0.002623 |
| Maml3 | -1.292553 | 1.19E-06 |
| Jag2 | -1.292576305 | 0.034814 |
| Spry2 | -1.293684693 | 0.003432 |
| Zscan18 | -1.296557 | 0.005047 |
| Rtp4 | -1.297426 | 0.00134 |
| B630019K0 | -1.301313 | 0.033613 |
| Ahdc1 | -1.308628 | 4.62E-05 |
| Tespa1 | -1.309002 | 3.62E-07 |

| | | |
|---|---|---|
| Ctr9 | -1.310185652 | 1.86E-32 |
| Msrb2 | -1.317844 | 0.006668 |
| Djb2 | -1.326657 | 0.004774 |
| Zscan2 | -1.327291 | 1.01E-05 |
| Klrd1 | -1.330537 | 0.038752 |
| Ptp4a3 | -1.331953 | 4.96E-25 |
| Irak1bp1 | -1.335656 | 0.000664 |
| Ldlrad4 | -1.336518 | 0.001258 |
| Pacsin1 | -1.349878 | 0.040981 |
| Pomgnt2 | -1.351898 | 0.000572 |
| Clcf1 | -1.358387 | 0.001212 |
| Lmntd2 | -1.3616 | 0.03276 |
| Spag1 | -1.362417 | 0.031069 |
| Stxbp4 | -1.362523689 | 0.014975 |
| Kdm5b | -1.363648 | 0.038632 |
| Prx | -1.372555 | 0.01366 |
| Socs5 | -1.373064 | 1.62E-06 |
| Mex3b | -1.376919 | 4.04E-06 |
| Satb1 | -1.383001 | 1.38E-09 |
| Zfpm1 | -1.389981 | 0.018844 |
| Fgr | -1.394501 | 5.15E-05 |
| Ets2 | -1.396236 | 0.018731 |
| Tmem8b | -1.402438 | 0.004731 |
| Vangl2 | -1.402504 | 0.001953 |
| Sema4c | -1.405943 | 1.20E-06 |
| Dab2ip | -1.411769 | 1.27E-27 |
| Ypel1 | -1.418399 | 0.000239 |
| Zfp239 | -1.425314 | 0.000802 |
| Cyp2j6 | -1.432936 | 0.030887 |
| Spag4 | -1.438821 | 0.036808 |
| Tnfrsf18 | -1.43925 | 0.004601 |
| Cand2 | -1.447241 | 4.42E-06 |
| Rundc3a | -1.448098392 | 0.020502 |
| Ptpn3 | -1.457849 | 0.020098 |
| Srgap3 | -1.458151 | 4.84E-08 |
| Adgrl1 | -1.466416671 | 2.73E-07 |
| Lhfpl2 | -1.468527 | 0.000604 |
| Tmem119 | -1.481971 | 1.76E-06 |
| Tle2 | -1.48261 | 5.26E-05 |
| Tal1 | -1.485748 | 7.20E-05 |
| Gm16867 | -1.487212 | 1.06E-11 |
| Ninl | -1.494636 | 0.001654 |
| Gcnt2 | -1.494786376 | 1.91E-25 |
| Kif17 | -1.497103 | 7.10E-09 |
| Bdh1 | -1.497675 | 0.009972 |
| Zbtb10 | -1.508494 | 0.000548 |
| Tes | -1.509295 | 1.99E-08 |

| | | |
|---|---|---|
| Dusp9 | -1.523199 | 0.036777 |
| Afap1 | -1.523231 | 0.000648 |
| Cd27 | -1.527451 | 9.09E-14 |
| Itgax | -1.531003 | 0.000131 |
| Rtn4rl1 | -1.533387 | 2.72E-05 |
| Pdgfrb | -1.53636 | 3.85E-05 |
| Dbn1 | -1.541633 | 0.000782 |
| Cdh24 | -1.544578 | 0.000825 |
| Ednrb | -1.548902989 | 0.003161 |
| Ttyh2 | -1.552949 | 1.79E-05 |
| Sox4 | -1.556391 | 0.004917 |
| Clnk | -1.569257 | 0.000788 |
| Cd34 | -1.572908905 | 1.79E-13 |
| Mast4 | -1.580239 | 0.01615 |
| Mdfi | -1.588198 | 0.019102 |
| Cxxc5 | -1.598312 | 8.56E-09 |
| Sall2 | -1.599586 | 0.002341 |
| Btnl9 | -1.600174 | 0.004049 |
| Gstm5 | -1.608218875 | 3.53E-05 |
| Tspan13 | -1.61474821 | 3.06E-05 |
| Zfp521 | -1.615308 | 9.85E-08 |
| Ptpdc1 | -1.619155 | 0.003478 |
| Vamp5 | -1.628006 | 0.00096 |
| Fads3 | -1.634026 | 2.73E-07 |
| Slc22a17 | -1.637746455 | 0.006451 |
| Peak1 | -1.639568 | 1.64E-08 |
| Evpl | -1.641444 | 7.86E-06 |
| Gm16897 | -1.65245 | 1.19E-05 |
| H1f10 | -1.653128 | 0.000483 |
| Ankrd6 | -1.669609 | 0.037747 |
| Bach2 | -1.672594 | 2.64E-07 |
| Reep1 | -1.675733 | 0.030599 |
| Zfp507 | -1.677241 | 3.84E-06 |
| Adgrg3 | -1.678851 | 2.73E-18 |
| Serpinh1 | -1.683049 | 0.003565 |
| Cd59a | -1.686208 | 0.013962 |
| Marcksl1 | -1.688315 | 8.95E-10 |
| Dusp6 | -1.693264909 | 0.000221 |
| Pmepa1 | -1.693671 | 0.002042 |
| Myh10 | -1.694914553 | 0.005115 |
| Myl4 | -1.69654 | 2.13E-07 |
| Peg13 | -1.699558 | 1.47E-07 |
| Trp53i11 | -1.705816 | 6.89E-09 |
| Havcr2 | -1.707775484 | 0.002395 |
| Adgrl2 | -1.718306 | 0.000206 |
| Slc1a4 | -1.718656099 | 0.026776 |
| Pdzd4 | -1.720013921 | 1.00E-08 |

| | | |
|---|---|---|
| Fgfr1 | -1.720992 | 0.031803 |
| Il12a | -1.725894 | 4.78E-13 |
| Armcx1 | -1.744412 | 0.010562 |
| 2700081O1 | -1.758645 | 4.06E-15 |
| F2r | -1.772328 | 0.015906 |
| Tnfrsf13c | -1.773613 | 1.65E-08 |
| Klhl12 | -1.784573 | 7.43E-07 |
| Mmp11 | -1.787715442 | 0.007759 |
| Cnn3 | -1.789602 | 1.94E-14 |
| Dusp5 | -1.791101 | 2.93E-09 |
| Dzip1 | -1.793499 | 0.024402 |
| Fam174b | -1.798622 | 0.000184 |
| Rundc3b | -1.804669 | 0.001529 |
| Whrn | -1.805988 | 2.09E-07 |
| Gbp8 | -1.806326 | 1.52E-05 |
| Rbpms | -1.808421 | 8.44E-07 |
| Itgad | -1.811025 | 0.00319 |
| Adora2a | -1.811606312 | 0.017924 |
| Uggt2 | -1.813236 | 0.007437 |
| Chst15 | -1.823275 | 7.71E-14 |
| Lrig1 | -1.824 | 0.019752 |
| Tuba8 | -1.837241 | 0.00099 |
| Cdc42bpa | -1.840373 | 3.99E-05 |
| Gm5086 | -1.844852 | 0.020742 |
| Basp1 | -1.853495 | 1.79E-06 |
| Tfr2 | -1.85357 | 0.007539 |
| Cldn34c1 | -1.855012 | 0.011057 |
| Daf4 | -1.855452 | 0.000592 |
| Baiap3 | -1.860912 | 2.15E-06 |
| Gja1 | -1.862591 | 0.016036 |
| Il27ra | -1.866516808 | 0.000829 |
| Ccdc63 | -1.867217 | 0.02674 |
| Cgnl1 | -1.868506 | 0.029505 |
| Dipk1b | -1.870137 | 0.000439 |
| Il18rap | -1.873974 | 3.08E-17 |
| Traf4 | -1.877943401 | 0.000328 |
| Tmem231 | -1.88678 | 0.000365 |
| Gm12504 | -1.899677 | 0.023756 |
| Slc16a11 | -1.904619 | 0.006227 |
| Zfp459 | -1.905565 | 0.018254 |
| Ctxn1 | -1.909938 | 1.53E-05 |
| Epb41l3 | -1.916172 | 0.001884 |
| Cd81 | -1.919091 | 2.13E-17 |
| Gm3383 | -1.923476 | 0.00903 |
| Gm2974 | -1.927914 | 0.013941 |
| Tanc2 | -1.928629 | 6.35E-05 |
| B3gnt7 | -1.933081 | 0.00012 |

| | | |
|---|---|---|
| Sv2a | -1.933756 | 4.12E-05 |
| Cd209a | -1.934973 | 3.73E-05 |
| Apbb1 | -1.942357 | 0.000144 |
| Ccdc9b | -1.947617 | 0.003897 |
| Sorbs3 | -1.948816837 | 0.000223 |
| Clic5 | -1.950927 | 0.005165 |
| Wdr78 | -1.956793 | 0.000403 |
| Eya1 | -1.964795 | 0.001193 |
| Ptger3 | -1.967877 | 0.000118 |
| Zeb1 | -1.976728 | 2.24E-10 |
| Drc7 | -1.982878 | 1.55E-06 |
| Lm | -1.991157 | 0.025209 |
| Epha7 | -1.992288 | 0.001087 |
| Snn | -1.995213 | 6.94E-08 |
| Coro2b | -1.995922 | 0.021836 |
| Jakmip1 | -2.017469 | 0.000786 |
| Adam11 | -2.018145073 | 8.80E-16 |
| Armcx4 | -2.021736 | 7.54E-08 |
| Ephb6 | -2.021845 | 0.021766 |
| Fbln1 | -2.030145319 | 0.043601 |
| Azin2 | -2.049804 | 2.10E-08 |
| Epha2 | -2.052540228 | 8.16E-07 |
| Plod2 | -2.052876 | 0.016105 |
| Podxl2 | -2.057175 | 0.005647 |
| Ccr7 | -2.057817 | 0.00121 |
| Mctp2 | -2.059904 | 2.13E-05 |
| Crim1 | -2.067186 | 0.000648 |
| Abtb2 | -2.068231 | 0.031364 |
| Them7 | -2.07834 | 0.008821 |
| Ncs1 | -2.080916 | 9.01E-05 |
| Car3 | -2.081507 | 0.002705 |
| Rhbdl3 | -2.083578809 | 0.002739 |
| Pygm | -2.08583 | 0.000252 |
| Entpd4b | -2.093075295 | 4.35E-10 |
| Apbb2 | -2.094658 | 1.63E-06 |
| Pde5a | -2.098043 | 0.034193 |
| Shf | -2.100233 | 0.00157 |
| Entpd4 | -2.100661 | 3.58E-09 |
| Wnt10b | -2.10535 | 0.012689 |
| Ctnnd2 | -2.111268986 | 6.27E-05 |
| Ephb2 | -2.115287 | 1.93E-10 |
| Nudt12 | -2.121176 | 1.17E-07 |
| Ccdc8 | -2.123572 | 0.00025 |
| Akap12 | -2.129554 | 0.009972 |
| Marcks | -2.145211 | 2.09E-07 |
| Aff3 | -2.150382 | 1.29E-05 |
| Klk8 | -2.15344 | 7.23E-21 |

| | | |
|---|---|---|
| Dja4 | -2.154506 | 0.006706 |
| D630045J1 | -2.166907 | 1.03E-07 |
| Otud7b | -2.167149 | 7.80E-10 |
| Cobll1 | -2.173246 | 3.53E-09 |
| Frmd6 | -2.176685 | 5.39E-09 |
| Etv5 | -2.188714472 | 8.49E-08 |
| Smpd3 | -2.197215 | 5.84E-05 |
| Prr36 | -2.206799 | 0.005671 |
| Abcb4 | -2.208172 | 0.000702 |
| Glis2 | -2.214304211 | 2.54E-07 |
| Adam22 | -2.225997 | 0.002584 |
| Gata2 | -2.238663305 | 4.63E-05 |
| St3gal6 | -2.263186 | 0.037417 |
| Kcb3 | -2.271698723 | 8.75E-05 |
| Plk2 | -2.272632217 | 0.021512 |
| Ccr9 | -2.273571 | 4.26E-08 |
| Cnrip1 | -2.276599 | 0.00183 |
| H2-Ob | -2.28549 | 1.96E-14 |
| Slain1 | -2.291092 | 2.38E-11 |
| Dcbld2 | -2.297908 | 0.004076 |
| Ptprf | -2.313527 | 3.87E-10 |
| C530008M | -2.319124 | 0.006839 |
| Bex4 | -2.319942 | 0.007611 |
| Mrgpre | -2.33356 | 4.27E-05 |
| Djc6 | -2.338508 | 0.023117 |
| Dclk2 | -2.340574 | 1.75E-10 |
| Adcy6 | -2.344883 | 6.52E-15 |
| Slc22a23 | -2.352811 | 0.001742 |
| Scml4 | -2.378024 | 0.044179 |
| Tspan2 | -2.390958 | 7.39E-51 |
| Ef4 | -2.404083 | 6.78E-05 |
| Klf1 | -2.406422 | 0.039386 |
| Tcaf1 | -2.407074 | 0.004522 |
| D930048N: | -2.41399 | 0.000254 |
| Cass4 | -2.41817 | 0.000214 |
| Magee2 | -2.426604 | 0.002331 |
| Maged1 | -2.435097 | 1.62E-07 |
| Bmp1 | -2.435718291 | 1.03E-06 |
| Bfsp2 | -2.442166 | 3.98E-09 |
| Siglech | -2.454893 | 1.23E-36 |
| Pcbd1 | -2.4622783 | 0.044661 |
| 1700001O: | -2.46725 | 0.017073 |
| Casc4 | -2.475747 | 1.47E-07 |
| Kcnk5 | -2.480265 | 0.000508 |
| Zcchc18 | -2.485597 | 7.13E-05 |
| Robo4 | -2.497072 | 8.41E-05 |
| Uaca | -2.497414 | 0.000316 |

| | | |
|---|---|---|
| Map1a | -2.505605 | 0.040107 |
| Tacc2 | -2.508814 | 5.75E-15 |
| Fam171a2 | -2.511081 | 0.001451 |
| Sytl4 | -2.525204 | 0.037346 |
| Nefh | -2.528981469 | 1.59E-07 |
| Phlda3 | -2.541926 | 1.78E-09 |
| Slc35f2 | -2.546333 | 7.78E-05 |
| AI854703 | -2.570675 | 0.000731 |
| Uchl1 | -2.574556 | 0.018803 |
| Faah | -2.589983 | 1.71E-09 |
| Aqp9 | -2.595355 | 0.000895 |
| Il1r1 | -2.606565 | 2.64E-21 |
| Ptprcap | -2.636577 | 7.52E-13 |
| Mex3a | -2.653563 | 4.87E-07 |
| 2900026A( | -2.655259 | 2.67E-13 |
| Vegfa | -2.655523 | 0.001113 |
| Thbs3 | -2.666311 | 0.000609 |
| Adgrg1 | -2.67492 | 1.34E-29 |
| Il17rb | -2.683986066 | 4.45E-16 |
| Med12l | -2.686492 | 1.43E-08 |
| Paqr5 | -2.68961 | 1.39E-16 |
| Rab6b | -2.700713 | 0.003798 |
| Atp1b1 | -2.705824 | 1.23E-18 |
| Sdc4 | -2.707548301 | 4.10E-06 |
| N4bp3 | -2.70885837 | 1.55E-17 |
| Rab37 | -2.714336256 | 5.09E-35 |
| Sytl1 | -2.720329 | 3.41E-14 |
| Camk2b | -2.723026 | 1.19E-05 |
| Gm3696 | -2.729407 | 0.021449 |
| Itm2a | -2.744824 | 0.012013 |
| Rasd1 | -2.757542 | 0.015809 |
| Traf1 | -2.760012 | 0.009313 |
| Cox6a2 | -2.781349 | 1.37E-08 |
| Nedd4 | -2.810531 | 5.74E-07 |
| Gm5124 | -2.811372 | 0.003494 |
| Zfp2 | -2.816878 | 2.43E-09 |
| Rasl11b | -2.817442 | 0.008573 |
| Tnik | -2.817721 | 0.03064 |
| Cecr2 | -2.825448 | 0.009988 |
| Synpo2l | -2.831634 | 0.049451 |
| Rgl1 | -2.839353 | 0.000451 |
| Ets1 | -2.846789 | 4.27E-06 |
| Peg12 | -2.868878 | 1.43E-05 |
| Emid1 | -2.882923 | 1.40E-23 |
| Tert | -2.901487294 | 0.000648 |
| Slc12a5 | -2.905639299 | 0.001072 |
| Egfl7 | -2.906325 | 5.02E-50 |

| | | |
|---|---|---|
| Mrvi1 | -2.915741398 | 3.50E-07 |
| Synpo2 | -2.926804 | 1.32E-08 |
| Ociad2 | -2.934878 | 4.72E-08 |
| Ly6d | -2.940824 | 6.01E-20 |
| Prkca | -2.94988 | 1.77E-09 |
| Mzb1 | -2.960349 | 0.029168 |
| Il18r1 | -2.976414 | 0.015453 |
| Depp1 | -2.980193 | 1.17E-06 |
| Sema3d | -2.989889 | 0.018292 |
| Car2 | -2.992147 | 0.002193 |
| Gpr25 | -3.006822 | 0.01134 |
| Cd6 | -3.015333 | 2.44E-21 |
| Gprc5b | -3.02533927 | 2.64E-09 |
| Xrcc5 | -3.028361 | 0.002635 |
| Kcng1 | -3.033127 | 3.44E-14 |
| Mn1 | -3.061385 | 1.05E-07 |
| 1700001C1 | -3.063277 | 0.005689 |
| Dscam | -3.067043 | 0.035567 |
| Sept1 | -3.067622713 | 5.31E-36 |
| Serpinb6b | -3.078291 | 0.010851 |
| Fam184a | -3.080199473 | 0.000884 |
| Gpm6b | -3.088854 | 0.036384 |
| Tanc1 | -3.098212 | 3.98E-12 |
| Abcg4 | -3.102449 | 0.025296 |
| Ak4 | -3.112857 | 1.02E-05 |
| Vmn2r29 | -3.118156 | 0.001482 |
| B230217O: | -3.12061 | 0.018377 |
| Gstp3 | -3.123939 | 0.020447 |
| Rcor2 | -3.141539 | 2.38E-07 |
| Zap70 | -3.147544 | 1.49E-05 |
| cad | -3.151357 | 0.023583 |
| Cd96 | -3.158033 | 1.03E-12 |
| Gimap9 | -3.163034 | 2.26E-09 |
| St6gal1 | -3.170916 | 0.002781 |
| Gm2897 | -3.175026 | 0.003216 |
| Cmah | -3.186183914 | 2.43E-10 |
| Arhgdig | -3.19863 | 0.028689 |
| Adam2 | -3.201828092 | 3.82E-05 |
| Tmem121b | -3.208316 | 0.001208 |
| Ttc26 | -3.209116 | 0.030125 |
| Cd72 | -3.215785 | 1.73E-26 |
| Khdrbs3 | -3.225109403 | 9.92E-09 |
| Nlrc3 | -3.232396 | 7.60E-38 |
| Shisa8 | -3.233128 | 1.77E-09 |
| Prrg4 | -3.259247 | 7.61E-06 |
| Dtx1 | -3.26466 | 3.08E-06 |
| Cd5 | -3.269879 | 1.45E-05 |

| | | |
|---|---|---|
| Tcf7 | -3.273222635 | 1.87E-08 |
| Prrt3 | -3.273883 | 0.00361 |
| Nhsl1 | -3.288115 | 4.76E-07 |
| Sorcs2 | -3.297413 | 0.009862 |
| Src | -3.297898 | 4.44E-08 |
| Gimap8 | -3.328291 | 8.65E-25 |
| 5830418P1 | -3.331738 | 0.001945 |
| Chrnb1 | -3.334628 | 0.005532 |
| Patj | -3.339585 | 0.000508 |
| Mcoln2 | -3.353420238 | 1.24E-09 |
| Rhobtb3 | -3.357271946 | 0.001432 |
| Icos | -3.358899 | 0.004364 |
| Spns2 | -3.367799 | 8.50E-16 |
| Hes1 | -3.375198 | 0.024666 |
| Dmwd | -3.376147 | 5.49E-07 |
| Carmil2 | -3.381028 | 7.39E-51 |
| Gm10406 | -3.385341 | 0.024352 |
| Slc27a2 | -3.390714 | 0.011039 |
| Blnk | -3.403229 | 9.87E-06 |
| Nrxn1 | -3.409123 | 0.039715 |
| Timd2 | -3.411065 | 0.026776 |
| P2ry14 | -3.417528 | 1.49E-27 |
| Mei4 | -3.430193 | 0.000232 |
| Col4a2 | -3.436411 | 1.01E-05 |
| Sema7a | -3.448975 | 0.000154 |
| Syde2 | -3.462762 | 7.97E-13 |
| Itih5 | -3.488853 | 1.30E-18 |
| Cttn | -3.492352 | 9.05E-11 |
| Tpm2 | -3.503139 | 0.000952 |
| Rgs7bp | -3.504547656 | 0.03205 |
| Emp1 | -3.505416 | 0.033963 |
| Bex1 | -3.526445 | 7.66E-10 |
| Fam169b | -3.533321 | 0.013902 |
| Cyb561 | -3.534307182 | 0.00044 |
| Pear1 | -3.537471 | 6.61E-11 |
| Lat | -3.539208 | 4.14E-09 |
| Gm42372 | -3.541045 | 0.009149 |
| Tmem44 | -3.567568 | 2.14E-06 |
| Gimap5 | -3.585003 | 1.45E-05 |
| Fam171a1 | -3.593766 | 1.58E-06 |
| Mboat2 | -3.610027666 | 0.001093 |
| Blk | -3.646916075 | 4.35E-07 |
| 9030619P0 | -3.65391 | 1.98E-11 |
| Gm2237 | -3.662481 | 0.000371 |
| Il7r | -3.662962236 | 3.48E-12 |
| Rasal1 | -3.673437 | 1.51E-24 |
| Gypa | -3.682043 | 0.003921 |

| | | |
|---|---|---|
| Ablim1 | -3.724062 | 4.49E-43 |
| 6430571L1 | -3.72407 | 0.038615 |
| Lrrc66 | -3.726512 | 0.000623 |
| Pglyrp2 | -3.767501 | 6.34E-05 |
| Kcnh2 | -3.769749 | 5.74E-05 |
| Tbxa2r | -3.87011 | 8.54E-25 |
| Tek | -3.87046159 | 0.03696 |
| Sh2d2a | -3.902337 | 0.022088 |
| Xkrx | -3.916917 | 4.56E-07 |
| Myo5c | -3.929163 | 0.035969 |
| Notch3 | -3.933318 | 0.002011 |
| Havcr1 | -3.936708 | 0.001411 |
| Vmn2r29 | -3.958797 | 0.0308 |
| Slc22a3 | -3.967497 | 2.26E-07 |
| Gcnt4 | -3.979996 | 0.005039 |
| Rorc | -3.981339 | 0.049919 |
| Ddx4 | -3.991323778 | 0.001102 |
| Large2 | -3.991711 | 6.72E-05 |
| Msrb3 | -4.001162 | 1.23E-07 |
| Bank1 | -4.020229 | 0.045744 |
| 1700019D( | -4.034114 | 0.00813 |
| Gm3488 | -4.035439 | 0.037529 |
| Sox6 | -4.069186 | 0.024632 |
| Arpp21 | -4.079713 | 0.007718 |
| Nsg2 | -4.107405202 | 0.017856 |
| Srpk3 | -4.111206729 | 0.001185 |
| Mmp15 | -4.114562 | 0.0002 |
| Dok2 | -4.122130002 | 7.30E-10 |
| Tgfbr3 | -4.13382 | 0.006451 |
| Myl10 | -4.143476142 | 0.000152 |
| Spock2 | -4.147476 | 0.029918 |
| Dmd | -4.159221 | 0.007623 |
| Rab30 | -4.165289 | 0.000618 |
| Syn3 | -4.176664 | 0.002049 |
| Lck | -4.191724295 | 0.000256 |
| Gm3591 | -4.220548 | 0.024083 |
| Bpifc | -4.233575 | 5.19E-15 |
| Tox | -4.243677 | 2.70E-11 |
| Diras2 | -4.252949 | 0.035327 |
| Gm3558 | -4.258976 | 0.023184 |
| Galnt5 | -4.274697 | 6.04E-05 |
| Art4 | -4.294988 | 0.013159 |
| Hs3st1 | -4.298873 | 0.018025 |
| Cmya5 | -4.314163 | 1.78E-06 |
| Tshz3 | -4.319727814 | 0.010463 |
| Nrgn | -4.328347 | 9.60E-12 |
| Il2rb | -4.335522 | 0.001367 |

| | | |
|---|---|---|
| Klhl30 | -4.339935 | 3.29E-10 |
| Cep126 | -4.362491 | 0.000633 |
| Ctla2b | -4.363075 | 0.006793 |
| Tmtc1 | -4.383975 | 0.011486 |
| Sema4g | -4.397489 | 1.24E-05 |
| Sgsm1 | -4.402696 | 1.45E-05 |
| Zfp831 | -4.453494 | 2.93E-09 |
| Slc35d3 | -4.484751 | 7.58E-05 |
| Bcam | -4.518892154 | 0.006512 |
| Klra1 | -4.526428 | 0.045744 |
| Mycn | -4.529538 | 6.55E-17 |
| Bend5 | -4.536925 | 1.89E-18 |
| Gimap6 | -4.560529 | 4.29E-07 |
| Gem | -4.564922 | 1.12E-05 |
| Gm10409 | -4.568029 | 0.021593 |
| Igsf10 | -4.663224 | 0.000168 |
| Igf2r | -4.700046 | 6.04E-05 |
| Gimap1 | -4.706519 | 3.62E-25 |
| Gm30948 | -4.717531 | 0.000254 |
| Ctsw | -4.758449 | 1.03E-09 |
| Six5 | -4.776616 | 4.83E-05 |
| Gm3636 | -4.801862 | 0.000146 |
| Fam167a | -4.817317 | 3.34E-08 |
| Gimap7 | -4.831594 | 3.51E-11 |
| Clu | -4.838050836 | 0.001136 |
| Cachd1 | -4.882064 | 4.47E-05 |
| Nectin3 | -4.90241 | 7.20E-05 |
| Lax1 | -4.936814 | 1.72E-12 |
| Rag1 | -4.958923 | 1.07E-10 |
| Armc3 | -4.972184 | 0.013902 |
| Pkhd1l1 | -4.976338 | 0.038554 |
| Tox2 | -5.010123 | 0.001646 |
| 1500009L1 | -5.033767 | 0.010975 |
| Dntt | -5.053453 | 2.63E-16 |
| Arhgap42 | -5.071138 | 0.004635 |
| Lrriq3 | -5.090176 | 0.027564 |
| Tmem98 | -5.356021 | 0.000672 |
| Rag2 | -5.363274 | 3.31E-20 |
| Tspan8 | -5.429119 | 0.00615 |
| Igdcc4 | -5.441923 | 9.34E-05 |
| Plekha7 | -5.518519 | 3.48E-20 |
| Sytl3 | -5.545062 | 0.022959 |
| Aire | -5.570680222 | 0.001192 |
| Camkv | -5.586951 | 3.05E-06 |
| Hnf4a | -5.653532987 | 0.001815 |
| Mcam | -5.655109 | 0.034193 |
| Socs2 | -5.69631076 | 6.39E-05 |

| | | |
|---|---|---|
| Tnfsf11 | -5.698578152 | 0.0017 |
| Angpt1 | -5.701744225 | 2.23E-27 |
| Slc40a1 | -5.733114 | 5.58E-07 |
| Hmga2 | -5.783173 | 0.003382 |
| Ednra | -5.789259 | 0.021349 |
| Kif21a | -5.832226 | 0.041532 |
| Pllp | -5.840842 | 0.035874 |
| Arhgap44 | -5.848807 | 0.002846 |
| Kitl | -5.877570149 | 0.005101 |
| Ctla2a | -5.901503 | 8.17E-13 |
| Gls2 | -5.908259 | 0.001029 |
| Odf3b | -5.938687 | 0.036127 |
| Gfra4 | -6.019608 | 0.002456 |
| Gm3173 | -6.044026 | 0.042041 |
| 2010300C0 | -6.066554 | 1.29E-05 |
| 5830411N0 | -6.118251 | 0.048756 |
| Rufy4 | -6.147529 | 0.027089 |
| Dyrk4 | -6.148959 | 0.000203 |
| 2210416O1 | -6.218399928 | 0.001215 |
| Farp1 | -6.250452 | 3.77E-06 |
| Gm5111 | -6.251366 | 0.018483 |
| Gucy1a1 | -6.313595 | 0.013333 |
| Glt8d2 | -6.33133717 | 0.018592 |
| AA467197 | -6.340893 | 1.11E-13 |
| Sstr2 | -6.350973 | 3.80E-16 |
| Il17re | -6.360033 | 0.029428 |
| Tex26 | -6.40135 | 0.038615 |
| Gm10125 | -6.452951 | 0.014293 |
| Ar | -6.471354 | 1.45E-05 |
| Eps8l2 | -6.606722 | 0.00061 |
| Colq | -6.635461 | 1.45E-05 |
| Cxcr5 | -6.66045 | 0.012759 |
| Kbtbd12 | -6.6632 | 3.76E-05 |
| Boll | -6.670002 | 0.000118 |
| Zfp879 | -6.778353 | 0.029885 |
| 2900079G2 | -6.794678 | 9.10E-05 |
| Pitx2 | -6.803644 | 0.026616 |
| Fxyd1 | -6.831451 | 4.49E-11 |
| Slc24a2 | -6.841972 | 0.007923 |
| Klhl4 | -6.846076 | 0.000126 |
| Mtcl1 | -6.900336 | 0.043848 |
| Peg10 | -6.908767 | 0.006075 |
| Sox5 | -6.968878 | 0.005426 |
| Obsl1 | -6.969447 | 0.003282 |
| Mpped2 | -7.03381471 | 0.004237 |
| Bok | -7.067224 | 0.005045 |
| Wif1 | -7.092248978 | 0.00319 |

| | | |
|---|---|---|
| Fgf13 | -7.116158 | 5.83E-11 |
| Shank3 | -7.117132 | 0.043 |
| Kcnj11 | -7.141173 | 0.010445 |
| Scube3 | -7.154625 | 1.57E-11 |
| Etv4 | -7.163140316 | 0.025417 |
| Myh14 | -7.172913 | 0.034193 |
| Rai2 | -7.20837 | 0.001463 |
| Crispld1 | -7.264954 | 0.001242 |
| Cd163l1 | -7.303657 | 0.036141 |
| Ankrd33b | -7.306946828 | 9.60E-06 |
| Myog | -7.321152 | 0.000478 |
| Gata3 | -7.421924905 | 0.000556 |
| Vldlr | -7.458832 | 4.80E-08 |
| Efcc1 | -7.459605 | 5.80E-06 |
| Cald1 | -7.464368 | 0.028111 |
| Dlk2 | -7.5103 | 0.007537 |
| Pdzk1ip1 | -7.548733 | 0.000351 |
| Tspan6 | -7.549037 | 1.70E-11 |
| Mpl | -7.660428281 | 0.000184 |
| Hmgcs2 | -7.670308 | 9.47E-05 |
| St8sia2 | -7.731146 | 7.21E-05 |
| Samd4 | -7.812057329 | 0.001147 |
| Tspan9 | -7.865057 | 8.11E-06 |
| Enpp6 | -7.932124 | 0.000268 |
| Dcaf12l1 | -7.932183 | 8.68E-06 |
| Gcsam | -8.26967 | 3.96E-06 |
| Zfp57 | -8.279294 | 2.15E-06 |
| Emcn | -8.285648 | 3.43E-05 |
| Sez6l | -8.296763 | 1.45E-06 |
| Ccser1 | -8.419023 | 4.65E-07 |
| Dpf3 | -8.478793887 | 5.33E-07 |
| Pgr | -8.729507 | 2.67E-05 |
| Itga9 | -8.914568 | 0.010043 |
| Eya2 | -9.112571074 | 8.59E-09 |
| Tnni1 | -9.481857 | 0.000793 |
| Lrch2 | -9.63579 | 9.86E-11 |
| Zfp827 | -10.05616 | 9.11E-14 |
| Lpar1 | -10.1402 | 0.001819 |
| Sgce | -10.73913914 | 0.004674 |
