## Supplemental Table 2 for "Genomic Analysis of Progenitors in Viral Infection Implicates Glucocorticoids as Suppressors of Plasmacytoid Dendritic Cell Generation"

Supplementary Table 2. List of genes that are differentially expressed in $Lin^{-}c\text{-}kit^{int/lo}Flt3^{+}$ progenitors from day 8 p.i. Cl13-infected vs. uninfected mice by RNA-seq.

| symbol | log2FoldChange (Cl13 vs. Un) | FDR |
|---|---|---|
| H2-Q7 | 2.919312724 | 7.16E-198 |
| H2-Q6 | 2.727541379 | 1.23E-139 |
| H2-K1 | 1.756961615 | 1.59E-89 |
| Rab37 | -4.778282528 | 9.26E-86 |
| Gbp5 | 3.206793212 | 1.09E-84 |
| H2-D1 | 1.464876359 | 1.51E-83 |
| Gbp7 | 1.967093272 | 6.41E-79 |
| Gbp2 | 4.922232887 | 2.22E-73 |
| Ifi204 | 3.56771646 | 3.12E-73 |
| Carmil2 | -4.135960007 | 9.66E-73 |
| Ifitm3 | 2.440684443 | 2.31E-72 |
| H2-Q5 | 2.65743981 | 2.31E-68 |
| Siglech | -3.369998794 | 5.27E-68 |
| Gcnt2 | -2.477555011 | 2.67E-67 |
| Tspan2 | -2.75349898 | 2.38E-65 |
| AB124611 | 2.613839858 | 2.60E-62 |
| H2-T22 | 1.777140587 | 9.09E-61 |
| Ablim1 | -4.385441603 | 5.36E-59 |
| Mt1 | 3.238856214 | 6.75E-59 |
| F10 | 2.699542077 | 1.05E-56 |
| Gvin1 | 2.046140671 | 1.24E-53 |
| Ifi211 | 2.580996514 | 1.14E-52 |
| Scarf1 | 4.38235928 | 2.06E-52 |
| Ly6c1 | 2.434699312 | 8.78E-52 |
| Atp2b4 | -1.980899645 | 2.96E-51 |
| Rasal1 | -5.709908143 | 6.01E-51 |
| B2m | 1.242352272 | 2.48E-50 |
| Dab2ip | -1.870228086 | 6.60E-48 |
| Nlrc3 | -3.820074115 | 5.87E-47 |
| Camk1d | -2.589577311 | 3.41E-46 |
| Ly6a | 6.452332896 | 4.14E-46 |
| Dntt | -8.668269546 | 6.04E-46 |
| Gda | 3.933951783 | 8.02E-46 |
| Ms4a4c | 2.220196307 | 1.46E-45 |
| Muc13 | 1.362947953 | 2.31E-45 |
| Gm4070 | 2.048768321 | 2.97E-45 |
| Ly6d | -4.835230386 | 1.66E-44 |
| Il15ra | 2.452840027 | 9.52E-44 |
| Adam11 | -3.425970514 | 9.45E-43 |
| Axl | 2.482444203 | 1.92E-42 |
| Plekha7 | -8.577451017 | 1.02E-40 |
| Ly6c2 | 2.382195114 | 1.18E-40 |
| Emid1 | -3.985395243 | 1.18E-40 |

| | | |
|---|---|---|
| Calhm6 | 2.823625051 | 1.28E-40 |
| Adgrg1 | -3.092486002 | 1.20E-39 |
| Ctr9 | -1.444695879 | 1.94E-39 |
| H2-Q1 | 1.436340213 | 2.07E-39 |
| Gpx3 | 3.434452839 | 1.03E-38 |
| Xdh | 2.510432537 | 1.31E-38 |
| Gimap1 | -6.859358282 | 4.12E-38 |
| Fyn | -1.699606609 | 5.18E-38 |
| Gm4951 | 8.38923877 | 6.72E-38 |
| Paqr5 | -5.661330389 | 1.14E-37 |
| Chd7 | 2.659101354 | 3.59E-37 |
| Gbp3 | 1.444154804 | 3.80E-37 |
| Spon1 | 7.661955316 | 2.59E-36 |
| Sept1 | -3.074261642 | 2.61E-36 |
| Iigp1 | 5.946160859 | 7.33E-36 |
| Padi2 | -2.229393644 | 8.26E-36 |
| H2-Q4 | 1.925566885 | 8.26E-36 |
| Cnn3 | -2.830551378 | 8.70E-36 |
| Fgl2 | 2.302993035 | 2.22E-35 |
| Selenom | 3.145664205 | 2.22E-35 |
| Cd72 | -3.717114062 | 2.79E-35 |
| Tap1 | 1.361400017 | 4.11E-35 |
| Ptp4a3 | -1.564259556 | 1.24E-34 |
| Cp | 3.962827786 | 2.61E-34 |
| Ifi205 | 1.94720116 | 3.25E-34 |
| Ly6a2 | 2.92909824 | 3.50E-34 |
| Il1r1 | -3.368000599 | 5.24E-34 |
| Nod1 | 1.379569456 | 8.34E-34 |
| Myof | 2.956897689 | 1.02E-33 |
| Tle2 | -4.09918422 | 1.37E-33 |
| Itgax | -4.500171684 | 1.63E-33 |
| Serpina3i | 5.3058034 | 2.03E-33 |
| Oas1g | 3.280846937 | 8.56E-33 |
| Klk8 | -2.76910426 | 1.05E-32 |
| Plekhg5 | -3.551954065 | 4.72E-32 |
| Parp12 | 3.327237527 | 1.02E-31 |
| Tbxa2r | -5.691497971 | 1.13E-31 |
| C3 | 2.294544321 | 1.39E-31 |
| Selp | 3.257860258 | 2.39E-31 |
| Hp | 2.59926026 | 2.39E-31 |
| Ifitm1 | 2.381421502 | 1.80E-30 |
| Cmah | -5.612333603 | 2.08E-30 |
| Batf2 | 6.021121778 | 2.08E-30 |
| Net1 | -1.448928283 | 9.66E-30 |
| Htra2 | 1.051378599 | 1.07E-29 |
| Ctss | 1.331671317 | 2.29E-29 |
| H2-Ob | -3.286374068 | 3.03E-29 |

| | | |
|---|---|---|
| Hmgn1 | -1.517698293 | 3.12E-29 |
| Zbp1 | 3.837946449 | 3.77E-29 |
| Lpl | 3.020219367 | 4.06E-29 |
| Il7r | -6.288794355 | 5.38E-29 |
| Cd209a | -5.271899888 | 7.99E-29 |
| Igtp | 2.405607553 | 1.26E-28 |
| App | 1.944735174 | 1.27E-28 |
| Marcks | -4.713130623 | 1.67E-28 |
| Ripor2 | -1.173455345 | 1.77E-28 |
| Adgrg3 | -2.09282115 | 1.80E-28 |
| Nlrc5 | 1.973818335 | 1.99E-28 |
| Naip2 | 1.915220861 | 2.22E-28 |
| Rbpms | -3.866177455 | 2.89E-28 |
| Stat1 | 1.840954612 | 3.93E-28 |
| Serpina3g | 5.320183391 | 5.13E-28 |
| Egfl7 | -2.149160137 | 1.01E-27 |
| Oasl2 | 3.558498779 | 1.39E-27 |
| Syne2 | -2.650651597 | 3.54E-27 |
| Shtn1 | 3.110538617 | 4.31E-27 |
| Adcy6 | -3.377382829 | 4.69E-27 |
| Atp1b1 | -3.338199193 | 4.94E-27 |
| Ifi207 | 2.431195418 | 7.02E-27 |
| Tmem178 | 2.007551381 | 7.40E-27 |
| Angpt1 | -6.117852278 | 1.28E-26 |
| Cybb | 2.116825266 | 1.67E-26 |
| Ctsc | 2.087047264 | 2.52E-26 |
| Mgst1 | 2.383680528 | 4.34E-26 |
| Satb1 | -2.301998847 | 4.46E-26 |
| Mef2c | -1.180251025 | 4.72E-26 |
| Synpo2 | -5.903296968 | 6.61E-26 |
| Ptprf | -4.019364074 | 8.28E-26 |
| Kcng1 | -4.153009662 | 8.33E-26 |
| Apobec1 | 1.726491023 | 1.67E-25 |
| Gatm | 1.531518337 | 1.73E-25 |
| Ifitm6 | 2.12481088 | 4.39E-25 |
| Gm12185 | 4.21112433 | 4.50E-25 |
| Irf7 | 2.352195859 | 4.64E-25 |
| Epb41l4b | -2.658263093 | 4.94E-25 |
| Chst15 | -2.471872308 | 5.45E-25 |
| Oas1a | 2.520126614 | 5.50E-25 |
| Tnfrsf13c | -3.31780801 | 6.89E-25 |
| Aldoa | 1.027120628 | 6.95E-25 |
| Nrgn | -7.463076275 | 7.34E-25 |
| Mn1 | -6.502533097 | 7.47E-25 |
| Clec5a | 1.799781593 | 9.98E-25 |
| Snn | -3.69320651 | 1.34E-24 |
| Ly6i | 4.423330238 | 1.58E-24 |

| | | |
|---|---|---|
| Fgr | -3.334904482 | 3.36E-24 |
| Tmem106a | 2.130113542 | 3.68E-24 |
| Lap3 | 1.452424437 | 4.41E-24 |
| Samhd1 | 1.30315879 | 4.42E-24 |
| Kalrn | 1.697404435 | 5.77E-24 |
| Pear1 | -5.299268364 | 8.25E-24 |
| Ifngr1 | 1.463347987 | 1.24E-23 |
| Cd96 | -5.639822104 | 1.73E-23 |
| Prtn3 | 1.418354479 | 2.11E-23 |
| Alas1 | 1.345517013 | 2.67E-23 |
| Cxxc5 | -2.693641894 | 2.90E-23 |
| Myl4 | -3.148560284 | 4.47E-23 |
| Trpm2 | 2.142647173 | 6.64E-23 |
| Tmem38b | 2.016203679 | 7.60E-23 |
| Mllt6 | -1.359682103 | 8.03E-23 |
| Cfp | -1.324917438 | 8.40E-23 |
| Fcgr1 | 3.359999509 | 9.80E-23 |
| Kmo | -2.087344215 | 9.80E-23 |
| Cd34 | -2.044178995 | 9.81E-23 |
| Plekha5 | -1.048011325 | 1.00E-22 |
| Srgap3 | -2.537535014 | 1.02E-22 |
| Ifi209 | 1.024647168 | 1.06E-22 |
| 2700081O1 | -2.181119463 | 1.09E-22 |
| Marcksl1 | -2.604953114 | 1.36E-22 |
| Acss1 | -2.432575288 | 1.39E-22 |
| Spata13 | 1.425069153 | 1.52E-22 |
| Casp4 | 1.913845272 | 1.73E-22 |
| B4galt6 | 1.986486171 | 2.83E-22 |
| Atf5 | 1.814300732 | 3.49E-22 |
| Gstm1 | 2.488553651 | 4.10E-22 |
| Fads3 | -2.96721406 | 5.51E-22 |
| Cd68 | 1.031952919 | 5.66E-22 |
| C4a | 3.393344237 | 6.61E-22 |
| Plek | -1.900492835 | 7.86E-22 |
| Pik3ip1 | 1.261512058 | 8.02E-22 |
| Gm7030 | 1.247003019 | 8.02E-22 |
| Il12rb1 | 3.039377544 | 8.76E-22 |
| 1600014C1 | 1.540316056 | 1.19E-21 |
| Rtn4rl1 | -3.377130736 | 1.53E-21 |
| Btla | 1.353864936 | 1.61E-21 |
| Lamp2 | 1.040463954 | 1.68E-21 |
| Mex3b | -2.783389326 | 2.48E-21 |
| Nrp2 | 1.465107065 | 2.66E-21 |
| Gpnmb | 5.800031718 | 3.11E-21 |
| H2-Q10 | 1.948599654 | 3.47E-21 |
| Ms4a6b | 1.033658854 | 4.51E-21 |
| 2610035D1 | -2.940568429 | 4.53E-21 |

| | | |
|---|---|---|
| Arl4c | -2.95683868 | 4.67E-21 |
| Aoah | 2.002875191 | 5.44E-21 |
| Il17rb | -3.076359331 | 7.32E-21 |
| Runx2 | -2.167876516 | 8.77E-21 |
| Gm8909 | 1.357973477 | 8.86E-21 |
| Cd27 | -1.879138401 | 9.04E-21 |
| Cobll1 | -3.317156662 | 9.48E-21 |
| P2ry14 | -2.933208659 | 1.59E-20 |
| Il12a | -2.15854082 | 2.38E-20 |
| Lax1 | -6.991696385 | 3.63E-20 |
| Prkca | -4.568416768 | 3.97E-20 |
| Ifi47 | 1.772380908 | 4.04E-20 |
| Aff3 | -4.198998869 | 4.54E-20 |
| Slc31a2 | 1.466205829 | 4.54E-20 |
| Tmem119 | -2.698533844 | 4.86E-20 |
| Entpd4b | -2.977984561 | 6.21E-20 |
| Ifi214 | 1.02098042 | 6.85E-20 |
| Lat | -5.801075397 | 7.13E-20 |
| 9030619P0 | -5.890317251 | 8.58E-20 |
| Itsn1 | -1.391397971 | 9.98E-20 |
| Gsn | -1.966053965 | 1.04E-19 |
| Fcgr3 | 2.424635465 | 1.25E-19 |
| Rag2 | -7.747804773 | 1.53E-19 |
| Blnk | -6.858077043 | 1.56E-19 |
| Ctsg | 1.192136493 | 1.74E-19 |
| Gbp10 | 3.222161735 | 1.77E-19 |
| Zfp608 | -2.057787095 | 1.87E-19 |
| Pdzd4 | -2.603811807 | 1.89E-19 |
| Pygl | 1.340593488 | 1.91E-19 |
| Apcdd1 | -3.45095787 | 2.02E-19 |
| Gm18852 | 3.047330417 | 2.45E-19 |
| Parp14 | 1.292925416 | 3.82E-19 |
| Sytl1 | -3.236796158 | 3.95E-19 |
| Uhrf1bp1 | 1.296811308 | 4.01E-19 |
| Wdr6 | -1.149915145 | 4.48E-19 |
| Entpd4 | -3.059347353 | 5.09E-19 |
| L1cam | -2.149787285 | 5.13E-19 |
| N4bp3 | -2.844417151 | 8.53E-19 |
| Serpina3f | 6.902829535 | 9.33E-19 |
| Trem3 | 2.752094649 | 1.05E-18 |
| Ifi44 | 2.414033885 | 1.10E-18 |
| Hopx | 2.644312913 | 1.14E-18 |
| Phf11d | 4.070878882 | 1.32E-18 |
| Nrg1 | 1.648561629 | 1.36E-18 |
| F830016B0 | 6.122439855 | 1.75E-18 |
| Lgals3bp | 2.03413478 | 2.27E-18 |
| Gm18853 | 2.986645388 | 2.66E-18 |

| | | |
|---|---|---|
| Sema4c | -2.380998592 | 3.55E-18 |
| Ndrg2 | 4.444493045 | 3.94E-18 |
| Hdac7 | -1.027090498 | 3.96E-18 |
| Mtus1 | 1.25079048 | 4.10E-18 |
| Faah | -3.762522832 | 4.24E-18 |
| Slc4a8 | 1.666948745 | 5.39E-18 |
| Ets1 | -5.278359814 | 7.05E-18 |
| Cd6 | -2.741484799 | 7.65E-18 |
| Bach2 | -2.67448753 | 9.67E-18 |
| Ccr9 | -3.491353761 | 1.07E-17 |
| Gimap8 | -10.93944619 | 1.12E-17 |
| Dstn | 1.999791083 | 1.44E-17 |
| Evpl | -3.038183085 | 1.56E-17 |
| Oas2 | 2.675188236 | 1.79E-17 |
| Rpgrip1 | -1.26042179 | 1.91E-17 |
| C4b | 7.151088639 | 2.39E-17 |
| Dbp | -1.835526607 | 2.46E-17 |
| Mctp2 | -3.866148254 | 2.79E-17 |
| Gpr141 | 1.983551933 | 2.85E-17 |
| Amotl2 | 10.82163388 | 2.91E-17 |
| Zfp422 | -1.089689009 | 5.16E-17 |
| Vamp4 | 1.015219444 | 6.83E-17 |
| Gsr | 1.854359425 | 6.90E-17 |
| Bpifc | -6.55637428 | 6.91E-17 |
| H2-M3 | 1.238903865 | 7.11E-17 |
| Ifi213 | 5.474831836 | 8.09E-17 |
| Oasl1 | 3.140470226 | 8.59E-17 |
| Cxcl10 | 3.578038897 | 8.94E-17 |
| Camk2d | 1.682258032 | 1.13E-16 |
| Khk | 1.373168203 | 1.17E-16 |
| H2-T10 | 1.523943653 | 1.18E-16 |
| Abca1 | -3.744754502 | 1.27E-16 |
| Tgtp1 | 2.446964245 | 1.30E-16 |
| Plin2 | 1.697956672 | 1.61E-16 |
| Tgtp2 | 2.090226278 | 2.16E-16 |
| Cd22 | -2.548048936 | 2.28E-16 |
| Ppp1r16b | -2.784804024 | 2.96E-16 |
| Lbp | 1.458548872 | 3.00E-16 |
| Kif23 | -1.345648092 | 3.18E-16 |
| Tfec | 1.430880784 | 3.82E-16 |
| Papss2 | 1.419633954 | 3.91E-16 |
| Tex2 | -1.110541663 | 4.73E-16 |
| Dusp5 | -2.433736767 | 5.58E-16 |
| Upb1 | -2.369691365 | 6.35E-16 |
| Eps8 | 1.997574674 | 6.56E-16 |
| Irgm2 | 2.056216498 | 7.31E-16 |
| Clec7a | 1.12165503 | 7.34E-16 |

| | | |
|---|---|---|
| Ctnnd2 | -4.44175391 | 7.51E-16 |
| Gimap9 | -4.346228717 | 1.01E-15 |
| Neu1 | 1.287021462 | 1.03E-15 |
| Tanc1 | -3.517102912 | 1.03E-15 |
| Tcf7 | -4.902007436 | 1.11E-15 |
| Abca13 | 4.933372605 | 1.11E-15 |
| Sesn1 | -1.181651098 | 1.20E-15 |
| Zscan2 | -2.277181962 | 1.28E-15 |
| Sema6b | 1.622463211 | 1.33E-15 |
| Spns2 | -3.383875749 | 1.39E-15 |
| Tapbpl | 1.105026551 | 1.54E-15 |
| Ugt1a7c | 1.381341748 | 1.58E-15 |
| Mrvi1 | -9.218763518 | 1.70E-15 |
| Dclk2 | -2.908899006 | 1.94E-15 |
| Parp10 | 1.536962443 | 2.01E-15 |
| Tctex1d1 | 2.393099214 | 2.14E-15 |
| Glis2 | -3.329700219 | 2.35E-15 |
| Grap | 1.895963728 | 2.62E-15 |
| Soat1 | 1.297313875 | 3.07E-15 |
| Bcl2 | -1.837811414 | 3.07E-15 |
| Ripor1 | -1.342452791 | 3.12E-15 |
| Adgrb1 | 12.0476952 | 3.15E-15 |
| Bcr | -1.050371086 | 3.71E-15 |
| Gm11127 | 1.23610763 | 3.75E-15 |
| Psmb10 | 1.001350949 | 4.49E-15 |
| Blk | -6.641412175 | 4.61E-15 |
| Bmx | 2.175438359 | 5.00E-15 |
| Ctxn1 | -3.558910634 | 5.17E-15 |
| Drc7 | -3.062802237 | 6.58E-15 |
| Ttc3 | -1.036760232 | 6.71E-15 |
| Nedd4 | -4.167978793 | 6.95E-15 |
| H6pd | 1.07003072 | 7.77E-15 |
| Msrb1 | 1.412003084 | 7.77E-15 |
| Ikbke | 1.255007434 | 7.81E-15 |
| AI504432 | -1.721676287 | 7.99E-15 |
| Xaf1 | 2.142109052 | 8.11E-15 |
| Ddah2 | -2.445457104 | 9.42E-15 |
| Atp11a | 2.093751453 | 9.52E-15 |
| Med12l | -3.595962207 | 9.52E-15 |
| Gm807 | 3.036726426 | 9.52E-15 |
| 0610009E0 | -1.222511195 | 1.02E-14 |
| Arhgef40 | -2.148065747 | 1.10E-14 |
| Sh2d5 | -1.560125321 | 1.32E-14 |
| Igsf6 | 1.146868763 | 1.68E-14 |
| Tcf4 | -1.166767793 | 1.72E-14 |
| Nav2 | 1.412982112 | 1.87E-14 |
| Tox | -5.381761147 | 1.95E-14 |

| | | |
|---|---|---|
| BB031773 | 2.662594825 | 2.20E-14 |
| Myl10 | -8.230157848 | 2.63E-14 |
| Ms4a3 | 3.301465967 | 2.63E-14 |
| Peg13 | -2.397162624 | 2.64E-14 |
| Cldn15 | 2.18477154 | 3.07E-14 |
| H2-Ab1 | 2.075394255 | 3.19E-14 |
| Fads2 | -2.686110684 | 3.22E-14 |
| Zdhhc8 | -1.316789533 | 3.34E-14 |
| Epha2 | -3.035126307 | 3.34E-14 |
| Flt3 | -1.22458552 | 3.44E-14 |
| Hpgd | 3.292155476 | 3.74E-14 |
| Bcl9l | -1.759024776 | 3.74E-14 |
| Rhou | 4.915152165 | 3.87E-14 |
| Abcc4 | 1.115258593 | 4.07E-14 |
| Inka1 | -1.746257068 | 4.07E-14 |
| Tspan13 | -2.740113682 | 4.20E-14 |
| Slfn2 | 1.921338048 | 4.27E-14 |
| Zeb1 | -2.306161629 | 5.42E-14 |
| Ostm1 | 1.214898641 | 6.72E-14 |
| Trp53i11 | -2.14361076 | 6.90E-14 |
| Phc1 | -1.112672079 | 7.61E-14 |
| Bcl11a | -1.760171845 | 9.38E-14 |
| Hes6 | -1.206380765 | 9.57E-14 |
| Tut4 | -1.022134124 | 1.00E-13 |
| Clcf1 | -2.811124546 | 1.29E-13 |
| Zfp2 | -3.713255929 | 1.36E-13 |
| Prkch | -1.980496067 | 1.84E-13 |
| Hdac10 | -1.145946142 | 2.08E-13 |
| Agrn | 1.412559076 | 2.14E-13 |
| Trim45 | 1.29096564 | 2.27E-13 |
| Klrk1 | 1.469460188 | 2.40E-13 |
| Basp1 | -2.975852662 | 2.44E-13 |
| Apbb1 | -8.314247622 | 2.48E-13 |
| Tbc1d10c | -1.689380906 | 2.56E-13 |
| Atp6v0a1 | -1.583041944 | 3.00E-13 |
| Rag1 | -8.290819292 | 3.07E-13 |
| Gbp6 | 3.239238705 | 3.29E-13 |
| Arpp21 | -13.17753597 | 3.37E-13 |
| Spib | -3.173568738 | 3.59E-13 |
| Gramd3 | 1.690945919 | 3.87E-13 |
| Fam171a2 | -5.6054065 | 4.68E-13 |
| Ifi27l2a | 2.090485479 | 4.74E-13 |
| Mycn | -7.239848252 | 5.21E-13 |
| Ugt1a6a | 1.410595115 | 5.75E-13 |
| Gm4841 | 4.486035013 | 6.23E-13 |
| Gimap6 | -6.383490845 | 6.36E-13 |
| Tsc22d1 | -1.573195641 | 6.58E-13 |

| | | |
|---|---|---|
| Cd74 | 1.988748147 | 6.58E-13 |
| Fam241a | 1.558351833 | 6.58E-13 |
| Lpcat2 | -3.060745556 | 7.33E-13 |
| Notch1 | -1.103903544 | 7.92E-13 |
| Cd28 | -3.137290205 | 8.53E-13 |
| Aldh3b1 | 1.316652046 | 8.89E-13 |
| Shisa8 | -3.893353998 | 9.29E-13 |
| Mier3 | -1.291329421 | 9.46E-13 |
| Nfil3 | -1.01702028 | 9.52E-13 |
| AA467197 | -5.885691421 | 9.82E-13 |
| Sstr2 | -8.963978124 | 1.07E-12 |
| Dtx3l | 1.053401786 | 1.37E-12 |
| Tet1 | -2.601733131 | 1.39E-12 |
| Depp1 | -4.301573489 | 1.66E-12 |
| Sdc4 | -4.218022006 | 1.76E-12 |
| Pdgfrb | -2.470473997 | 1.95E-12 |
| Smox | -1.70031046 | 2.05E-12 |
| Spns3 | -1.20267307 | 2.28E-12 |
| Ptprcap | -2.568439418 | 2.29E-12 |
| St6galnac3 | -5.183101397 | 2.32E-12 |
| Clec4a2 | 4.437726979 | 2.41E-12 |
| Slfn5 | 2.650971563 | 2.47E-12 |
| Mast4 | -3.969688904 | 2.49E-12 |
| Polm | 1.67079223 | 2.80E-12 |
| Phf11b | 2.149042256 | 3.01E-12 |
| Rnf217 | 1.838877867 | 3.25E-12 |
| Gpr171 | -1.502022516 | 3.31E-12 |
| Kansl1l | -1.298155366 | 3.32E-12 |
| Ugt1a6b | 1.346591422 | 3.36E-12 |
| Gata2 | -3.717377834 | 3.48E-12 |
| Itgad | -3.951236914 | 3.56E-12 |
| Id1 | 2.19995903 | 3.74E-12 |
| Gca | 2.46176239 | 4.13E-12 |
| Pde7a | -1.431413946 | 4.14E-12 |
| Zbtb7b | 1.038106889 | 4.15E-12 |
| Kif17 | -1.750868275 | 4.47E-12 |
| Emilin2 | 1.180152357 | 4.82E-12 |
| Ptprr | 3.640916406 | 5.06E-12 |
| Rnf213 | 1.64578259 | 5.09E-12 |
| Zfp521 | -2.085336313 | 5.10E-12 |
| Pdcd4 | 1.010037828 | 6.06E-12 |
| Mapk13 | 3.309618802 | 6.16E-12 |
| Stx11 | 1.212771882 | 6.55E-12 |
| Itih5 | -2.705717464 | 6.71E-12 |
| Gimap7 | -7.428340236 | 8.31E-12 |
| Maml3 | -1.745391043 | 8.41E-12 |
| Anxa9 | 1.491067787 | 8.63E-12 |

| | | |
|---|---|---|
| Bin1 | -1.279559053 | 8.75E-12 |
| Zcchc18 | -4.377402568 | 8.75E-12 |
| Cyp4f18 | 1.284955991 | 8.79E-12 |
| Znfx1 | 1.043102836 | 8.79E-12 |
| A530040E1 | 1.829339883 | 9.17E-12 |
| Ahdc1 | -2.049573528 | 1.00E-11 |
| Epsti1 | 1.054275887 | 1.13E-11 |
| Bfsp2 | -2.848897344 | 1.13E-11 |
| Gm16867 | -1.477215227 | 1.17E-11 |
| Acacb | 2.424527461 | 1.33E-11 |
| Adgre5 | 1.924923411 | 1.34E-11 |
| Aif1 | 1.102721708 | 1.35E-11 |
| Rnase4 | 1.793641865 | 1.48E-11 |
| Ctla2a | -6.671888343 | 1.60E-11 |
| Gm21188 | 2.497359403 | 1.71E-11 |
| Gab1 | -1.662174007 | 1.75E-11 |
| Abcg1 | -2.094874359 | 1.91E-11 |
| Glrx | 1.381452126 | 1.95E-11 |
| Msi2 | -1.638751288 | 2.03E-11 |
| Zap70 | -5.624948148 | 2.04E-11 |
| 1700025G( | -1.283734358 | 2.06E-11 |
| Pstpip2 | 1.748084419 | 2.19E-11 |
| C530008M | -5.24179729 | 2.21E-11 |
| Dmxl2 | 1.640449204 | 2.33E-11 |
| Nampt | 1.431545739 | 2.38E-11 |
| Nhsl1 | -4.316329556 | 2.41E-11 |
| Robo4 | -9.243075293 | 2.47E-11 |
| Tnik | -11.07582871 | 2.55E-11 |
| Ppfia4 | -1.017687537 | 2.58E-11 |
| Irgm1 | 1.43097944 | 2.78E-11 |
| Glul | 1.20684597 | 2.87E-11 |
| Cebpb | 2.092413934 | 3.04E-11 |
| Pnp2 | 1.181775071 | 3.12E-11 |
| Gyg | 1.168287853 | 3.20E-11 |
| Glt1d1 | 3.642121396 | 3.26E-11 |
| St6gal1 | -6.458890573 | 3.66E-11 |
| Jakmip1 | -3.621534461 | 3.71E-11 |
| Ugt1a1 | 1.334808203 | 3.94E-11 |
| Ugt1a5 | 1.334474127 | 4.04E-11 |
| Ptger4 | -1.977653075 | 4.06E-11 |
| Ugt1a2 | 1.335338767 | 4.35E-11 |
| Tchh | 3.274035042 | 4.73E-11 |
| Ugt1a9 | 1.324034718 | 5.17E-11 |
| 2900026A( | -2.386370993 | 5.20E-11 |
| Mefv | 1.039729928 | 5.71E-11 |
| Ccr7 | -4.02848321 | 5.71E-11 |
| Strbp | -1.020706852 | 5.91E-11 |

| | | |
|---|---|---|
| Thsd1 | -2.135658762 | 5.91E-11 |
| Dnmt3l | 9.312011015 | 6.06E-11 |
| Scai | -1.444719651 | 6.45E-11 |
| Tspan6 | -9.235324817 | 6.52E-11 |
| Lpin1 | -1.744554426 | 6.71E-11 |
| Bend5 | -8.656334846 | 6.76E-11 |
| Camkv | -11.00321975 | 7.23E-11 |
| Aldh1b1 | 1.086652599 | 7.29E-11 |
| Rhob | 1.051881504 | 7.33E-11 |
| Fxyd1 | -9.279335028 | 7.71E-11 |
| Lgmn | -1.300469154 | 7.77E-11 |
| Stox2 | 1.493595649 | 7.79E-11 |
| Lair1 | 1.357675997 | 8.00E-11 |
| H2-Eb1 | 2.449220061 | 8.60E-11 |
| Plaur | 1.184654563 | 9.15E-11 |
| Tes | -1.70337669 | 9.47E-11 |
| Retreg1 | -1.492641392 | 9.53E-11 |
| Scube3 | -6.047916834 | 9.79E-11 |
| Gprc5b | -3.486040534 | 1.06E-10 |
| Ms4a6d | 2.749668674 | 1.06E-10 |
| Rhoh | -1.041735534 | 1.07E-10 |
| Chpt1 | 1.946587864 | 1.14E-10 |
| Dmwd | -4.783271071 | 1.22E-10 |
| Deptor | 1.372461285 | 1.22E-10 |
| Klrd1 | -3.465497765 | 1.50E-10 |
| Fam167a | -6.715062416 | 1.57E-10 |
| Trp53i13 | -1.339871187 | 1.73E-10 |
| Csf1 | 1.401770513 | 1.81E-10 |
| Gm21188 | 2.646079259 | 1.85E-10 |
| Maged1 | -2.889413271 | 1.94E-10 |
| Rgs12 | -1.27096779 | 2.01E-10 |
| Ppargc1b | 1.284405912 | 2.40E-10 |
| Zfp827 | -3.259342917 | 2.44E-10 |
| Ccm2l | 2.614158542 | 2.57E-10 |
| Btbd3 | -1.796663603 | 2.68E-10 |
| Cd244a | -1.482079878 | 2.85E-10 |
| Elane | 2.416523257 | 3.20E-10 |
| Cd40 | 1.95135782 | 3.23E-10 |
| Pafah1b3 | -1.36374912 | 3.48E-10 |
| Dhx58 | 1.741914641 | 3.48E-10 |
| Pcbp3 | -1.075761422 | 3.53E-10 |
| Cd163 | -2.094746342 | 3.53E-10 |
| H2-Aa | 2.047510768 | 3.55E-10 |
| Ugt1a10 | 1.377683361 | 3.75E-10 |
| Slc14a1 | -2.004599368 | 4.28E-10 |
| Lrba | -1.306553289 | 4.73E-10 |
| Mcf2l | 1.207908612 | 5.12E-10 |

| | | |
|---|---|---|
| Dnajc12 | 1.511940559 | 5.18E-10 |
| Xkrx | -5.161594602 | 5.27E-10 |
| Gsdme | -1.002119231 | 5.62E-10 |
| Lrch2 | -6.431875729 | 5.65E-10 |
| Aatk | 1.405863624 | 5.85E-10 |
| Cx3cr1 | -1.624846605 | 6.08E-10 |
| Slc35d3 | -7.002939814 | 6.70E-10 |
| Zfp831 | -6.247609857 | 6.86E-10 |
| Vangl2 | -2.537514369 | 6.92E-10 |
| Cttn | -3.392457514 | 7.74E-10 |
| Chst11 | -1.411501052 | 7.92E-10 |
| Cd300lf | 2.240775454 | 9.08E-10 |
| Asns | 2.378299921 | 9.38E-10 |
| Coro2b | -5.579688501 | 9.92E-10 |
| Sectm1a | 6.105610038 | 9.92E-10 |
| Kif1b | 1.154384319 | 1.19E-09 |
| Cd7 | -1.931101082 | 1.22E-09 |
| Gm7694 | -1.366778499 | 1.28E-09 |
| Mcoln2 | -3.425452099 | 1.36E-09 |
| Il27ra | -3.175244622 | 1.40E-09 |
| Pid1 | -2.494725431 | 1.41E-09 |
| Fam184a | -6.660191581 | 1.50E-09 |
| Il31ra | -1.130690363 | 1.52E-09 |
| Mcemp1 | 2.185051291 | 1.66E-09 |
| Dio2 | 2.214354758 | 1.98E-09 |
| Abca9 | -1.781931644 | 1.98E-09 |
| Vopp1 | -1.264736394 | 2.11E-09 |
| Siglece | 1.481089958 | 2.21E-09 |
| Mx1 | 1.255380328 | 2.25E-09 |
| Fgf13 | -8.114793212 | 2.26E-09 |
| Meis1 | -1.17529206 | 2.31E-09 |
| Tgfbi | 1.785245751 | 2.39E-09 |
| Pitpnm1 | 1.000579572 | 2.49E-09 |
| Kcnh2 | -8.955631803 | 2.52E-09 |
| Fmnl2 | -1.141151958 | 2.59E-09 |
| Ptgr1 | 1.009515597 | 2.60E-09 |
| Irf4 | -2.989546958 | 2.81E-09 |
| Cass4 | -3.748514738 | 2.98E-09 |
| Mxd3 | -1.239118381 | 3.11E-09 |
| Saa3 | 4.340218133 | 3.26E-09 |
| Slc7a7 | 1.212091149 | 3.52E-09 |
| Arhgap24 | 1.776007127 | 3.58E-09 |
| Ccl19 | 4.280215247 | 3.59E-09 |
| Klhl30 | -8.327695812 | 3.63E-09 |
| Tnfsf13 | 1.618790753 | 3.65E-09 |
| Natd1 | -1.230285267 | 3.69E-09 |
| Sgsm1 | -8.874474087 | 3.79E-09 |

| | | |
|---|---|---|
| Rnf141 | -1.285921981 | 3.84E-09 |
| Csf2ra | 1.264560307 | 4.00E-09 |
| Ephb2 | -1.943865431 | 4.81E-09 |
| Slc25a23 | -1.90928115 | 4.95E-09 |
| Palld | -2.784027142 | 5.09E-09 |
| Kcna3 | -1.749850029 | 5.17E-09 |
| Stum | -8.874782269 | 5.66E-09 |
| Rab30 | -9.426260083 | 5.69E-09 |
| Adgra2 | -1.669550784 | 5.87E-09 |
| Gadd45b | 1.381961793 | 5.98E-09 |
| Flnb | -1.402092941 | 6.27E-09 |
| Nos1ap | -1.292266825 | 6.37E-09 |
| Epb41l3 | -3.458113632 | 6.47E-09 |
| Naaa | 1.457875037 | 6.54E-09 |
| Sorbs3 | -3.069405197 | 6.83E-09 |
| Tfrc | -1.257844129 | 7.15E-09 |
| Magee2 | -4.604464793 | 7.83E-09 |
| Sdc1 | -2.910550703 | 8.39E-09 |
| Wnt10b | -6.07699699 | 8.46E-09 |
| Ctsw | -4.440613087 | 8.68E-09 |
| Mex3a | -2.991038919 | 8.91E-09 |
| Upk1b | -3.12089838 | 9.02E-09 |
| Ccdc8 | -3.211535648 | 9.12E-09 |
| Slc35f2 | -3.819615594 | 9.15E-09 |
| Msrb3 | -4.863673877 | 9.45E-09 |
| Tef | -1.078617888 | 9.63E-09 |
| Adam2 | -5.188744273 | 9.87E-09 |
| Lsp1 | -1.048271515 | 1.04E-08 |
| Gimap5 | -8.282932858 | 1.11E-08 |
| Abhd11 | 1.007664463 | 1.16E-08 |
| Cbx2 | -1.738769444 | 1.17E-08 |
| Stk17b | -1.260500882 | 1.21E-08 |
| Ldlrad4 | -2.20737278 | 1.21E-08 |
| Plxna1 | 2.102301478 | 1.25E-08 |
| Gpc1 | 1.041377818 | 1.37E-08 |
| Hsh2d | 1.4108924 | 1.61E-08 |
| Zdhhc15 | -1.937793684 | 1.62E-08 |
| Gm16897 | -2.0730871 | 1.71E-08 |
| H2-Bl | 6.923287496 | 1.74E-08 |
| Etv5 | -2.274248456 | 1.80E-08 |
| Eya2 | -8.843429462 | 1.80E-08 |
| Dbn1 | -2.397781288 | 1.91E-08 |
| C1rl | 1.417940334 | 1.99E-08 |
| Ifih1 | 1.399948923 | 2.11E-08 |
| Heg1 | -1.326036952 | 2.13E-08 |
| Bmp1 | -2.822040516 | 2.23E-08 |
| Rgs18 | -1.01470545 | 2.29E-08 |

| | | |
|---|---|---|
| Pld1 | 1.588807628 | 2.34E-08 |
| F630028O1 | 1.250332032 | 2.34E-08 |
| Mgst2 | 1.659552409 | 2.36E-08 |
| Usp18 | 2.150390004 | 2.36E-08 |
| Adora2a | -3.927991994 | 2.43E-08 |
| Slc22a3 | -8.097163688 | 2.48E-08 |
| Cox6a2 | -2.717319847 | 2.50E-08 |
| Plpp5 | 1.060074887 | 2.70E-08 |
| Abcd2 | 1.152329942 | 2.89E-08 |
| Gpr157 | 1.892671272 | 2.94E-08 |
| Maged2 | -1.471921984 | 2.98E-08 |
| Gpsm1 | -1.497910976 | 3.24E-08 |
| Whrn | -1.890348896 | 3.24E-08 |
| Padi4 | 1.281756067 | 3.35E-08 |
| Calcoco1 | -1.581065278 | 3.68E-08 |
| Lockd | -1.134075398 | 3.77E-08 |
| Ttll9 | 1.342429543 | 3.95E-08 |
| Acap1 | -1.130896802 | 4.07E-08 |
| St3gal5 | -4.239518193 | 4.11E-08 |
| Dok2 | -3.670473078 | 4.17E-08 |
| Cmya5 | -5.249918521 | 4.17E-08 |
| Hmces | -1.283538806 | 4.46E-08 |
| Slfn1 | 4.547621096 | 4.46E-08 |
| Isg15 | 2.811285676 | 4.48E-08 |
| Il18bp | 3.099487396 | 4.58E-08 |
| Olfr56 | 2.390350405 | 4.71E-08 |
| Nfkbie | -1.406251334 | 4.90E-08 |
| Zcwpw1 | -1.625237935 | 4.97E-08 |
| Dennd2d | 1.429529571 | 5.03E-08 |
| F5 | 2.007811836 | 5.38E-08 |
| Msantd3 | 1.252456527 | 6.37E-08 |
| Slc27a1 | -1.110291109 | 6.39E-08 |
| Nlrp1b | 2.195439981 | 6.61E-08 |
| Muc3a | -3.084492793 | 6.67E-08 |
| Fut7 | -1.146988582 | 6.86E-08 |
| Homer1 | 1.223228693 | 7.14E-08 |
| Tapt1 | -1.290853796 | 7.44E-08 |
| Mroh2a | -1.405675654 | 8.09E-08 |
| Cd5 | -4.563750055 | 8.10E-08 |
| Flot2 | 1.004502526 | 8.28E-08 |
| Rtl5 | -1.562556249 | 8.90E-08 |
| Mmp15 | -7.757505287 | 9.20E-08 |
| Phldb1 | -1.354625129 | 9.85E-08 |
| Tpst1 | 1.135334608 | 9.93E-08 |
| Trpv4 | -1.655024471 | 1.02E-07 |
| Tjp2 | -1.168673798 | 1.11E-07 |
| Rell1 | -1.349285611 | 1.13E-07 |

| | | |
|---|---|---|
| Hsd17b14 | 8.105122858 | 1.15E-07 |
| Nxn | 1.132589603 | 1.20E-07 |
| Vegfc | -3.672331954 | 1.21E-07 |
| Acot1 | 1.402722765 | 1.30E-07 |
| Ano8 | -1.265313464 | 1.30E-07 |
| Ifit2 | 1.088578582 | 1.36E-07 |
| Gstm4 | 1.302406987 | 1.39E-07 |
| Card11 | -1.30666439 | 1.42E-07 |
| Scnn1a | 1.47396108 | 1.46E-07 |
| Tmem63a | 1.056970239 | 1.50E-07 |
| Adamts10 | -1.086357098 | 1.53E-07 |
| Otud7b | -1.861383086 | 1.56E-07 |
| Clec4e | 2.089084813 | 1.57E-07 |
| Nxpe4 | 1.821483802 | 1.59E-07 |
| Peak1 | -1.525130283 | 1.63E-07 |
| Lck | -5.680827286 | 1.71E-07 |
| Plaat3 | 1.374938794 | 1.73E-07 |
| Tnni2 | -1.326537581 | 1.74E-07 |
| Sh3pxd2a | -1.004793519 | 1.82E-07 |
| Ppp1r13b | -1.05664171 | 1.82E-07 |
| Eif2ak2 | 1.046712323 | 1.92E-07 |
| Tnfsfm13 | 1.105522958 | 1.96E-07 |
| H1f10 | -2.342757851 | 2.22E-07 |
| Chst3 | -4.839512813 | 2.43E-07 |
| Plcb4 | -1.087660589 | 2.49E-07 |
| Nacc2 | 1.764413508 | 2.50E-07 |
| Dzip1 | -7.487386359 | 2.52E-07 |
| Ydjc | 1.063311412 | 2.69E-07 |
| Gask1b | -5.550528971 | 2.75E-07 |
| Havcr2 | -2.75485927 | 2.86E-07 |
| Repin1 | -1.03617444 | 2.87E-07 |
| Gng11 | 2.306472933 | 2.88E-07 |
| S100a4 | -1.082970997 | 2.90E-07 |
| C1ra | 2.074374997 | 2.93E-07 |
| Dtx1 | -7.379215249 | 2.97E-07 |
| E330009J0 | 1.245303622 | 3.00E-07 |
| Pklr | -4.655981792 | 3.00E-07 |
| Cd200r4 | 7.264137888 | 3.03E-07 |
| Il2rg | 1.200559513 | 3.10E-07 |
| H2-K2 | 1.089411368 | 3.14E-07 |
| Baiap3 | -1.970538704 | 3.20E-07 |
| Icos | -8.449223347 | 3.26E-07 |
| Pygm | -2.773306701 | 3.32E-07 |
| Ralgps1 | -1.236077271 | 3.41E-07 |
| Ctsf | -1.918884263 | 3.41E-07 |
| Cd302 | 1.622721679 | 3.46E-07 |
| Gstm3 | 2.546900432 | 3.65E-07 |

| | | |
|---|---|---|
| P2ry13 | -3.678437966 | 3.85E-07 |
| C1qtnf6 | 1.452219962 | 3.88E-07 |
| F2r | -3.781765703 | 3.89E-07 |
| Tdrkh | -1.504410659 | 3.90E-07 |
| Khdrbs3 | -2.846185176 | 4.07E-07 |
| Slc22a23 | -3.545699211 | 4.09E-07 |
| Prr36 | -3.96581827 | 4.20E-07 |
| Trim30a | 1.01107218 | 4.20E-07 |
| Fbxw17 | 1.107252026 | 4.40E-07 |
| Vldlr | -5.568583155 | 4.77E-07 |
| Zfp184 | -1.046614892 | 4.93E-07 |
| Mtss1 | -1.345546634 | 5.07E-07 |
| Plcg1 | -1.190335492 | 5.21E-07 |
| Hdac9 | -1.085844826 | 5.38E-07 |
| Lcn2 | 5.498415806 | 5.40E-07 |
| Ube2l6 | 1.494871848 | 5.56E-07 |
| Tpm2 | -4.959895038 | 5.58E-07 |
| AW112010 | 2.059761565 | 5.64E-07 |
| Pde7b | 2.228965483 | 5.66E-07 |
| Zscan18 | -2.129131312 | 5.85E-07 |
| Dmkn | 2.41622334 | 5.94E-07 |
| Zbtb10 | -2.085008416 | 6.14E-07 |
| Smpdl3a | 1.287429954 | 6.14E-07 |
| Fgfrl1 | -2.411784543 | 6.55E-07 |
| Tmem44 | -3.717463695 | 6.55E-07 |
| Lrp1 | 1.075032518 | 6.73E-07 |
| Ffar2 | 1.179959511 | 6.88E-07 |
| Rtl6 | -1.111718013 | 6.98E-07 |
| Emp1 | -7.092712731 | 7.12E-07 |
| Oas3 | 3.319150486 | 7.40E-07 |
| Prkd2 | -1.254466651 | 7.43E-07 |
| Tnfsf8 | 1.582679283 | 7.51E-07 |
| Tgm1 | 2.398801667 | 7.53E-07 |
| Gem | -6.371104752 | 7.90E-07 |
| Ccser1 | -8.149881065 | 8.34E-07 |
| Chchd10 | 1.123869699 | 8.49E-07 |
| Efna4 | -2.97847021 | 8.67E-07 |
| Sh3bp5 | 1.051271635 | 8.71E-07 |
| Parp9 | 1.022409923 | 9.17E-07 |
| Adgra3 | -1.651593272 | 9.17E-07 |
| Ssc4d | 1.195923905 | 9.17E-07 |
| Grap2 | -2.269403194 | 9.26E-07 |
| Dpf3 | -8.209652303 | 9.40E-07 |
| Pi16 | 1.243983578 | 9.94E-07 |
| Sema6c | -5.483200973 | 9.99E-07 |
| Rgl1 | -3.731000431 | 1.03E-06 |
| Slfn4 | 2.953141051 | 1.04E-06 |

| | | |
|---|---|---|
| Tgif2 | -1.262913521 | 1.04E-06 |
| Pgr | -10.9877498 | 1.05E-06 |
| Apoe | 1.538602046 | 1.05E-06 |
| Hmga2 | -10.69807302 | 1.08E-06 |
| Ptk2 | -1.961947814 | 1.10E-06 |
| Cecr2 | -5.160681984 | 1.14E-06 |
| Bex1 | -2.751077574 | 1.15E-06 |
| Adgrl2 | -2.144970832 | 1.17E-06 |
| Bend4 | -3.34484447 | 1.17E-06 |
| Arvcf | -2.306503316 | 1.21E-06 |
| Lrrc66 | -7.090043528 | 1.24E-06 |
| Colq | -7.734790122 | 1.28E-06 |
| Tmem40 | 1.309828974 | 1.28E-06 |
| Clic5 | -3.211067242 | 1.29E-06 |
| Igf2bp3 | 2.077439066 | 1.29E-06 |
| Mfsd7a | 1.905585495 | 1.38E-06 |
| Ankrd33b | -7.827569845 | 1.38E-06 |
| Soga1 | -1.402110243 | 1.40E-06 |
| Mcf2 | -7.256162598 | 1.41E-06 |
| Rnf122 | -2.51879738 | 1.41E-06 |
| Zfp395 | -1.324044168 | 1.42E-06 |
| Prx | -2.400409406 | 1.43E-06 |
| Kank2 | -1.248048084 | 1.47E-06 |
| 2810408A1 | -1.880020704 | 1.50E-06 |
| Clec2i | -1.343264738 | 1.51E-06 |
| Ptpdc1 | -2.535434993 | 1.51E-06 |
| Abcb4 | -2.964196211 | 1.51E-06 |
| Slc22a4 | 1.463544491 | 1.52E-06 |
| Cysltr1 | 1.862578108 | 1.59E-06 |
| Sox4 | -2.407208657 | 1.61E-06 |
| Itga3 | -7.112970915 | 1.69E-06 |
| Frmd6 | -1.796882024 | 1.72E-06 |
| 2900079G2 | -8.191217385 | 1.74E-06 |
| Bag3 | 1.065382903 | 1.92E-06 |
| Rcor2 | -2.881299279 | 1.95E-06 |
| Strip2 | -1.92441379 | 1.95E-06 |
| Adgrg7 | 1.686644786 | 2.10E-06 |
| Pim1 | 1.215470168 | 2.10E-06 |
| Them6 | 1.012219612 | 2.18E-06 |
| Tlr4 | 2.836207523 | 2.21E-06 |
| Pglyrp2 | -4.368271452 | 2.21E-06 |
| Tspyl4 | -1.884615059 | 2.29E-06 |
| Lrrc1 | -1.682959767 | 2.29E-06 |
| Slc12a5 | -4.17956169 | 2.33E-06 |
| Sez6l | -8.027620968 | 2.45E-06 |
| Acpp | 3.493407775 | 2.49E-06 |
| Itga8 | -3.296699708 | 2.54E-06 |

| | | |
|---|---|---|
| Gpr25 | -7.801422571 | 2.56E-06 |
| Mycl | -2.968185457 | 2.62E-06 |
| Tsc22d3 | 1.234351698 | 2.65E-06 |
| Tle6 | -1.270604268 | 2.76E-06 |
| Stat4 | -1.334870136 | 2.91E-06 |
| Nod2 | 2.469843343 | 2.95E-06 |
| Xrcc5 | -4.892517354 | 3.27E-06 |
| St8sia1 | -2.953511702 | 3.27E-06 |
| Myo18b | 2.717360114 | 3.57E-06 |
| Zfp57 | -8.010152576 | 3.59E-06 |
| Mmp11 | -2.862985534 | 3.62E-06 |
| Man2a1 | 1.045619609 | 3.69E-06 |
| Cpa3 | -7.016915247 | 3.77E-06 |
| Krba1 | -1.547030377 | 3.80E-06 |
| Tfr2 | -2.90602489 | 3.90E-06 |
| Crem | -1.714307634 | 3.92E-06 |
| Tlr7 | -1.204578855 | 4.02E-06 |
| Cand2 | -1.440542537 | 4.05E-06 |
| Slain1 | -1.614365428 | 4.06E-06 |
| Tmem231 | -2.394894342 | 4.21E-06 |
| Cbx6 | -1.031786173 | 4.34E-06 |
| Mylip | -1.000991072 | 4.48E-06 |
| Nav1 | -1.002652683 | 4.58E-06 |
| Srl | -1.292294636 | 4.91E-06 |
| Clec4a4 | 4.853105147 | 5.05E-06 |
| Rhobtb1 | -1.647488105 | 5.07E-06 |
| Oaf | -1.299875159 | 5.09E-06 |
| Ypel1 | -1.696501284 | 5.44E-06 |
| Src | -2.745918573 | 5.44E-06 |
| Rtn1 | -2.30757693 | 5.47E-06 |
| Chrnb1 | -5.129958087 | 5.49E-06 |
| Zfp979 | 2.316999345 | 5.55E-06 |
| Erg | -1.318259027 | 5.56E-06 |
| Rgcc | 2.208817866 | 5.77E-06 |
| Cachd1 | -6.143759988 | 6.04E-06 |
| Nectin3 | -6.936430091 | 6.14E-06 |
| Dgat2 | 2.484892841 | 6.35E-06 |
| Gcsam | -8.000528713 | 6.43E-06 |
| Aldh1a1 | -5.864862921 | 6.44E-06 |
| Spaca9 | -1.615357863 | 6.72E-06 |
| Cchcr1 | -1.424757816 | 6.78E-06 |
| Casp12 | 1.95327114 | 6.86E-06 |
| Fggy | 1.121070056 | 6.97E-06 |
| Kazn | 1.582869433 | 6.97E-06 |
| B430010I2 | 1.542722777 | 6.97E-06 |
| Peg12 | -3.002437821 | 7.00E-06 |
| Gpc2 | -1.473366013 | 7.12E-06 |

| | | |
|---|---|---|
| B4galnt4 | 1.333309763 | 7.12E-06 |
| Phlda3 | -1.908174735 | 7.25E-06 |
| Izumo4 | -1.582009658 | 7.37E-06 |
| Tigar | 1.280265788 | 7.56E-06 |
| Il18r1 | -4.941031778 | 7.62E-06 |
| Gpr65 | 1.14044213 | 7.69E-06 |
| Tjp1 | -6.858534583 | 7.96E-06 |
| Trim30c | 5.067872092 | 8.02E-06 |
| Slc40a1 | -5.137857994 | 8.48E-06 |
| Pmepa1 | -2.425198271 | 8.61E-06 |
| Fam214a | -2.134460808 | 8.74E-06 |
| Sbk1 | -1.350770799 | 9.01E-06 |
| Havcr1 | -6.879126809 | 9.03E-06 |
| Samd14 | -5.72578706 | 9.03E-06 |
| Rab4a | -4.135227313 | 9.23E-06 |
| Rspo1 | 3.078545915 | 9.46E-06 |
| Rarg | -1.505882346 | 9.57E-06 |
| Sh3d19 | 1.353643262 | 9.67E-06 |
| Clec4a3 | 2.546267057 | 9.82E-06 |
| Guca1a | 1.346793344 | 9.83E-06 |
| Arhgap32 | -1.204419064 | 9.89E-06 |
| Anks6 | -3.353756357 | 9.89E-06 |
| Usp44 | 2.041598529 | 9.97E-06 |
| Hnf4a | -9.058395181 | 1.07E-05 |
| Ar | -7.107611254 | 1.07E-05 |
| Sall2 | -2.200006133 | 1.07E-05 |
| Tspan9 | -6.897682906 | 1.10E-05 |
| Arhgef37 | 2.899571469 | 1.10E-05 |
| Epb41l4aos | 1.155620499 | 1.12E-05 |
| F7 | 2.359627513 | 1.14E-05 |
| Tespa1 | -1.132622978 | 1.24E-05 |
| Zcchc24 | -1.013868896 | 1.24E-05 |
| Dcaf12l1 | -7.663041364 | 1.37E-05 |
| C130050O1 | 1.150696064 | 1.39E-05 |
| Klhl4 | -7.506269678 | 1.41E-05 |
| Cadm3 | -2.836286581 | 1.43E-05 |
| Slc2a10 | -7.76820871 | 1.53E-05 |
| Kdm5b | -2.448998293 | 1.56E-05 |
| Syde2 | -2.112659253 | 1.64E-05 |
| Carhsp1 | -1.598796787 | 1.65E-05 |
| Boll | -6.314398761 | 1.71E-05 |
| Car2 | -3.933722682 | 1.71E-05 |
| Mxra7 | 4.260989308 | 1.73E-05 |
| Pcbp4 | -1.22827909 | 1.74E-05 |
| Klra2 | 7.254314872 | 1.74E-05 |
| Cfb | 5.777379577 | 1.76E-05 |
| Tspan33 | -1.857324927 | 1.77E-05 |

| | | |
|---|---|---|
| Ttc39c | 1.151711903 | 1.83E-05 |
| Armcx6 | -1.134044765 | 1.86E-05 |
| Tmem121b | -4.795816385 | 1.98E-05 |
| Pros1 | 1.11015467 | 2.05E-05 |
| Sema7a | -3.957525746 | 2.11E-05 |
| Prss34 | -8.870853529 | 2.14E-05 |
| Mecom | -7.923398863 | 2.26E-05 |
| Gfra1 | -11.22100363 | 2.31E-05 |
| Kcnq3 | -2.390730799 | 2.34E-05 |
| Fam174b | -1.995139083 | 2.38E-05 |
| Socs2 | -5.919327377 | 2.42E-05 |
| Mmp19 | 1.101579923 | 2.43E-05 |
| Cep97 | -1.04583078 | 2.54E-05 |
| Fcgr4 | 3.10342613 | 2.63E-05 |
| Ccl6 | 1.125554186 | 2.63E-05 |
| Ggt5 | -1.674311048 | 2.69E-05 |
| Armcx4 | -1.599514934 | 2.72E-05 |
| Mfsd6l | 2.814333681 | 2.73E-05 |
| Stk36 | -5.878228305 | 2.77E-05 |
| H2-T24 | 1.296513052 | 2.98E-05 |
| Hectd2 | -2.173279046 | 3.00E-05 |
| Tanc2 | -1.983948025 | 3.05E-05 |
| Sh2d6 | 2.322511018 | 3.06E-05 |
| Large2 | -4.380570197 | 3.08E-05 |
| Slc25a53 | -1.141699757 | 3.10E-05 |
| Pde4b | -1.510539964 | 3.10E-05 |
| Aifm2 | 1.5557647 | 3.11E-05 |
| Klrg2 | 7.062710451 | 3.35E-05 |
| Plscr1 | 1.338634234 | 3.62E-05 |
| Dennd2c | -1.152758475 | 3.70E-05 |
| Tmem154 | 1.199183716 | 3.72E-05 |
| Gja1 | -3.04858725 | 3.75E-05 |
| Osbpl5 | -1.351824994 | 3.91E-05 |
| Sema3d | -4.868951987 | 3.93E-05 |
| Zik1 | -1.800495429 | 4.15E-05 |
| Dach1 | -3.613924897 | 4.17E-05 |
| Gm5086 | -3.157827223 | 4.18E-05 |
| Letm2 | -1.85412021 | 4.27E-05 |
| Prss16 | 1.339889574 | 4.36E-05 |
| Fgd1 | -1.302319778 | 4.44E-05 |
| Lmna | -3.226102238 | 4.47E-05 |
| Sh3rf1 | -1.171281547 | 4.65E-05 |
| Emcn | -8.016506809 | 4.89E-05 |
| Slc22a17 | -2.32731335 | 4.99E-05 |
| Crim1 | -2.459288514 | 4.99E-05 |
| Timp2 | -2.610191528 | 4.99E-05 |
| Rnd3 | -1.169001548 | 5.29E-05 |

| | | |
|---|---|---|
| Slc9a2 | -7.146020971 | 5.34E-05 |
| Iqcb1 | -1.021522827 | 5.35E-05 |
| Fam171a1 | -3.035597106 | 5.41E-05 |
| Ephb6 | -3.209597846 | 5.50E-05 |
| Zfp808 | -1.242099754 | 5.64E-05 |
| Smim5 | -1.64961077 | 5.73E-05 |
| Gpr174 | -4.682208205 | 6.37E-05 |
| Rcn3 | -1.131150707 | 6.38E-05 |
| Cnr2 | 1.27911636 | 6.47E-05 |
| Apoc2 | 3.010878065 | 6.47E-05 |
| Rnf150 | -1.399116184 | 6.55E-05 |
| Nupr1 | 1.781751324 | 6.56E-05 |
| Hacd4 | 1.038784943 | 6.64E-05 |
| Zfp773 | -1.732387953 | 6.66E-05 |
| Sv2a | -1.879576213 | 6.75E-05 |
| Gabbr1 | -5.579515284 | 7.11E-05 |
| Arid3b | -1.739063681 | 7.12E-05 |
| Neurl1b | -1.00439694 | 7.20E-05 |
| Cd300c | -1.769996762 | 7.72E-05 |
| Slc26a11 | -1.984561108 | 7.94E-05 |
| B3gnt7 | -1.958833404 | 7.94E-05 |
| Paqr8 | -1.596817244 | 8.03E-05 |
| Tal1 | -1.455585398 | 8.07E-05 |
| Sorcs2 | -4.671792909 | 8.11E-05 |
| Sh2d2a | -7.678630061 | 8.48E-05 |
| Pdzd7 | -7.554163412 | 8.48E-05 |
| Vmn2r29 | -7.961519134 | 8.52E-05 |
| Abca4 | -6.519152979 | 8.66E-05 |
| Nol4l | -2.314256627 | 9.00E-05 |
| Eya1 | -2.276499303 | 9.09E-05 |
| Glrp1 | 2.662474818 | 9.12E-05 |
| Galm | 2.138272148 | 9.33E-05 |
| Satb2 | 1.242116067 | 9.45E-05 |
| Rhag | -4.316836618 | 9.47E-05 |
| Sv2b | -6.843831109 | 9.54E-05 |
| Slc9a9 | -1.067834977 | 0.000102 |
| Zfpm1 | -2.064151127 | 0.000104 |
| Snx29 | -1.10910958 | 0.000104 |
| St8sia2 | -7.462004903 | 0.000104 |
| Zfyve9 | 1.747455235 | 0.000105 |
| Kcnk13 | 1.99326335 | 0.000106 |
| Il27 | 3.399558949 | 0.000107 |
| Zbtb46 | -2.055050954 | 0.000108 |
| Slamf8 | 1.638495804 | 0.00011 |
| Mcub | 1.251325445 | 0.00011 |
| Hoxa10 | -4.401304346 | 0.000116 |
| Ccr1 | 1.675877502 | 0.000116 |

| | | |
|---|---|---|
| Sesn3 | -1.220876961 | 0.000116 |
| Cd276 | -7.19747063 | 0.000119 |
| Socs5 | -1.117397153 | 0.000121 |
| Slamf6 | -1.465372947 | 0.000127 |
| Ift81 | -1.580252855 | 0.000128 |
| Tmem98 | -7.18097197 | 0.00013 |
| Kcnj2 | 1.084215453 | 0.00013 |
| Mrgpre | -2.175181945 | 0.00013 |
| Ttc39a | 3.346217094 | 0.000131 |
| Ebf1 | -10.02789212 | 0.000136 |
| Tmtc1 | -6.342628394 | 0.000136 |
| Vmn2r29 | -3.825783176 | 0.000137 |
| Mmp8 | 1.409110215 | 0.000138 |
| Traf4 | -1.9464046 | 0.000139 |
| Rhbdl3 | -2.583074763 | 0.000141 |
| Slc13a3 | 3.379371375 | 0.000142 |
| Cnrip1 | -2.731071085 | 0.000144 |
| Osgin1 | 2.583247629 | 0.000145 |
| Tacc2 | -1.291244833 | 0.000161 |
| Tent5d | 7.544133764 | 0.000162 |
| Sox6 | -6.196876112 | 0.000164 |
| Galc | 1.021484995 | 0.000166 |
| Sec31b | -1.329061798 | 0.000166 |
| Hnmt | 1.456348064 | 0.000173 |
| Rem1 | 2.258642326 | 0.000174 |
| Spink2 | 4.984754882 | 0.000177 |
| C1rb | 2.28998032 | 0.000181 |
| Kctd12b | -1.534302029 | 0.000182 |
| 4930538E2 | -2.515053105 | 0.000183 |
| Hcst | 1.121511347 | 0.000184 |
| Gypa | -4.569572584 | 0.000184 |
| Smad7 | -1.091927032 | 0.000187 |
| Dusp6 | -1.681695778 | 0.000191 |
| Sgms2 | 2.566481003 | 0.000193 |
| Slc16a1 | 1.022960896 | 0.000197 |
| Slc4a3 | -4.32211409 | 0.000199 |
| Sema4g | -3.683998267 | 0.000202 |
| Fam83a | 4.814683912 | 0.000203 |
| Dennd5b | -2.479799923 | 0.000205 |
| Efcc1 | -4.031461258 | 0.000207 |
| Grik5 | -1.120763357 | 0.000207 |
| Smim3 | -1.131529093 | 0.000213 |
| Lzts2 | -1.124855984 | 0.000215 |
| Dapk1 | -1.284486059 | 0.000217 |
| Cdh24 | -1.666825724 | 0.000222 |
| Hal | 2.065561552 | 0.000225 |
| Adam3 | 3.692325482 | 0.000229 |

| | | |
|---|---|---|
| A730081D( | -2.459196004 | 0.000234 |
| Rorc | -7.903861757 | 0.000241 |
| Art2b | 7.880411656 | 0.000243 |
| Cbarp | -1.362612673 | 0.000248 |
| Mpl | -7.391286806 | 0.000254 |
| Tlr1 | 1.126317272 | 0.000255 |
| Flt1 | 2.078010845 | 0.000258 |
| Bank1 | -7.081842617 | 0.000264 |
| Armcx2 | -1.194030532 | 0.000265 |
| Ccdc150 | 2.272365676 | 0.00027 |
| Dmtn | -5.508584131 | 0.000279 |
| Ypel3 | -1.105148515 | 0.000279 |
| Rec8 | -5.262015034 | 0.000281 |
| Maml2 | -1.607299713 | 0.000289 |
| Gm30948 | -5.540808331 | 0.000303 |
| Il17re | -10.39084037 | 0.000306 |
| Reck | -2.860319416 | 0.00031 |
| Carns1 | -1.59259052 | 0.000325 |
| Ninl | -1.655681047 | 0.000339 |
| Cdc42bpa | -1.612268609 | 0.000339 |
| Bcam | -6.829160195 | 0.000343 |
| Myh10 | -2.039189504 | 0.000346 |
| 9430069I0' | 6.507114125 | 0.000347 |
| Filip1 | 2.656100146 | 0.000353 |
| Enpp6 | -7.66298289 | 0.000355 |
| Gm44805 | 2.970272491 | 0.00036 |
| Mcoln3 | -7.508507059 | 0.000374 |
| Ttc28 | -2.06914651 | 0.000377 |
| Catsperg1 | 1.795992814 | 0.000381 |
| Cacna2d4 | -4.128181452 | 0.000383 |
| F830208F2 | 1.765855881 | 0.000383 |
| Pip5k1b | -1.77439956 | 0.000386 |
| Naip1 | 1.363402947 | 0.000392 |
| Cd177 | 1.897011725 | 0.000393 |
| Zfp934 | -1.73184713 | 0.000407 |
| D630045J1 | -1.487965157 | 0.000409 |
| Fzd6 | -3.239035838 | 0.000411 |
| Cacna1b | 1.364410683 | 0.000437 |
| Fn3k | -7.506871913 | 0.00044 |
| Ddx4 | -4.250613091 | 0.00044 |
| Cpne5 | -3.470568072 | 0.000446 |
| Lrg1 | 7.113682425 | 0.000455 |
| Ak8 | -1.784452863 | 0.000456 |
| Castor2 | -1.334235243 | 0.000462 |
| Cux2 | 3.9802585 | 0.000462 |
| Epb41l5 | -3.118468088 | 0.000464 |
| Ttc21a | 1.766903616 | 0.000467 |

| | | |
|---|---|---|
| Wdr78 | -1.904251747 | 0.000477 |
| Adamtsl2 | -5.744660831 | 0.000478 |
| Clec4b2 | 6.857609381 | 0.000478 |
| Cfh | -1.743192897 | 0.000483 |
| Zfp579 | -1.347865177 | 0.000488 |
| Meis3 | -3.409289639 | 0.000488 |
| Myo1b | 1.916994041 | 0.000495 |
| Agmo | 7.16621449 | 0.000499 |
| Begain | -6.84398474 | 0.000507 |
| Pmaip1 | 1.263103912 | 0.000514 |
| Rgs1 | -4.668237485 | 0.000523 |
| Hsd11b1 | 1.860252158 | 0.00053 |
| Gpr15 | 2.382679875 | 0.000533 |
| Cd79a | -5.508009119 | 0.000534 |
| Klf1 | -3.656811691 | 0.000546 |
| Gimap4 | -3.976703424 | 0.000549 |
| 2010016I1 | 1.59596506 | 0.000551 |
| Cldn34c1 | -2.36396449 | 0.000568 |
| Epb42 | -6.322220151 | 0.000577 |
| Rasl12 | -7.056678263 | 0.000577 |
| Gpr45 | -4.747081879 | 0.000577 |
| Pard3b | -1.326285341 | 0.000577 |
| Dennd2a | -2.532692466 | 0.000596 |
| Nkd1 | -7.300374564 | 0.000616 |
| Ncs1 | -1.824175296 | 0.000624 |
| Myog | -7.052010175 | 0.000628 |
| H2-Eb2 | -6.857636598 | 0.00063 |
| Ifi206 | 2.523352331 | 0.000651 |
| Trim36 | -1.16600884 | 0.000652 |
| Actn2 | -7.075327567 | 0.000662 |
| 5830444B0 | -2.951563375 | 0.000671 |
| Fv1 | -1.293488197 | 0.000684 |
| Smpdl3b | 3.191163105 | 0.000708 |
| Clec4b1 | 2.021369095 | 0.000712 |
| Gata3 | -7.15278346 | 0.000714 |
| Slc1a4 | 2.122404611 | 0.000717 |
| Plekha6 | -1.30341595 | 0.000719 |
| Lgr4 | 1.160218318 | 0.000737 |
| Pcyt1b | -3.776260401 | 0.000741 |
| Porcn | 3.905006909 | 0.000756 |
| Mturn | -3.074753353 | 0.00077 |
| Mzb1 | -5.579587907 | 0.00078 |
| Ptgs1 | 1.62295993 | 0.000782 |
| Ccdc120 | -3.02483683 | 0.000803 |
| Gfra4 | -6.235857417 | 0.000819 |
| Smpd3 | -1.845163365 | 0.000827 |
| Gla | 1.275862626 | 0.00083 |

| | | |
|---|---|---|
| Dkkl1 | -3.783433079 | 0.000836 |
| Acot3 | 1.593558386 | 0.000838 |
| Apol7d | 2.067258144 | 0.000842 |
| Bvht | 3.948333908 | 0.000842 |
| D930048N: | -2.196607917 | 0.000844 |
| Bst1 | 2.25012244 | 0.000859 |
| Sbspon | -4.61038679 | 0.000869 |
| Cd33 | -1.01222703 | 0.000877 |
| Sytl4 | -3.632321472 | 0.000887 |
| Camsap3 | -3.200121336 | 0.000897 |
| Zfp873 | -1.180701516 | 0.000955 |
| Ccl24 | 5.931004171 | 0.000955 |
| Sdc3 | 1.114934548 | 0.000978 |
| Hes1 | -5.332767976 | 0.000986 |
| Azin2 | -1.256599203 | 0.001005 |
| Pnck | -2.881022418 | 0.001005 |
| Atp9a | 1.226356697 | 0.001006 |
| Tert | -2.779152858 | 0.001011 |
| Gm16548 | 2.488372654 | 0.001013 |
| Gfi1b | -4.517956938 | 0.001016 |
| Mmp25 | 3.832760965 | 0.00104 |
| Atat1 | -1.25612257 | 0.001064 |
| Sh3tc2 | -7.507583897 | 0.001064 |
| Gpc3 | 1.721219013 | 0.001077 |
| Abcg4 | -4.132791649 | 0.001078 |
| Plpp1 | -3.840788736 | 0.00108 |
| Nectin2 | 1.30169549 | 0.001086 |
| Lrp4 | 1.890492169 | 0.001094 |
| Methig1 | 1.474746734 | 0.001125 |
| Ifit1 | 2.440562962 | 0.001134 |
| Tlr13 | 2.185782921 | 0.001216 |
| AI854703 | -2.431343095 | 0.001219 |
| Pira6 | 1.063079005 | 0.001224 |
| Stbd1 | -1.374204605 | 0.001231 |
| Krt7 | -3.480405149 | 0.001231 |
| P2ry12 | -3.444273317 | 0.001257 |
| Fbn1 | 1.721720033 | 0.001287 |
| Inpp5j | 5.917905155 | 0.00129 |
| Kif5a | -1.834953524 | 0.001291 |
| A930037H( | 1.218142129 | 0.001315 |
| Car1 | -3.316643392 | 0.001321 |
| Hdc | 1.21480003 | 0.001346 |
| Arl3 | -1.017910595 | 0.001352 |
| Samd4 | -7.542915984 | 0.001368 |
| Fn1 | 2.008746685 | 0.001371 |
| Glis3 | 1.16394022 | 0.001385 |
| Dyrk4 | -4.829748035 | 0.001388 |

| | | |
|---|---|---|
| Zfp507 | -1.203550547 | 0.001388 |
| Gm16731 | -1.009435709 | 0.001432 |
| Zfp658 | -1.486670587 | 0.001446 |
| Sh2d4a | -6.294478647 | 0.00145 |
| Kcnab3 | -1.867755372 | 0.001458 |
| Cc2d2b | -6.129501105 | 0.001466 |
| Sult1a1 | 2.410327732 | 0.001493 |
| Hs3st1 | -5.372461547 | 0.001496 |
| Ltb | -1.501327814 | 0.001497 |
| Ppp1r36 | -3.428556201 | 0.001516 |
| Igsf9 | -3.389943979 | 0.001522 |
| Per1 | 1.615887039 | 0.001523 |
| Crispld1 | -6.995812462 | 0.001531 |
| Ackr1 | -5.523444393 | 0.001534 |
| Fam110b | -2.17593763 | 0.001591 |
| Dyrk1b | -1.149290701 | 0.001597 |
| Armcx1 | -2.025272513 | 0.001616 |
| Bex4 | -2.693852253 | 0.001659 |
| Endou | -3.30754107 | 0.00174 |
| Cbr1 | -1.013112427 | 0.001746 |
| Prkcq | -4.792728095 | 0.001754 |
| Nlrp1c-ps | 1.872583793 | 0.001756 |
| Gm13031 | -2.107039594 | 0.001848 |
| Vangl1 | -5.112501835 | 0.001854 |
| Clcn2 | -1.421656096 | 0.001857 |
| Pir | 1.466816617 | 0.001866 |
| Mocos | 1.413501577 | 0.001901 |
| Trpc2 | -1.354651635 | 0.001901 |
| Cish | -1.667819303 | 0.001958 |
| Gdpd1 | -4.511393638 | 0.002015 |
| Nsg2 | -5.038247844 | 0.002043 |
| Art4 | -5.581404562 | 0.002064 |
| Pdzk1ip1 | -5.975349893 | 0.002084 |
| Six5 | -3.489994639 | 0.002118 |
| Ppp2r3a | -1.011756591 | 0.002223 |
| Psd3 | -1.061909223 | 0.002224 |
| Pkhd1l1 | -6.717071902 | 0.002224 |
| Epha7 | -1.85879525 | 0.002255 |
| Fgfr1 | -2.247290194 | 0.002277 |
| Adamts6 | -1.187219374 | 0.002286 |
| Arhgap42 | -5.597570948 | 0.002346 |
| Pard6g | -1.507065155 | 0.002346 |
| Tnfsf11 | -5.375508053 | 0.002391 |
| Trp53inp1 | -1.354422909 | 0.002438 |
| Cxcl12 | -8.129270943 | 0.002438 |
| Plekhg4 | 2.90926196 | 0.002443 |
| Serpinh1 | -1.711387384 | 0.002451 |

| | | |
|---|---|---|
| Spred3 | -1.217733257 | 0.002467 |
| Kel | -5.350027163 | 0.002514 |
| Rapsn | 1.537251042 | 0.002534 |
| Tshz3 | -5.660852719 | 0.002541 |
| Plk2 | -2.835880496 | 0.002632 |
| Raph1 | -1.022572769 | 0.002721 |
| Tox3 | -6.863071602 | 0.002782 |
| BC025920 | -1.809958865 | 0.00279 |
| Casc4 | -1.489412709 | 0.002807 |
| Thy1 | -3.064559271 | 0.002807 |
| 1700001C1 | -3.202217458 | 0.00282 |
| Gm15408 | -6.278190018 | 0.002897 |
| Gm3383 | -2.090446619 | 0.002906 |
| Actn3 | -6.502285828 | 0.002922 |
| Nrg4 | 1.024987835 | 0.002936 |
| Acy3 | 1.716612773 | 0.003184 |
| Enc1 | -1.114504769 | 0.003212 |
| Cd209d | -4.088993979 | 0.003223 |
| Il2rb | -3.890534666 | 0.00327 |
| Pax5 | -8.649853901 | 0.003304 |
| Fbxo32 | -1.791915498 | 0.003308 |
| St3gal6 | -3.015775616 | 0.003404 |
| Igdcc4 | -3.733318837 | 0.003404 |
| Mrc2 | -7.345064771 | 0.003517 |
| Hmgcs2 | -5.286683917 | 0.00353 |
| Mir99ahg | -6.249687324 | 0.003547 |
| Pcdh7 | 5.174950702 | 0.003548 |
| Ank1 | -3.778834984 | 0.003556 |
| C230035I1 | 2.114406731 | 0.003561 |
| Tek | -5.981580219 | 0.003628 |
| Dscam | -4.128075601 | 0.003663 |
| Mboat2 | -3.182914013 | 0.003713 |
| Pdlim4 | -2.393146726 | 0.003739 |
| Wif1 | -6.823107614 | 0.003769 |
| Socs1 | 1.936435364 | 0.003857 |
| Optn | -2.239964716 | 0.003922 |
| Prss30 | -1.976179661 | 0.003923 |
| Scin | -2.840205586 | 0.00395 |
| Plod2 | -2.336175617 | 0.003966 |
| Sfmbt2 | -1.598375626 | 0.003967 |
| Itga2b | -3.193554058 | 0.004173 |
| Mgll | -5.000661474 | 0.004188 |
| Adam19 | -1.083810606 | 0.004323 |
| Col5a1 | -4.231194082 | 0.004328 |
| Ermap | -4.135973836 | 0.00433 |
| Tox2 | -5.509122251 | 0.004356 |
| Slc28a2b | 1.141878759 | 0.004392 |

| | | |
|---|---|---|
| Phf11c | 1.269996117 | 0.004398 |
| Kcnh7 | 2.351524455 | 0.004418 |
| Shroom1 | -3.813875525 | 0.00457 |
| Txlnb | -1.953893936 | 0.004642 |
| Pfn2 | -4.999797448 | 0.004809 |
| Inpp4b | -2.875190765 | 0.00482 |
| Ccl2 | 2.059812244 | 0.004904 |
| Mpped2 | -6.764673364 | 0.004921 |
| Smim10l2a | -2.204936601 | 0.004963 |
| Mmel1 | -3.851993497 | 0.004967 |
| St8sia6 | -5.331424162 | 0.005027 |
| Mterf2 | -1.495592654 | 0.00504 |
| Dagla | 5.278793983 | 0.005081 |
| Mmp2 | -7.007671789 | 0.005321 |
| Fcrl1 | -1.005818871 | 0.005327 |
| Myo1d | -4.942243915 | 0.005367 |
| Stra6l | 2.334377533 | 0.005616 |
| Tmem8b | -1.345816076 | 0.005627 |
| Ccdc116 | 1.399508432 | 0.005796 |
| Eps8l2 | -4.933515355 | 0.005806 |
| Zfp113 | -1.138270926 | 0.005879 |
| LOC11548[illegible] | 4.430876463 | 0.005895 |
| Hmox1 | 1.752357204 | 0.005952 |
| Fcnb | 1.787859681 | 0.005974 |
| Ctla4 | -5.999891668 | 0.006012 |
| Igfbp7 | -6.404048978 | 0.006041 |
| Cdcp1 | -6.665234037 | 0.006043 |
| Hoxa2 | -3.279699857 | 0.006103 |
| Gnb4 | -1.241392459 | 0.006145 |
| Sox5 | -6.699737021 | 0.006175 |
| Tspan8 | -5.322990829 | 0.006237 |
| Mettl7a2 | 1.236738948 | 0.006243 |
| Rfx8 | -5.613834635 | 0.006309 |
| Rgs7bp | -5.005063717 | 0.006314 |
| Lgalsl | 1.276001018 | 0.006416 |
| Uaca | -1.910069382 | 0.006429 |
| Klf9 | 1.317864271 | 0.006438 |
| Nhsl2 | 1.206151797 | 0.006464 |
| Gm15972 | 2.708082609 | 0.006546 |
| Gbp9 | 1.334381586 | 0.006551 |
| Igf2r | -3.326371102 | 0.006554 |
| Raet1e | 2.312809329 | 0.006674 |
| Lmntd2 | -1.629701525 | 0.006741 |
| Synpo | -3.644832882 | 0.006772 |
| Gm4924 | -1.646272619 | 0.00685 |
| Peg10 | -6.639626138 | 0.006854 |
| P2ry10 | -1.940795035 | 0.006929 |

| | | |
|---|---|---|
| Fbln1 | -2.625830091 | 0.006961 |
| 4930555A( | -1.308862089 | 0.006987 |
| Patj | -2.628250853 | 0.007044 |
| Pla2r1 | 3.53204327 | 0.00706 |
| Cep126 | -3.235324683 | 0.007174 |
| Mak | 6.359953039 | 0.007435 |
| Synpo2l | -3.536139721 | 0.00747 |
| Adam22 | -1.968052268 | 0.007481 |
| Fhl1 | -5.949437492 | 0.007493 |
| Mei4 | -2.546341863 | 0.007493 |
| Gm42372 | -3.605662078 | 0.007534 |
| Tlr6 | 1.092678217 | 0.007553 |
| Gm5914 | -1.345416602 | 0.007683 |
| Hotairm1 | 2.109256013 | 0.007686 |
| Slit1 | -3.740383981 | 0.007756 |
| Lin28a | 3.810976994 | 0.007859 |
| Rnase2a | 3.942257658 | 0.008199 |
| Tmem198 | -5.692577503 | 0.008607 |
| Mamld1 | -5.526943094 | 0.00867 |
| 2010300C( | -2.886746099 | 0.008681 |
| Dnajb2 | -1.217250246 | 0.008689 |
| Acsm3 | -1.110589374 | 0.008689 |
| Ankrd6 | -1.955117236 | 0.008728 |
| Slc18a1 | -2.257282451 | 0.008863 |
| Tmem121 | -8.802207749 | 0.008896 |
| Rai2 | -5.976594136 | 0.009005 |
| Myo6 | -4.391367305 | 0.009058 |
| Etl4 | -5.826873862 | 0.009069 |
| Spocd1 | 5.205190299 | 0.009273 |
| Rasgrp4 | -1.152162737 | 0.009344 |
| Gm10505 | -2.139791629 | 0.009357 |
| Arhgef15 | 2.083562991 | 0.009363 |
| E130215H2 | -4.67812912 | 0.009366 |
| Cdkl3 | 1.212325657 | 0.00938 |
| Slc27a2 | -3.361646389 | 0.009453 |
| Tmem221 | -1.567311899 | 0.009512 |
| Gpr155 | -1.008544217 | 0.009595 |
| Slc11a1 | 1.443804076 | 0.009835 |
| Spef1 | -1.231550637 | 0.009857 |
| B3galt5 | -2.331589302 | 0.009913 |
| Itga9 | -8.645426769 | 0.009972 |
| Tnfaip8l3 | 3.953189441 | 0.010041 |
| Cxcl9 | 3.656510868 | 0.010236 |
| Traf1 | -2.75455038 | 0.010256 |
| Dnah7a | -7.303104322 | 0.010324 |
| Trim6 | 1.450163216 | 0.010329 |
| Oas1c | 1.96055658 | 0.010409 |

| | | |
|---|---|---|
| Slc24a2 | -6.265498417 | 0.010413 |
| Egfr | -4.545191168 | 0.010441 |
| Plcl1 | -2.617201314 | 0.010524 |
| Ust | -1.657592061 | 0.010749 |
| Gjc1 | -5.522192666 | 0.010792 |
| Apol7e | -2.724993524 | 0.010827 |
| Pdgfc | -5.668336658 | 0.010883 |
| Ccdc173 | -1.909208108 | 0.010982 |
| Kcnj11 | -6.872031477 | 0.011045 |
| Sytl3 | -6.066775051 | 0.011123 |
| Zfhx2 | -1.031995139 | 0.011277 |
| Acta2 | 6.09843641 | 0.011384 |
| Lrrn4 | -3.243825165 | 0.011405 |
| Nrep | -6.523376506 | 0.011602 |
| Zfp867 | -1.467737338 | 0.011875 |
| Obsl1 | -5.916329299 | 0.012006 |
| Auts2 | -2.289794785 | 0.012031 |
| Ankmy1 | 1.283388231 | 0.012082 |
| Rhof | -1.388134532 | 0.012162 |
| Gstm5 | -1.029308597 | 0.012224 |
| Nexn | 4.744351435 | 0.012224 |
| Ly75 | 1.179111914 | 0.012235 |
| Gm16675 | 1.448645876 | 0.012323 |
| Ccdc9b | -1.699450629 | 0.012344 |
| Zfp708 | -1.32802926 | 0.012522 |
| Col16a1 | -5.882838073 | 0.012575 |
| Gm10406 | -3.625737318 | 0.012688 |
| Rab11fip3 | -1.020653851 | 0.012785 |
| Ttc41 | -1.250226033 | 0.012886 |
| Plk3 | 2.082578633 | 0.012924 |
| 1500009L1 | -4.768527482 | 0.013079 |
| Insyn2b | 4.964157906 | 0.013218 |
| Hes5 | -3.786979259 | 0.013228 |
| Gm16001 | 1.276259683 | 0.013309 |
| Notch3 | -2.998679238 | 0.013441 |
| Lrig1 | -1.845300606 | 0.013458 |
| Cxcr5 | -6.391309081 | 0.013649 |
| Gm5111 | -6.2843421 | 0.013668 |
| Fbxl2 | 1.038381918 | 0.013903 |
| Akap12 | -1.997599784 | 0.013925 |
| Tlr8 | 1.419588501 | 0.014039 |
| Arhgap6 | -3.23185022 | 0.014063 |
| Cyp2j9 | -1.609569677 | 0.014211 |
| Itgb4 | -1.329381167 | 0.014259 |
| Add2 | -3.243976155 | 0.014418 |
| Prelid2 | -1.238967903 | 0.014596 |
| Shisa3 | 4.850393823 | 0.014736 |

| | | |
|---|---|---|
| Nkx2-3 | -6.007550286 | 0.015002 |
| Slc36a2 | 1.698683408 | 0.015167 |
| Serinc2 | 5.618245574 | 0.015231 |
| Mtus2 | 1.13121999 | 0.015352 |
| Nsg1 | -1.849170528 | 0.015707 |
| Blvrb | -1.14223191 | 0.015707 |
| Cdh5 | -2.396771649 | 0.016078 |
| Scarf2 | 2.093858841 | 0.016231 |
| Gdf11 | -4.768516155 | 0.016259 |
| Unc5a | -1.214912017 | 0.016307 |
| Dipk1b | -1.326411206 | 0.016379 |
| Gm2897 | -2.536495188 | 0.016548 |
| Ociad2 | -1.41447765 | 0.016573 |
| Ltb4r1 | 1.193758438 | 0.016591 |
| Spon2 | -6.488399803 | 0.016702 |
| 4931431B1 | -4.237519196 | 0.016718 |
| Gm1976 | 1.057233339 | 0.016843 |
| Hoxb3 | 1.238861138 | 0.017048 |
| Arap3 | -1.08705415 | 0.017087 |
| BC016579 | -5.874835883 | 0.01725 |
| Nrbp2 | -3.771741963 | 0.017297 |
| Tnip3 | -2.161598001 | 0.01733 |
| Myo5c | -4.713875295 | 0.017434 |
| Fcer2a | -3.395074265 | 0.017559 |
| Slc4a5 | -3.040932574 | 0.017574 |
| Aqp9 | -1.877160072 | 0.017806 |
| Col4a2 | -1.896693382 | 0.018021 |
| Klra1 | -5.3740858 | 0.018085 |
| Alas2 | -3.741788463 | 0.018128 |
| Tpmt | -1.268851803 | 0.018531 |
| Ifit1bl2 | 4.633932254 | 0.018531 |
| Sapcd1 | -1.568354394 | 0.018585 |
| Plvap | -1.816949849 | 0.018613 |
| Slc37a3 | -1.418656903 | 0.019146 |
| Spic | -5.146952641 | 0.019531 |
| Lhfpl2 | -1.038654333 | 0.01983 |
| Fgd6 | 2.155711325 | 0.019925 |
| Rab6b | -2.193965969 | 0.02022 |
| Cfap53 | 6.150488866 | 0.020286 |
| Il7 | 3.17508957 | 0.02041 |
| Kcnk5 | -1.700002657 | 0.020444 |
| Gprc5c | -4.890118446 | 0.020459 |
| B230208H1 | 1.292903555 | 0.020469 |
| Zfp566 | -1.117293256 | 0.020527 |
| Slc22a18 | 1.081227028 | 0.020548 |
| 6820408C1 | -6.105458638 | 0.020647 |
| Mgam | 2.469643962 | 0.020808 |

| | | |
|---|---|---|
| Gas2l1 | 1.063084655 | 0.020887 |
| Plppr2 | 1.705227058 | 0.021069 |
| Sycp3 | 2.19222459 | 0.02161 |
| Mrgpra2b | 3.267526653 | 0.021803 |
| Myom3 | -6.483843978 | 0.021853 |
| Them7 | -1.801849079 | 0.021898 |
| Col23a1 | -3.059793377 | 0.021903 |
| Vwa7 | -1.599920066 | 0.021934 |
| Cd200 | -3.571868777 | 0.022 |
| Pyroxd2 | 1.2910751 | 0.022233 |
| Slc39a4 | 1.806969319 | 0.022426 |
| Nacad | -2.994330746 | 0.022493 |
| Npr2 | -8.665821263 | 0.022574 |
| H2-Q2 | 3.298952709 | 0.022746 |
| Msrb2 | -1.104295943 | 0.022987 |
| Acot4 | 3.151827597 | 0.023151 |
| Fhl2 | -4.053807389 | 0.02317 |
| Pipox | 3.108993605 | 0.02317 |
| Gm2974 | -1.729821096 | 0.02317 |
| Eaf2 | -2.841053682 | 0.023427 |
| 5830418P1 | -2.491836567 | 0.023592 |
| Cldn7 | 6.301346335 | 0.024085 |
| Serpini1 | -1.623657993 | 0.024444 |
| Gpr85 | 2.257294713 | 0.024871 |
| Plekha4 | 2.151204795 | 0.025465 |
| Tmem91 | -2.174411898 | 0.025465 |
| Adgrf3 | 5.315224114 | 0.025675 |
| Prrg4 | -1.710257947 | 0.025698 |
| Foxd2 | -1.674598176 | 0.025698 |
| Rsph9 | -3.710539303 | 0.025772 |
| 5830432E0 | -1.245345506 | 0.025788 |
| Srpk3 | -2.792787174 | 0.025869 |
| Klre1 | -6.224513115 | 0.025871 |
| Arhgef17 | -2.323665968 | 0.025927 |
| Mafb | 2.302768891 | 0.026018 |
| Nuak1 | -3.957895293 | 0.026025 |
| Slc16a11 | -1.558473617 | 0.026038 |
| Plekhg1 | 1.003538519 | 0.026242 |
| Igf2bp2 | 1.192396874 | 0.02635 |
| Ccdc63 | -1.816472377 | 0.026693 |
| Padi6 | -5.860781019 | 0.026752 |
| Hoxb5 | 6.379877241 | 0.027054 |
| Kif1a | 3.900188191 | 0.027087 |
| Cald1 | -7.195227503 | 0.027296 |
| Cd80 | 1.718869927 | 0.027438 |
| Slc24a3 | -5.8805088 | 0.027591 |
| Ppm1l | -1.906721605 | 0.027634 |

| | | |
|---|---|---|
| Lyz2 | 1.040876428 | 0.028259 |
| Bfsp1 | 4.420232031 | 0.028337 |
| Scml4 | -2.442682107 | 0.028375 |
| Plcd3 | -1.79628708 | 0.028629 |
| Pglyrp1 | 2.561242857 | 0.02865 |
| Kitl | -4.013870515 | 0.028778 |
| Arhgap22 | -1.391657634 | 0.02905 |
| Hif3a | -4.627946238 | 0.029339 |
| Ifi208 | 2.640606488 | 0.029358 |
| Kcnab1 | 3.196362529 | 0.029673 |
| Zfp879 | -6.509211805 | 0.029673 |
| Penk | 4.488430911 | 0.029728 |
| Acsl6 | -7.219807625 | 0.029755 |
| Mlxipl | 6.423463646 | 0.029768 |
| Calr3 | 1.210482422 | 0.029879 |
| Gbgt1 | 6.323501986 | 0.029927 |
| Rhoc | -4.724594892 | 0.030321 |
| Aif1l | -4.737958876 | 0.030342 |
| Zfp975 | -1.032691264 | 0.030379 |
| Gm3696 | -2.870233314 | 0.030755 |
| Ect2l | 5.783660569 | 0.031154 |
| Lgr5 | -2.114019612 | 0.03125 |
| Usp27x | -1.125241867 | 0.031488 |
| Ddit4l | -6.739356642 | 0.031544 |
| Slc6a12 | -2.084968016 | 0.032256 |
| Trim69 | 5.652544489 | 0.032298 |
| Plxna3 | -1.213360751 | 0.032309 |
| Pcolce | -2.358460475 | 0.032461 |
| Zfp30 | -3.964496706 | 0.032484 |
| Alpk3 | -3.344742328 | 0.032533 |
| Prkcz | -1.457967558 | 0.032558 |
| Gm3636 | -2.630564577 | 0.032585 |
| Rap1gap | 1.305732579 | 0.033152 |
| Myh14 | -6.903772328 | 0.033269 |
| Gm12709 | 1.573887435 | 0.034059 |
| Hoxb6 | 5.035994748 | 0.034816 |
| Chst1 | 3.776347294 | 0.034894 |
| Cd163l1 | -7.034516177 | 0.034992 |
| Dach2 | -2.964629007 | 0.034994 |
| Abtb2 | -1.952720262 | 0.03518 |
| Reps2 | -3.603594881 | 0.035243 |
| Rasgrp3 | -1.757899443 | 0.035958 |
| Aldh1l2 | 6.420339012 | 0.035964 |
| Rab20 | 2.054039755 | 0.036346 |
| Spag6l | -4.88272262 | 0.036661 |
| Tnfrsf18 | -1.08402922 | 0.037139 |
| Cfap74 | -2.242913158 | 0.037455 |

| | | |
|---|---|---|
| Ppef2 | 4.908384175 | 0.037664 |
| Dusp2 | -1.449255322 | 0.038073 |
| Tex26 | -6.13220903 | 0.038879 |
| Zc3h12b | -2.163601573 | 0.039058 |
| 4833418N( | -1.508636064 | 0.039058 |
| Ell2 | -1.371454733 | 0.039077 |
| 1600010M | 1.40571975 | 0.039222 |
| Cebpe | 1.903520028 | 0.039266 |
| Gm3667 | -4.437958075 | 0.039484 |
| Celsr3 | 2.725151102 | 0.039646 |
| Gna14 | -4.4582148 | 0.039698 |
| Slc34a1 | -1.799449142 | 0.040522 |
| Aire | -3.668570416 | 0.040702 |
| Nrxn1 | -3.208777342 | 0.040904 |
| Fgf1 | 2.410603802 | 0.041173 |
| Spag1 | -1.254845182 | 0.041326 |
| Daam2 | 1.61354401 | 0.041818 |
| 6430571L1 | -3.543742301 | 0.041983 |
| Fgfbp3 | -1.269661336 | 0.042089 |
| Slc44a4 | 3.269183274 | 0.042174 |
| Shank3 | -6.847991297 | 0.042195 |
| Upk3bl | 1.372578671 | 0.042213 |
| 2210416O1 | -3.475981106 | 0.042258 |
| Fam169b | -2.868676729 | 0.042341 |
| Tg | 4.343096361 | 0.042831 |
| Hectd2os | -1.603732405 | 0.04294 |
| Tnni1 | -5.357665231 | 0.043364 |
| B430319G1 | -4.875471197 | 0.04343 |
| Gsg1 | -3.643950255 | 0.043478 |
| Btnl9 | -1.169051343 | 0.043491 |
| Gucy1a1 | -5.160967846 | 0.043585 |
| Celf4 | -2.628927455 | 0.044349 |
| Antxr1 | 1.281589881 | 0.045247 |
| Cd8a | -1.631758384 | 0.045451 |
| 1600023N1 | 5.476437102 | 0.045463 |
| Gfod2 | -1.381255882 | 0.045565 |
| Gab3 | -1.178007637 | 0.045622 |
| Sptbn5 | 3.009563163 | 0.045867 |
| Npff | 1.367404538 | 0.046376 |
| Ntn4 | -5.310315369 | 0.04683 |
| Kctd19 | 6.681227559 | 0.046971 |
| Slc38a5 | -4.49563974 | 0.047019 |
| Celsr2 | -1.033936282 | 0.047019 |
| A930007I1 | 2.722817251 | 0.047845 |
| Fcrla | -2.209550197 | 0.04788 |
| Stxbp4 | -1.111090627 | 0.047994 |
| Shcbp1l | 1.068820452 | 0.048102 |

| | | |
|---|---|---|
| Cacna1c | -7.582137958 | 0.048559 |
| 5830411N( | -5.84910971 | 0.048972 |
| Vcan | 2.877628589 | 0.049018 |
| Tmeff1 | 3.020100629 | 0.049603 |
| Osm | 1.062047203 | 0.049697 |
| Ttyh1 | 8.226355172 | 0.049787 |
| D7Ertd443 | -6.013749347 | 0.04985 |
