## Supplemental Table 3 for "Genomic Analysis of Progenitors in Viral Infection Implicates Glucocorticoids as Suppressors of Plasmacytoid Dendritic Cell Generation"

Supplementary Table 3. List of genes that are differentially expressed in pro-DCs treated with 50nM corticosterone (CORT) or vehicle for 3.5 days by RNA-seq.
Differentially expressed genes were defined by FDR < 0.1 by DESeq2.

| Symbol | log2FoldChange (CORT vs. vehicle) | FDR |
|---|---|---|
| Ighg2b | -11.14994154 | 0.013333 |
| Unc79 | -7.399248777 | 0.000213 |
| Etl4 | -7.299057048 | 0.001056 |
| P3h2 | -6.896152573 | 0.014866 |
| Gm26533 | -6.39200726 | 0.049136 |
| Cald1 | -6.072019282 | 0.03647 |
| Mmp15 | -5.910254405 | 0.009013 |
| Vangl1 | -5.784412609 | 0.070394 |
| Reln | -5.451770573 | 0.049726 |
| Cd19 | -5.372177746 | 0.077544 |
| Nipal1 | -5.186971303 | 0.030493 |
| Cacna1h | -5.170559638 | 0.054174 |
| Lrrc66 | -4.937316437 | 0.067183 |
| Epcip | -4.921597155 | 0.089366 |
| Arhgef26 | -4.663854286 | 0.016796 |
| Gm5547 | -4.577603383 | 0.023881 |
| Hic1 | -4.571100037 | 0.000588 |
| Pax5 | -4.561778865 | 0.084584 |
| Tubb1 | -4.549413495 | 0.000829 |
| Iglc2 | -4.379781773 | 0.004651 |
| Mzb1 | -4.333929715 | 0.002999 |
| Wdr86 | -4.298196933 | 0.054755 |
| Jchain | -4.254932589 | 0.051462 |
| Igkc | -4.212426877 | 0.098264 |
| Ptgfr | -4.131792698 | 0.001395 |
| Rag1 | -4.114138636 | 0.068229 |
| Gucy2e | -4.067062905 | 0.002999 |
| Slc16a9 | -4.066201731 | 0.018043 |
| Cd247 | -4.045559149 | 0.036029 |
| Grk4 | -3.891876757 | 0.000474 |
| Gm3650 | -3.793726129 | 0.014824 |
| Sec14l2 | -3.777977479 | 0.01718 |
| Psd2 | -3.702778333 | 0.054641 |
| Gpr55 | -3.669004325 | 0.010495 |
| Tmem178b | -3.660749287 | 0.092089 |
| Gimap4 | -3.564027046 | 0.000492 |
| Syt13 | -3.502664743 | 0.069042 |
| Gimap3 | -3.500918294 | 0.012723 |
| Adamtsl1 | -3.497961558 | 0.003522 |
| Samd13 | -3.471517806 | 0.000738 |

| | | |
|---|---|---|
| Megf10 | -3.471319093 | 0.000908 |
| Skap1 | -3.46265175 | 0.000304 |
| Lmo7 | -3.442132078 | 0.059667 |
| Blk | -3.305896131 | 0.030557 |
| Synpo2 | -3.292809438 | 0.054457 |
| Trib2 | -3.275952637 | 0.048994 |
| Lct | -3.238505891 | 0.070994 |
| Cacna1e | -3.236204286 | 0.002131 |
| Pls1 | -3.09633362 | 0.06151 |
| Plg | -3.090961975 | 0.036045 |
| Grb10 | -3.072240063 | 0.012245 |
| Ifit1bl1 | -3.046251899 | 0.013752 |
| Mycn | -2.956137997 | 0.032469 |
| Irag1 | -2.934528294 | 0.000111 |
| BC016579 | -2.896672241 | 0.002111 |
| Fat3 | -2.847169018 | 0.002344 |
| Abca4 | -2.843782529 | 0.035716 |
| Ccr3 | -2.830062472 | 1.88E-11 |
| Gprasp2 | -2.820683008 | 0.076467 |
| Nckap1 | -2.816420034 | 0.000495 |
| Chst1 | -2.804479357 | 0.052967 |
| Mogat2 | -2.798752222 | 0.054641 |
| Pnma8c | -2.798440251 | 0.040455 |
| F2rl2 | -2.798183934 | 0.007604 |
| Tmem98 | -2.792861156 | 0.028425 |
| Ceacam10 | -2.788024186 | 0.066367 |
| Ltf | -2.772807344 | 0.007763 |
| Csgalnact1 | -2.770573605 | 0.002311 |
| Gm35147 | -2.750150755 | 0.003854 |
| Rnase12 | -2.745243943 | 4.94E-06 |
| Aldh1a1 | -2.730839035 | 0.094996 |
| Myct1 | -2.713158019 | 0.04993 |
| Myom1 | -2.704832684 | 0.051629 |
| Zfp941 | -2.701605456 | 0.064038 |
| Olr1 | -2.700659909 | 0.000373 |
| Slc6a4 | -2.690129122 | 1.47E-06 |
| Cttn | -2.68739306 | 0.002996 |
| Gp6 | -2.67614646 | 0.085329 |
| Mcpt8 | -2.663924506 | 2.32E-11 |
| Xrcc5 | -2.657707424 | 0.040146 |
| Ltbp1 | -2.657463359 | 0.021067 |
| Tmem198 | -2.653564806 | 0.039179 |
| AI854703 | -2.629193543 | 0.098264 |
| Apol7e | -2.609594001 | 0.003052 |

| | | |
|---|---|---|
| Klhl30 | -2.605405237 | 0.066035 |
| Gm12758 | -2.601424232 | 0.090605 |
| Tmcc3 | -2.598105443 | 2.56E-05 |
| Tjp1 | -2.595954939 | 2.44E-05 |
| Cracd | -2.594824694 | 0.069611 |
| Gp5 | -2.594794915 | 0.022853 |
| Cmah | -2.593466507 | 0.010185 |
| Il6 | -2.575919484 | 1.52E-12 |
| Alox12 | -2.562175359 | 0.083931 |
| Ldhc | -2.553868157 | 0.034165 |
| Cavin2 | -2.55379768 | 0.025839 |
| Lax1 | -2.551462926 | 0.00209 |
| St6galnac5 | -2.537667012 | 7.60E-06 |
| Htr1b | -2.490410394 | 0.000458 |
| Gm10505 | -2.485470943 | 0.066316 |
| Robo3 | -2.474453693 | 0.033298 |
| Il18r1 | -2.46523902 | 5.39E-22 |
| Egr1 | -2.459085279 | 0.00127 |
| Ccl3 | -2.452198757 | 5.38E-19 |
| Cd7 | -2.43767528 | 2.92E-06 |
| Npr2 | -2.436810739 | 0.002424 |
| Pou6f1 | -2.431499054 | 0.085329 |
| Vsig10 | -2.407992455 | 0.008965 |
| Pcdh7 | -2.407653082 | 0.09484 |
| Alox12e | -2.403182902 | 0.000851 |
| St8sia6 | -2.400063516 | 2.72E-06 |
| Samd14 | -2.397073011 | 0.033255 |
| Atp8b5 | -2.396772315 | 0.013923 |
| Gpc4 | -2.393926738 | 0.000269 |
| Fcer1a | -2.392907267 | 0.016248 |
| Dapk2 | -2.388974944 | 0.003582 |
| F5 | -2.388651979 | 8.36E-09 |
| Pla2g3 | -2.385100166 | 0.086878 |
| Gata1 | -2.37950433 | 7.52E-06 |
| Ly6d | -2.375154462 | 0.005302 |
| Gpr34 | -2.374458356 | 4.58E-19 |
| Adgrg6 | -2.37421099 | 0.011109 |
| Ppp1r3f | -2.369922472 | 0.015483 |
| Mylk | -2.369234913 | 0.007106 |
| Vmn2r84 | -2.357997846 | 0.088332 |
| Lck | -2.357945502 | 0.000165 |
| Serpinb2 | -2.34444065 | 0.000279 |
| Ctsw | -2.343761158 | 0.059667 |
| Chi3l1 | -2.343391862 | 9.12E-05 |

| | | |
|---|---|---|
| Camp | -2.340570179 | 0.087969 |
| Cd200r3 | -2.333368196 | 4.03E-20 |
| Rec8 | -2.330576281 | 0.008185 |
| Zcchc18 | -2.32317555 | 0.068229 |
| Rtn4r | -2.31912392 | 0.002736 |
| Sema7a | -2.318609051 | 0.002166 |
| Tnfrsf18 | -2.31850649 | 0.002915 |
| Lag3 | -2.310235181 | 0.000206 |
| Ccl6 | -2.3075717 | 0.000319 |
| Csrp3 | -2.303849907 | 2.63E-06 |
| Ehd3 | -2.300776362 | 0.003724 |
| A530010L1 | -2.296381214 | 0.078268 |
| Ttc39b | -2.293355393 | 2.23E-06 |
| Bcas1 | -2.290984585 | 0.016877 |
| Slc12a5 | -2.289232936 | 0.048764 |
| Pilrb1 | -2.283549586 | 0.020888 |
| Prkcq | -2.28250016 | 5.15E-07 |
| Asb2 | -2.276989057 | 0.045121 |
| Tie1 | -2.2765379 | 0.044393 |
| Mmp27 | -2.271514227 | 0.05334 |
| Tent5c | -2.270960021 | 0.000693 |
| Eya2 | -2.266886191 | 0.001645 |
| Fam110c | -2.264748301 | 0.00017 |
| Hrh4 | -2.263784409 | 0.002183 |
| Panct2 | -2.252662936 | 0.005944 |
| Gimap8 | -2.249388253 | 0.033742 |
| Ccr9 | -2.243076633 | 0.039946 |
| Card10 | -2.242410436 | 0.096162 |
| Slc24a3 | -2.238563918 | 5.44E-13 |
| Gm973 | -2.238097164 | 0.070726 |
| Marveld2 | -2.237506782 | 0.013333 |
| Gzmb | -2.23528892 | 0.001055 |
| Slc22a3 | -2.231952847 | 3.73E-08 |
| Bank1 | -2.229861178 | 0.018909 |
| Enpp5 | -2.217320803 | 0.007943 |
| Hgfac | -2.212792223 | 0.04434 |
| Trpc6 | -2.205385674 | 0.066167 |
| Fzd6 | -2.204916123 | 0.011109 |
| Il2rb | -2.202599857 | 0.010283 |
| Olfm4 | -2.201073224 | 0.035506 |
| Plxdc2 | -2.193297024 | 4.23E-05 |
| D430036J1 | -2.187738669 | 0.056834 |
| Ica1l | -2.183527193 | 0.026607 |
| Cyp11a1 | -2.182191619 | 2.27E-27 |

| | | |
|---|---|---|
| S1pr1 | -2.182045124 | 0.009285 |
| Prss34 | -2.178959387 | 3.21E-05 |
| Hoxa10 | -2.177151981 | 0.098754 |
| Col7a1 | -2.176985083 | 0.029824 |
| Alox15 | -2.176826849 | 3.59E-25 |
| Cd200r4 | -2.174066457 | 8.21E-18 |
| Jazf1 | -2.17184759 | 3.57E-05 |
| Diras2 | -2.165847362 | 0.021476 |
| Ms4a2 | -2.163989297 | 9.17E-30 |
| Htra3 | -2.163914049 | 0.0876 |
| F2r | -2.159241399 | 1.59E-05 |
| Sdsl | -2.155172591 | 0.058075 |
| Gata2 | -2.15379721 | 1.20E-06 |
| Ets1 | -2.148589037 | 1.62E-29 |
| Ctla2a | -2.145640638 | 0.017216 |
| Faah | -2.139790691 | 8.81E-14 |
| Gna14 | -2.137195373 | 0.013908 |
| St6gal1 | -2.136409797 | 0.000909 |
| Rnf43 | -2.133184826 | 3.94E-05 |
| Cdh1 | -2.131814257 | 1.30E-39 |
| Ccdc8 | -2.117142281 | 0.002256 |
| Adora2b | -2.114825246 | 3.44E-06 |
| Inpp4b | -2.111361783 | 9.93E-12 |
| Adcyap1r1 | -2.109151334 | 0.07025 |
| Patj | -2.108733699 | 0.07152 |
| Lat | -2.107858269 | 8.44E-05 |
| Aqp1 | -2.105913257 | 0.091983 |
| Aqp9 | -2.105247885 | 6.13E-20 |
| Slc45a3 | -2.104404245 | 0.018812 |
| Cpa3 | -2.103805202 | 1.13E-19 |
| Isg20 | -2.086225386 | 0.000474 |
| Ngp | -2.08421931 | 0.065348 |
| Mmp25 | -2.078795174 | 1.09E-07 |
| Pcyt1b | -2.07844788 | 0.070338 |
| Nlrp12 | -2.076896166 | 0.012511 |
| Smpd5 | -2.075636306 | 0.077464 |
| Rufy4 | -2.07228517 | 0.016062 |
| Perp | -2.071960489 | 5.49E-06 |
| Alox8 | -2.070241927 | 0.000175 |
| Fam110b | -2.065012973 | 0.070726 |
| Ighm | -2.062366261 | 0.006628 |
| Nt5e | -2.060253106 | 0.073959 |
| Slc30a2 | -2.05546291 | 0.004026 |
| Irf4 | -2.047512929 | 0.00162 |

| | | |
|---|---|---|
| Itga2 | -2.044002344 | 1.30E-05 |
| Mrgpra6 | -2.040685535 | 0.048261 |
| Car2 | -2.038122034 | 0.000182 |
| Nqo1 | -2.037913681 | 0.017464 |
| Fgf3 | -2.024439676 | 0.02242 |
| Mylk3 | -2.019686666 | 0.053348 |
| Chst13 | -2.018393625 | 3.65E-07 |
| Reps2 | -2.014655842 | 0.028813 |
| Wfdc21 | -2.013907245 | 0.063605 |
| Paqr5 | -2.013146228 | 0.022818 |
| P2ry14 | -2.007621122 | 7.63E-19 |
| Prkca | -1.987332701 | 3.41E-06 |
| Atp8b1 | -1.985216788 | 0.037483 |
| Itgb2l | -1.98190329 | 0.03647 |
| Abca13 | -1.972500265 | 0.003582 |
| Ppp1r16b | -1.968742309 | 0.004855 |
| Rab37 | -1.96635236 | 1.16E-12 |
| Tnik | -1.963400556 | 1.19E-06 |
| Cd63 | -1.962341438 | 1.34E-14 |
| Kcne3 | -1.962220106 | 0.005231 |
| Arap3 | -1.961753081 | 6.13E-20 |
| Tshz3 | -1.954990424 | 0.020647 |
| Gp1ba | -1.945471607 | 0.03351 |
| Cpne5 | -1.944402059 | 0.066807 |
| A530064D( | -1.943300396 | 0.041903 |
| P2rx1 | -1.943030113 | 5.79E-08 |
| Itk | -1.941688975 | 1.45E-07 |
| Xkrx | -1.938721912 | 0.055996 |
| Alox5 | -1.938145166 | 1.34E-14 |
| Csf2rb2 | -1.935871126 | 7.28E-39 |
| Il7r | -1.932739794 | 2.51E-06 |
| Atp1b1 | -1.922671918 | 6.74E-09 |
| Lrg1 | -1.921494213 | 0.088432 |
| Gimap5 | -1.917162149 | 0.003254 |
| Hgf | -1.915229676 | 1.30E-39 |
| AA467197 | -1.912502167 | 0.094006 |
| P2ry10 | -1.909255728 | 3.98E-09 |
| Retnlg | -1.90542431 | 4.52E-06 |
| Lama5 | -1.903326252 | 1.67E-08 |
| Ccl4 | -1.903038148 | 0.012507 |
| Myo1d | -1.901642045 | 3.27E-09 |
| Gimap6 | -1.895802971 | 2.87E-08 |
| Rusc2 | -1.891529076 | 0.03787 |
| Oas1c | -1.890045601 | 0.0772 |

| | | |
|---|---|---|
| Amigo2 | -1.885591539 | 0.091983 |
| Sytl3 | -1.883619738 | 2.41E-13 |
| Rtp4 | -1.88317699 | 0.027442 |
| Pglyrp1 | -1.88254755 | 6.98E-06 |
| Sec16b | -1.882511975 | 0.079488 |
| Ddx60 | -1.881828895 | 0.000115 |
| Nrgn | -1.86727031 | 0.016573 |
| Syngap1 | -1.866022128 | 0.000237 |
| Armcx1 | -1.865274791 | 0.045869 |
| Spin4 | -1.86207855 | 0.008167 |
| Ikzf3 | -1.858451204 | 0.000373 |
| Pilra | -1.851976444 | 0.008272 |
| Tesc | -1.850710371 | 0.060503 |
| Rpgrip1 | -1.850305726 | 0.00732 |
| Naprt | -1.849676015 | 0.001246 |
| Tfr2 | -1.845435806 | 0.002182 |
| Tmem8b | -1.843766922 | 3.10E-05 |
| Rigi | -1.841396638 | 5.57E-06 |
| Itga2b | -1.839393575 | 0.026778 |
| Otub2 | -1.836581854 | 2.84E-06 |
| Adcy6 | -1.829768424 | 0.030689 |
| Chst15 | -1.828379851 | 5.95E-09 |
| Slc6a9 | -1.826955061 | 4.61E-05 |
| Ifit1 | -1.826617611 | 0.086463 |
| Grap2 | -1.826129786 | 4.58E-05 |
| Atp8a2 | -1.821863381 | 0.011109 |
| Adgre4 | -1.818812736 | 0.00039 |
| Il5ra | -1.816620889 | 0.026568 |
| Mrgpre | -1.815706869 | 0.011802 |
| B3gat2 | -1.813707717 | 0.075243 |
| C3ar1 | -1.807879205 | 5.06E-19 |
| Ankrd33b | -1.806035103 | 0.000752 |
| Il1rl1 | -1.803211532 | 3.30E-15 |
| Ffar2 | -1.802984186 | 0.000916 |
| Slc18a2 | -1.795685196 | 1.07E-08 |
| Cd226 | -1.792820362 | 0.000726 |
| Spns2 | -1.791100717 | 2.07E-06 |
| Tet1 | -1.785559023 | 6.50E-08 |
| Letm2 | -1.784158345 | 0.054174 |
| Ecm1 | -1.78086942 | 0.000544 |
| Krba1 | -1.780277543 | 3.66E-07 |
| Fam174b | -1.777181872 | 3.57E-10 |
| Igfbp7 | -1.770494191 | 0.043614 |
| Vamp5 | -1.766106491 | 0.028528 |

| | | |
|---|---|---|
| Cd55 | -1.765866128 | 5.35E-07 |
| Adora3 | -1.765020375 | 1.32E-08 |
| Septin5 | -1.76473836 | 0.049511 |
| Cd200r1 | -1.764065786 | 8.90E-17 |
| Ptgs1 | -1.763123195 | 0.003666 |
| Smim10l2a | -1.761421032 | 0.079325 |
| Optn | -1.756581786 | 0.000109 |
| Eps8l1 | -1.756350177 | 0.01328 |
| Zfp831 | -1.753582118 | 5.68E-05 |
| Abcg3 | -1.748432365 | 0.082871 |
| Pear1 | -1.734507383 | 5.33E-05 |
| Homer2 | -1.724948971 | 0.024915 |
| Pip5k1b | -1.722665602 | 0.000375 |
| Sytl1 | -1.719068062 | 8.23E-06 |
| Abca1 | -1.716916208 | 0.031718 |
| Angpt1 | -1.704905206 | 3.71E-13 |
| Tuba8 | -1.700930365 | 1.18E-06 |
| Gabbr1 | -1.700869103 | 1.61E-11 |
| Hdc | -1.688020847 | 5.40E-19 |
| Cd276 | -1.686822871 | 0.063412 |
| Myl10 | -1.674655482 | 0.000495 |
| Clnk | -1.670085892 | 0.056834 |
| Cxcr2 | -1.66487018 | 6.66E-18 |
| P2ry1 | -1.658712019 | 1.26E-11 |
| Septin1 | -1.652051598 | 4.64E-09 |
| Tulp3 | -1.646642612 | 3.01E-06 |
| Gimap9 | -1.641229978 | 5.75E-05 |
| Hyal3 | -1.635324842 | 0.081008 |
| Rab4a | -1.631721975 | 0.05552 |
| Sytl2 | -1.62875879 | 0.076434 |
| Syne1 | -1.624183046 | 6.69E-14 |
| Siglecf | -1.617910274 | 0.000106 |
| Tle6 | -1.617646589 | 0.004017 |
| Fer | -1.61663934 | 0.000807 |
| Ptger3 | -1.613714269 | 1.10E-09 |
| Cd9 | -1.611906169 | 2.00E-11 |
| Sp140l2 | -1.61142485 | 0.004018 |
| Ikzf2 | -1.610944378 | 6.66E-18 |
| Cd274 | -1.599942388 | 2.87E-08 |
| Lpcat2 | -1.599046583 | 2.50E-05 |
| Bhlhe40 | -1.595471462 | 1.55E-05 |
| Fnbp1l | -1.594541141 | 0.054311 |
| Slain1 | -1.591904668 | 1.67E-08 |
| Csf1 | -1.590810204 | 2.04E-23 |

| | | |
|---|---|---|
| Slc7a8 | -1.590334084 | 9.70E-17 |
| Timp2 | -1.584402803 | 0.020249 |
| Osbpl5 | -1.580384955 | 3.66E-07 |
| Cd300ld | -1.567268171 | 0.051609 |
| Adgrl2 | -1.564833552 | 0.001202 |
| Hemgn | -1.56277594 | 0.024751 |
| Poln | -1.561659957 | 0.000759 |
| Slco4c1 | -1.556873073 | 0.031727 |
| Rab11fip4 | -1.554514522 | 7.59E-08 |
| Nacc2 | -1.551234625 | 0.060671 |
| Ier3 | -1.549301778 | 8.73E-07 |
| Ccl5 | -1.539920359 | 0.043886 |
| Tmem62 | -1.536280366 | 0.008559 |
| Trib1 | -1.533808154 | 0.006013 |
| Ell2 | -1.528183582 | 1.20E-06 |
| Il27ra | -1.526797753 | 0.000907 |
| Basp1 | -1.525803221 | 0.011356 |
| Ston2 | -1.524621472 | 0.002984 |
| Plekhg5 | -1.521610471 | 5.57E-06 |
| St6galnac3 | -1.519115789 | 0.020516 |
| Plbd1 | -1.51383285 | 0.010162 |
| Mcemp1 | -1.494526762 | 0.061586 |
| Axin2 | -1.494366602 | 0.067086 |
| Tmem71 | -1.486141574 | 1.07E-09 |
| Itgax | -1.480241411 | 5.15E-05 |
| Padi2 | -1.479804002 | 8.61E-33 |
| Il18rap | -1.477284742 | 4.22E-16 |
| Tarm1 | -1.476370932 | 0.098264 |
| Tbc1d4 | -1.475039688 | 2.35E-22 |
| Klf5 | -1.473853189 | 0.095383 |
| Pheta2 | -1.472389907 | 0.098876 |
| Trim30b | -1.468687186 | 0.003254 |
| Pbx1 | -1.464346177 | 0.004913 |
| Acod1 | -1.463650223 | 0.047639 |
| Dach1 | -1.463226951 | 0.00901 |
| Rflnb | -1.46219021 | 1.82E-08 |
| Sgk1 | -1.453918969 | 0.011109 |
| Samd9l | -1.449988507 | 3.93E-09 |
| Rgs1 | -1.449174485 | 0.004226 |
| Fgr | -1.443689836 | 0.00196 |
| Gnb4 | -1.443605891 | 0.00571 |
| Cass4 | -1.433682646 | 0.058184 |
| Flnb | -1.431544079 | 1.07E-09 |
| Tacstd2 | -1.417150961 | 0.004465 |

| | | |
|---|---|---|
| Ptgir | -1.416779495 | 1.51E-05 |
| Hcar2 | -1.416656188 | 0.059335 |
| Ccdc171 | -1.403250108 | 0.014038 |
| Cmklr1 | -1.403132969 | 0.019053 |
| Cables1 | -1.400903159 | 0.000166 |
| Srr | -1.400485825 | 0.023857 |
| Lilrb4a | -1.398042125 | 7.95E-16 |
| Slc15a2 | -1.390650751 | 1.28E-05 |
| Armcx6 | -1.387892247 | 0.044923 |
| C5ar2 | -1.382182632 | 0.084336 |
| Il17rb | -1.3764469 | 0.043618 |
| Adam8 | -1.370266192 | 1.07E-10 |
| Cox6a2 | -1.350951244 | 0.023706 |
| Ncs1 | -1.3492915 | 0.002255 |
| Ltb | -1.348501814 | 0.000616 |
| Tmem140 | -1.347497598 | 0.000207 |
| Cish | -1.344464161 | 0.020089 |
| Emid1 | -1.341024477 | 0.020089 |
| Prkcb | -1.340892669 | 0.007846 |
| Acss2 | -1.338407842 | 2.42E-09 |
| Rsad2 | -1.331785306 | 0.059338 |
| Septin8 | -1.32849311 | 1.22E-07 |
| Cd84 | -1.324610943 | 1.87E-15 |
| Nedd4 | -1.322825805 | 1.19E-17 |
| Smim5 | -1.319683771 | 0.064688 |
| Rcn3 | -1.313266993 | 0.016062 |
| Actn1 | -1.312579348 | 0.021088 |
| Mctp2 | -1.310149023 | 0.00051 |
| Hacd4 | -1.303005539 | 8.15E-11 |
| Auts2 | -1.296842043 | 0.019974 |
| Pla2g7 | -1.292465774 | 0.003724 |
| Tec | -1.286492719 | 7.22E-14 |
| Kynu | -1.28540324 | 0.040343 |
| Rnf125 | -1.283228565 | 0.001605 |
| Zdhhc15 | -1.278372086 | 0.052215 |
| Mindy4 | -1.276893179 | 0.092317 |
| Ino80dos | -1.275650055 | 0.010718 |
| Gsn | -1.274170536 | 1.39E-08 |
| Fpr2 | -1.270424142 | 0.092873 |
| Niban1 | -1.264041013 | 3.59E-08 |
| Card11 | -1.255676444 | 0.054311 |
| Gimap1 | -1.255082014 | 0.031686 |
| Arid3b | -1.253731876 | 0.005151 |
| Siglecg | -1.253564687 | 0.018546 |

| | | |
|---|---|---|
| Padi4 | -1.251978232 | 0.002969 |
| Atp6v0a1 | -1.250695752 | 6.42E-09 |
| B3gnt7 | -1.24844427 | 0.008559 |
| Slc6a13 | -1.239696638 | 0.054773 |
| Osm | -1.230702526 | 0.001376 |
| Cmc4 | -1.22773198 | 0.009467 |
| Rab19 | -1.225572726 | 0.000904 |
| Stx3 | -1.222885157 | 4.89E-07 |
| Far2 | -1.219775292 | 5.93E-05 |
| Adgrg1 | -1.218438292 | 8.69E-07 |
| Slc2a3 | -1.21477048 | 1.82E-08 |
| Ptpn3 | -1.214679948 | 0.000961 |
| Slc40a1 | -1.214156787 | 0.023174 |
| Anxa1 | -1.213293923 | 3.77E-07 |
| Tgfbr2 | -1.212490555 | 4.25E-09 |
| Dgat1 | -1.207926545 | 1.79E-10 |
| Tecpr1 | -1.202705007 | 1.00E-11 |
| Zfp1008 | -1.195690617 | 0.049052 |
| Tmem64 | -1.195247322 | 7.27E-06 |
| Nlrc3 | -1.194861635 | 0.006995 |
| Cd72 | -1.194381905 | 0.012245 |
| Tbc1d9 | -1.193178652 | 0.076027 |
| Rom1 | -1.190826315 | 0.010182 |
| As3mt | -1.188272953 | 0.053872 |
| Arhgap6 | -1.179545338 | 0.013673 |
| Tespa1 | -1.172845957 | 3.10E-05 |
| Gpr174 | -1.171598678 | 0.0876 |
| Ccr1 | -1.17140566 | 4.94E-19 |
| Neurl3 | -1.170070567 | 3.97E-08 |
| Slc41a3 | -1.168558105 | 0.001032 |
| Tnfsf14 | -1.164313133 | 7.08E-05 |
| Trpm4 | -1.163424313 | 0.063883 |
| Ptger4 | -1.160250473 | 4.90E-08 |
| Atxn1 | -1.155779492 | 0.0005 |
| Hip1r | -1.153449827 | 2.54E-07 |
| Cd24a | -1.148089206 | 7.48E-09 |
| Cyp4f18 | -1.143646603 | 4.25E-09 |
| Gadd45a | -1.143419396 | 0.005709 |
| Ampd3 | -1.137326589 | 0.002422 |
| Eml5 | -1.134809021 | 0.001053 |
| Stau2 | -1.133519513 | 0.06193 |
| Med12l | -1.120989155 | 0.006594 |
| Dmxl2 | -1.117045249 | 0.000147 |
| Abcg1 | -1.116226512 | 0.021788 |

| | | |
|---|---|---|
| Trp53inp1 | -1.115401826 | 0.000124 |
| Pld3 | -1.112267627 | 1.51E-05 |
| Csf2rb | -1.1079912 | 1.46E-21 |
| Card6 | -1.106507835 | 0.034965 |
| Vat1 | -1.10480992 | 1.48E-06 |
| Nabp1 | -1.103795353 | 1.06E-06 |
| Arl4c | -1.099479102 | 3.93E-05 |
| Vopp1 | -1.096975613 | 1.18E-10 |
| Grina | -1.09662753 | 4.09E-09 |
| Unc13d | -1.092109641 | 7.30E-06 |
| Bicd1 | -1.089538198 | 0.081781 |
| Kdm5b | -1.088627043 | 3.45E-08 |
| Rap1gap2 | -1.088352238 | 0.030493 |
| Crebrf | -1.087666883 | 0.000327 |
| Paqr8 | -1.086133309 | 0.00347 |
| Lgalsl | -1.084886674 | 0.065926 |
| Havcr2 | -1.082053995 | 0.094605 |
| Igf1r | -1.07203596 | 7.40E-08 |
| Mxd1 | -1.071930354 | 3.27E-09 |
| Grm6 | -1.071321893 | 0.009285 |
| Slc14a1 | -1.071137939 | 0.031307 |
| Tnfrsf26 | -1.059925036 | 0.001557 |
| Ccr7 | -1.058737765 | 0.044982 |
| Ppm1e | -1.055787085 | 0.049511 |
| Enpp4 | -1.054460797 | 3.18E-05 |
| Cep162 | -1.045859241 | 0.002738 |
| Lrrc1 | -1.037190371 | 0.000302 |
| Suox | -1.036914159 | 0.00017 |
| Plgrkt | -1.036376833 | 0.000212 |
| Fgd1 | -1.032892212 | 0.042234 |
| Tsc22d1 | -1.028886631 | 0.000176 |
| Clec2d | -1.024589704 | 1.03E-09 |
| Chpf | -1.024230541 | 0.032772 |
| Galnt6 | -1.022041738 | 3.39E-13 |
| Arsb | -1.021908689 | 1.04E-10 |
| Plcg1 | -1.007661072 | 0.006594 |
| Fut8 | -1.002980502 | 5.15E-07 |
| Tmc8 | -0.996899835 | 0.043349 |
| Spns3 | -0.995574103 | 6.96E-05 |
| Smim3 | -0.993179948 | 1.45E-08 |
| Maml2 | -0.989408653 | 0.04237 |
| Nhsl2 | -0.98917702 | 0.000474 |
| Ablim1 | -0.98857164 | 0.065741 |
| Npl | -0.988193795 | 0.003465 |

| | | |
|---|---|---|
| Spint2 | -0.986997527 | 8.23E-06 |
| Bpifc | -0.985244942 | 0.004814 |
| Plcb2 | -0.97925347 | 3.40E-06 |
| Cand2 | -0.978459529 | 0.018568 |
| Fyb1 | -0.977560424 | 3.18E-13 |
| Tmem154 | -0.977503249 | 2.27E-08 |
| Ccpg1 | -0.97677947 | 1.82E-08 |
| Cracr2a | -0.975633343 | 0.000733 |
| Zfp507 | -0.97530631 | 0.015113 |
| Nynrin | -0.973161417 | 0.01763 |
| Hlf | -0.970642073 | 0.0908 |
| Dnajc6 | -0.970609818 | 0.002449 |
| Zfp949 | -0.96720389 | 0.0772 |
| C5ar1 | -0.964659304 | 0.090566 |
| Mns1 | -0.962117158 | 4.38E-05 |
| Armcx2 | -0.960894708 | 0.018387 |
| Camk1 | -0.960727069 | 0.000175 |
| Rgs18 | -0.956247954 | 8.97E-06 |
| Mylip | -0.955615283 | 0.000212 |
| Phf1 | -0.950821869 | 0.000721 |
| Jakmip1 | -0.94730947 | 0.004387 |
| Pygl | -0.945854746 | 0.003282 |
| Tspan13 | -0.941699913 | 5.54E-08 |
| Cpne2 | -0.937114735 | 1.29E-06 |
| Pim2 | -0.936268686 | 0.000977 |
| Mkrn2 | -0.93557805 | 0.016062 |
| Spry2 | -0.927572731 | 0.004014 |
| Trem6l | -0.921089059 | 0.06982 |
| Pde8a | -0.919540878 | 0.090938 |
| Asph | -0.918741872 | 5.15E-07 |
| Tspan2 | -0.915176273 | 0.01912 |
| P2rx4 | -0.91396315 | 8.71E-10 |
| Cd244a | -0.902840465 | 2.08E-06 |
| Mgat4a | -0.897947602 | 3.01E-05 |
| Nkg7 | -0.89687176 | 0.066721 |
| Clec2i | -0.896055267 | 0.000373 |
| Tnfsf10 | -0.887707183 | 0.066178 |
| Pi16 | -0.884846308 | 0.005482 |
| Otud7b | -0.876862356 | 0.042243 |
| Pglyrp2 | -0.87468223 | 0.099805 |
| Plek | -0.872732942 | 0.000172 |
| Il15 | -0.869458302 | 0.005676 |
| Zeb1 | -0.866997467 | 0.020255 |
| Rtn4rl1 | -0.866519113 | 0.005302 |

| | | |
|---|---|---|
| Cd96 | -0.858194289 | 0.091022 |
| Entpd4b | -0.857305327 | 0.006171 |
| Btg2 | -0.850691966 | 0.003147 |
| Hcfc2 | -0.847389058 | 0.022181 |
| Cd28 | -0.845725495 | 5.25E-05 |
| Nt5c3 | -0.840351477 | 1.65E-07 |
| Rcn1 | -0.839009381 | 1.31E-05 |
| Tcp11l2 | -0.837108939 | 7.94E-05 |
| Kmo | -0.834396884 | 7.30E-06 |
| Dtnb | -0.828324407 | 0.022833 |
| Treml2 | -0.827865211 | 0.00707 |
| Tns4 | -0.826378977 | 0.066035 |
| Adgre1 | -0.826180438 | 0.030122 |
| Slc16a3 | -0.82213438 | 0.000646 |
| Edem3 | -0.821319714 | 3.87E-12 |
| Cnn2 | -0.816971333 | 3.24E-07 |
| Rtl6 | -0.81585394 | 0.023925 |
| Unkl | -0.814581992 | 0.003594 |
| Klf7 | -0.811577398 | 0.009022 |
| Carmil2 | -0.811288742 | 0.055088 |
| Mob3c | -0.810701856 | 0.017829 |
| Hcst | -0.808757006 | 0.033592 |
| Lime1 | -0.806591191 | 0.092019 |
| Carns1 | -0.802556714 | 0.083949 |
| Rabgap1l | -0.80117842 | 0.009067 |
| Casp3 | -0.800137481 | 1.39E-08 |
| Rasgrp4 | -0.799369442 | 0.070994 |
| Flt3l | -0.798853161 | 0.06193 |
| Dcaf6 | -0.797786493 | 0.000671 |
| Cxxc5 | -0.795760133 | 0.032845 |
| Tbc1d8 | -0.794906696 | 0.000616 |
| Csgalnact2 | -0.7947853 | 1.13E-09 |
| Cep19 | -0.78016286 | 0.068026 |
| Sdr39u1 | -0.779819024 | 0.004999 |
| Cnr2 | -0.778807699 | 0.034893 |
| Tent5a | -0.768780841 | 0.000169 |
| Nxpe4 | -0.768081602 | 0.071078 |
| S1pr4 | -0.766296285 | 0.000988 |
| Slco3a1 | -0.765137396 | 0.000265 |
| Mycl | -0.764947942 | 0.070199 |
| Arhgef3 | -0.76283749 | 0.006846 |
| Fyn | -0.76237354 | 0.004074 |
| Il2rg | -0.757672173 | 0.000206 |
| Nck2 | -0.755915517 | 0.010063 |

| | | |
|---|---|---|
| Calcoco1 | -0.755780912 | 0.013134 |
| Rhof | -0.755493275 | 0.013394 |
| Actmap | -0.754386357 | 0.052101 |
| Jak2 | -0.75293783 | 1.67E-08 |
| Gfod1 | -0.752724355 | 0.000321 |
| Gapt | -0.74669362 | 0.000787 |
| Neat1 | -0.745292823 | 0.002913 |
| Fry | -0.744157565 | 0.057125 |
| Rhoh | -0.736546015 | 0.021547 |
| Trf | -0.728010915 | 0.002536 |
| Arl4a | -0.727683586 | 0.009399 |
| Flot1 | -0.727662631 | 1.64E-05 |
| Rrm2b | -0.725500519 | 0.022006 |
| Casz1 | -0.721906062 | 0.053411 |
| Trim34a | -0.717786566 | 0.066909 |
| Dennd1c | -0.709986425 | 0.013043 |
| Clip2 | -0.706396047 | 0.002111 |
| Slc9a9 | -0.702229551 | 0.000977 |
| Rps6ka3 | -0.701626624 | 8.44E-05 |
| Orai2 | -0.700934611 | 1.07E-09 |
| Mfsd6 | -0.699742221 | 3.53E-05 |
| Runx1 | -0.696081304 | 1.46E-08 |
| Mocos | -0.692333546 | 0.096155 |
| Ankrd12 | -0.692181549 | 4.20E-05 |
| Nat9 | -0.686619389 | 0.055443 |
| Rab44 | -0.683116363 | 8.90E-08 |
| Kif21b | -0.682609416 | 1.79E-07 |
| Irak3 | -0.682043871 | 0.003666 |
| Pnrc1 | -0.681672062 | 0.003054 |
| Rnf11 | -0.680279108 | 0.000135 |
| Antxr2 | -0.679054378 | 0.001246 |
| H1f2 | -0.676526585 | 0.054336 |
| Sec24d | -0.675358998 | 0.025826 |
| Ankrd27 | -0.674267955 | 2.44E-05 |
| Tmcc1 | -0.67122751 | 0.01718 |
| Ntng2 | -0.669183485 | 0.098147 |
| Dapp1 | -0.668055605 | 0.000851 |
| Inafm2 | -0.665334924 | 0.052598 |
| Ormdl3 | -0.660373214 | 0.010063 |
| Sipa1l1 | -0.65933417 | 0.000187 |
| Tk2 | -0.659290017 | 0.070613 |
| Zfp518b | -0.659051109 | 0.0136 |
| Serpinb1a | -0.657914167 | 0.028528 |
| Ppp1r15a | -0.655488275 | 0.059846 |

| | | |
|---|---|---|
| Hvcn1 | -0.655292066 | 0.002572 |
| Cebpa | -0.65368915 | 3.27E-05 |
| Pafah1b3 | -0.653561254 | 0.023904 |
| Nfkbie | -0.649718461 | 0.027409 |
| Acap1 | -0.649039066 | 0.000148 |
| Creb3l2 | -0.647655113 | 0.001886 |
| Srgn | -0.645793245 | 1.18E-06 |
| Carhsp1 | -0.643882502 | 0.01718 |
| Cd27 | -0.642986908 | 0.043262 |
| Ptafr | -0.642833751 | 0.04237 |
| Tbc1d10c | -0.641331292 | 0.000886 |
| Mex3a | -0.639820895 | 0.095603 |
| Fcgr3 | -0.639528476 | 0.014519 |
| Emilin2 | -0.638720323 | 0.00024 |
| Pim1 | -0.638223113 | 0.031745 |
| Tns1 | -0.634225048 | 0.052967 |
| Fut7 | -0.633876669 | 0.006595 |
| Ypel3 | -0.630755483 | 0.03413 |
| Vezf1 | -0.627015119 | 1.00E-05 |
| Dusp5 | -0.6251399 | 0.010565 |
| Nbeal2 | -0.625013095 | 0.010408 |
| Dok2 | -0.624518781 | 0.081008 |
| Suco | -0.621469304 | 0.013426 |
| Tal1 | -0.619751732 | 0.006637 |
| Pecam1 | -0.619366639 | 0.000448 |
| Nedd9 | -0.619102039 | 0.000272 |
| Arrdc3 | -0.61430052 | 0.066178 |
| Fbxo9 | -0.611107474 | 0.005429 |
| Mboat1 | -0.611064417 | 0.092228 |
| Lgals9 | -0.610001104 | 0.006468 |
| Phf21a | -0.606474478 | 0.037749 |
| Pitpnc1 | -0.605937583 | 0.002572 |
| Il1rap | -0.604995594 | 0.008609 |
| Pea15a | -0.602803535 | 0.037911 |
| Fgd3 | -0.599513874 | 0.067995 |
| Gse1 | -0.599371728 | 0.041202 |
| Lclat1 | -0.595846851 | 0.001572 |
| Txnip | -0.595555391 | 0.001943 |
| Entpd4 | -0.594956375 | 0.096369 |
| Abtb1 | -0.593998722 | 0.001865 |
| Oxr1 | -0.587753209 | 8.80E-07 |
| Sgms1 | -0.581714244 | 0.065241 |
| Med30 | -0.580847643 | 0.023174 |
| Pafah1b1 | -0.580502878 | 0.00065 |

| | | |
|---|---|---|
| Golim4 | -0.580472715 | 0.005293 |
| Lmo4 | -0.58028327 | 0.000285 |
| Prkacb | -0.580151174 | 7.05E-07 |
| Itgb2 | -0.579472295 | 0.000845 |
| Eml2 | -0.578173528 | 0.055125 |
| Ypel5 | -0.576037997 | 0.011488 |
| Ctsd | -0.57078154 | 0.000273 |
| Pnpla6 | -0.570463369 | 0.003348 |
| Tmem123 | -0.568248497 | 5.45E-05 |
| Gba2 | -0.562269759 | 0.028637 |
| Fmnl2 | -0.560350577 | 0.05265 |
| Trim12a | -0.559509402 | 0.00738 |
| Map7 | -0.559143101 | 0.01935 |
| C1galt1 | -0.557935726 | 0.004992 |
| Parp9 | -0.557625142 | 0.091983 |
| Dnmbp | -0.557271047 | 0.037368 |
| Rsbn1l | -0.55676338 | 0.001055 |
| Zfp608 | -0.556579523 | 0.00165 |
| Nucb1 | -0.556231795 | 3.19E-06 |
| Trim12c | -0.555240695 | 0.014159 |
| Als2 | -0.553871648 | 0.094996 |
| Lrba | -0.553073992 | 0.000141 |
| Abcd3 | -0.550103059 | 0.000834 |
| Tmem106b | -0.548255888 | 0.001432 |
| Irak2 | -0.548049215 | 0.094927 |
| Cbx7 | -0.546834496 | 0.098147 |
| Asns | -0.544301521 | 0.085337 |
| Zfp36l1 | -0.544163302 | 0.030863 |
| Prickle3 | -0.543018629 | 0.076975 |
| Tnk2 | -0.54299613 | 0.011922 |
| Msi2 | -0.540815627 | 0.008716 |
| Rnf166 | -0.535403753 | 0.019765 |
| Ptprcap | -0.534242125 | 0.0445 |
| Lpin1 | -0.532103638 | 0.072312 |
| Ngly1 | -0.531160542 | 0.009271 |
| Hipk2 | -0.527607311 | 0.092953 |
| Inpp5b | -0.527491777 | 0.01718 |
| Nxpe3 | -0.526512291 | 0.070994 |
| Aff3 | -0.525349769 | 0.065716 |
| St3gal1 | -0.524080264 | 0.012822 |
| Ugcg | -0.523376771 | 0.01463 |
| Cfl2 | -0.523066244 | 0.077749 |
| Rnf144a | -0.522190456 | 0.013022 |
| Zfp36 | -0.520727296 | 0.000437 |

| | | |
|---|---|---|
| Tbrg1 | -0.520542537 | 0.04053 |
| Numb | -0.520502927 | 0.016424 |
| St3gal4 | -0.520186384 | 0.002454 |
| Wdr44 | -0.519134765 | 0.04462 |
| Eeig1 | -0.517877828 | 0.068229 |
| Ift22 | -0.51666727 | 0.094006 |
| Chd6 | -0.515652437 | 0.045121 |
| S100a11 | -0.5147373 | 0.011502 |
| Vasp | -0.513569663 | 0.004393 |
| Ptms | -0.512956484 | 0.022181 |
| Susd1 | -0.508744619 | 0.001682 |
| Arhgap4 | -0.506115622 | 0.000949 |
| Lat2 | -0.505272961 | 0.016921 |
| Anapc4 | -0.503280836 | 0.068049 |
| Pttg1ip | -0.503125179 | 0.002 |
| Gpsm3 | -0.500083992 | 0.002501 |
| Mzt1 | -0.499534881 | 0.046013 |
| Rcbtb2 | -0.498661002 | 0.000608 |
| Entpd5 | -0.497608743 | 0.010408 |
| Alox5ap | -0.497372281 | 0.010135 |
| Cemip2 | -0.49598099 | 0.066404 |
| Chst11 | -0.493720301 | 0.048738 |
| Bcl11a | -0.493700049 | 0.021067 |
| Rnasel | -0.493691762 | 0.005853 |
| Jund | -0.493495813 | 0.089855 |
| Cul7 | -0.493059718 | 0.083571 |
| Itgam | -0.492895161 | 0.089481 |
| Litaf | -0.491545688 | 0.003094 |
| Rgs2 | -0.491520241 | 0.007759 |
| Mpc2 | -0.489721121 | 0.042779 |
| Ppcdc | -0.489067803 | 0.052987 |
| Fxyd5 | -0.487238607 | 1.46E-05 |
| Fam13b | -0.486744118 | 0.000781 |
| Zyx | -0.484633655 | 0.005675 |
| Tes | -0.483738555 | 0.034715 |
| Rell1 | -0.482410661 | 0.023686 |
| Ndst2 | -0.48102576 | 0.006684 |
| Iqce | -0.481002018 | 0.027625 |
| Rgs12 | -0.480871687 | 0.073022 |
| Prr13 | -0.480709274 | 0.003131 |
| Stmp1 | -0.480393323 | 0.056968 |
| Plp2 | -0.479386159 | 0.000722 |
| Runx2 | -0.477855306 | 0.022775 |
| Dpp4 | -0.477807167 | 0.024833 |

| | | |
|---|---|---|
| Usp37 | -0.475664368 | 0.074794 |
| Cd53 | -0.47515025 | 8.33E-05 |
| Eef2k | -0.471944254 | 0.000221 |
| Cyfip2 | -0.471623786 | 0.000373 |
| Nfe2 | -0.470661617 | 0.031815 |
| Ripor2 | -0.470326865 | 0.003901 |
| Stap1 | -0.465146105 | 0.084451 |
| Decr1 | -0.463737309 | 0.059983 |
| Mindy1 | -0.460288681 | 0.061586 |
| Sema4d | -0.459758739 | 0.01882 |
| AI504432 | -0.459099226 | 0.044687 |
| Klhl18 | -0.452168909 | 0.009024 |
| Rbm43 | -0.451323578 | 0.024949 |
| Rlf | -0.450567849 | 0.057128 |
| Gpbp1l1 | -0.449904675 | 0.003901 |
| Zfp280d | -0.4496514 | 0.097689 |
| Gpcpd1 | -0.44892244 | 0.049024 |
| Bmi1 | -0.443264491 | 0.037553 |
| Slc3a2 | -0.442649682 | 0.002879 |
| Armcx3 | -0.442339225 | 0.098094 |
| Atp2a3 | -0.442087168 | 0.014562 |
| Zfyve19 | -0.440549131 | 0.0772 |
| Leprotl1 | -0.439652749 | 0.002311 |
| Atg4b | -0.439415998 | 0.009467 |
| Adk | -0.436484638 | 0.003865 |
| Adpgk | -0.434690108 | 0.028528 |
| Mcu | -0.434469639 | 0.091983 |
| Mettl9 | -0.433033653 | 0.03516 |
| Rfx5 | -0.432768413 | 0.099805 |
| Pja2 | -0.429879762 | 0.002255 |
| Zfp592 | -0.427995926 | 0.036173 |
| Pfdn5 | -0.426829266 | 0.003094 |
| Kbtbd7 | -0.426729871 | 0.07084 |
| Chd2 | -0.425513266 | 0.011319 |
| Pip4k2c | -0.424171548 | 0.050775 |
| Cd82 | -0.423701522 | 0.026853 |
| Aff4 | -0.421267653 | 0.050243 |
| Ubap1 | -0.41978856 | 0.08513 |
| Fam177a | -0.41974494 | 0.083571 |
| Ccdc127 | -0.4195622 | 0.066721 |
| Chd3 | -0.419560571 | 0.021124 |
| Rasal3 | -0.417687378 | 0.00971 |
| Trip12 | -0.41608755 | 0.00017 |
| Parp4 | -0.414305351 | 0.006013 |

| | | |
|---|---|---|
| Tgoln1 | -0.411843958 | 0.000796 |
| Birc3 | -0.411273446 | 0.046081 |
| Gab2 | -0.411258875 | 0.084451 |
| Nucb2 | -0.40906389 | 0.080055 |
| Rap1a | -0.407579705 | 0.01569 |
| Hbp1 | -0.406181758 | 0.041695 |
| Jak1 | -0.405657024 | 8.59E-05 |
| Snap23 | -0.405470513 | 0.010063 |
| Gtpbp2 | -0.401978619 | 0.086402 |
| Rb1 | -0.401478922 | 0.088432 |
| Tax1bp1 | -0.399830054 | 3.83E-05 |
| Kras | -0.398754008 | 0.038031 |
| Zdhhc18 | -0.397144933 | 0.003099 |
| Cd300lb | -0.397018858 | 0.053885 |
| Rasgrp2 | -0.396989632 | 0.069891 |
| Aftph | -0.396837428 | 0.020198 |
| Ccng2 | -0.396398392 | 0.054655 |
| Stim2 | -0.394691354 | 0.065716 |
| Stx17 | -0.390251253 | 0.072142 |
| Gnai3 | -0.389470785 | 0.000908 |
| Phf20l1 | -0.388567456 | 0.0457 |
| Vps13d | -0.386234394 | 0.085991 |
| Acvr1b | -0.385393358 | 0.079488 |
| Mob3a | -0.385038705 | 0.018484 |
| Srp54a | -0.384327864 | 0.082871 |
| I830077J02 | -0.383367538 | 0.038414 |
| Myl12b | -0.380048857 | 0.030003 |
| Cdkn1b | -0.379016596 | 0.019559 |
| Rap1b | -0.378859785 | 0.002535 |
| Krcc1 | -0.377709411 | 0.029839 |
| Wbp2 | -0.377539487 | 0.070876 |
| Tmsb4x | -0.376449886 | 0.004122 |
| Pacs1 | -0.375983194 | 0.021982 |
| Supt20 | -0.374918765 | 0.088432 |
| Relt | -0.373013227 | 0.098224 |
| Sec62 | -0.372307541 | 0.066507 |
| Acsl4 | -0.369227909 | 0.07084 |
| Paip2 | -0.369209149 | 0.008037 |
| Nipsnap2 | -0.368167738 | 0.085722 |
| Por | -0.365723651 | 0.082871 |
| Rbl2 | -0.365331744 | 0.007241 |
| Tlk1 | -0.365094552 | 0.037831 |
| Itm2b | -0.362904036 | 0.027226 |
| Zfp68 | -0.36222868 | 0.044849 |

| | | |
|---|---|---|
| Kctd10 | -0.360587523 | 0.005021 |
| Ndel1 | -0.358685218 | 0.033174 |
| Erbin | -0.357251495 | 0.005956 |
| Itm2c | -0.355898887 | 0.096586 |
| Akap9 | -0.354660664 | 0.080737 |
| Ctnna1 | -0.35090663 | 0.01979 |
| Ube2e3 | -0.350845591 | 0.099664 |
| Adam17 | -0.349584411 | 0.017753 |
| Tfrc | -0.347379171 | 0.085722 |
| Sema4a | -0.344591064 | 0.008686 |
| Atp11b | -0.34404713 | 0.007011 |
| Ero1a | -0.343805165 | 0.07575 |
| Gpr171 | -0.343263292 | 0.085722 |
| Mindy2 | -0.341905732 | 0.02652 |
| Manf | -0.341026887 | 0.052505 |
| Zbtb33 | -0.340397851 | 0.059209 |
| Gpd2 | -0.340364669 | 0.056834 |
| Acss1 | -0.338976904 | 0.042364 |
| Sri | -0.337915895 | 0.084584 |
| Vbp1 | -0.336775986 | 0.098876 |
| Cerk | -0.334489734 | 0.098147 |
| Paip1 | -0.334421006 | 0.093626 |
| Esyt2 | -0.334294148 | 0.027409 |
| Cldnd1 | -0.333083503 | 0.085722 |
| Ube2r2 | -0.332957171 | 0.097245 |
| Limd2 | -0.331990232 | 0.053778 |
| Rab11a | -0.327163006 | 0.070994 |
| Add1 | -0.325709474 | 0.006022 |
| Ankrd44 | -0.32495631 | 0.061657 |
| Tnfaip8 | -0.324437657 | 0.020089 |
| Gatad1 | -0.321675522 | 0.078275 |
| Tmem30a | -0.319924545 | 0.031398 |
| Lcp2 | -0.319817065 | 0.077983 |
| Krit1 | -0.319742295 | 0.071585 |
| Gna15 | -0.316580341 | 0.035788 |
| Tprg1l | -0.316516561 | 0.09382 |
| Atp8b2 | -0.315928 | 0.083551 |
| Oip5os1 | -0.315812373 | 0.015916 |
| Ptprc | -0.314030883 | 0.017522 |
| Efcab14 | -0.313229399 | 0.069988 |
| Os9 | -0.312535902 | 0.059209 |
| Cd47 | -0.310322813 | 0.021476 |
| Arrb2 | -0.310240605 | 0.042243 |
| Anxa11 | -0.310121692 | 0.096607 |

| | | |
|---|---|---|
| Klf6 | -0.309901572 | 0.014642 |
| Khnyn | -0.306216078 | 0.084794 |
| Nqo2 | -0.306149638 | 0.088432 |
| Dcp2 | -0.30515475 | 0.098622 |
| Ncf2 | -0.303031716 | 0.09484 |
| Igfbp4 | -0.299699086 | 0.024412 |
| Npepps | -0.298553026 | 0.096667 |
| Raf1 | -0.298177541 | 0.072398 |
| Phf6 | -0.297189635 | 0.091983 |
| Nampt | -0.294965442 | 0.018331 |
| Myl12a | -0.294675246 | 0.054466 |
| Hsp90b1 | -0.29452641 | 0.031672 |
| Heatr5a | -0.293372764 | 0.014224 |
| Mdm2 | -0.293135437 | 0.086656 |
| Tmem59 | -0.291211512 | 0.019032 |
| Cstf3 | -0.290705928 | 0.094006 |
| Tmod3 | -0.289751506 | 0.042234 |
| Hk2 | -0.286816641 | 0.039707 |
| Mtmr14 | -0.285816702 | 0.039179 |
| Saraf | -0.28504169 | 0.096099 |
| Rnf130 | -0.279953457 | 0.049794 |
| Riok3 | -0.279040663 | 0.036861 |
| Btk | -0.27895639 | 0.044489 |
| Crlf3 | -0.275893143 | 0.040363 |
| Ripor1 | -0.273680732 | 0.059338 |
| Csrp1 | -0.270228908 | 0.060892 |
| Kidins220 | -0.264836664 | 0.083949 |
| Ptprs | -0.264060606 | 0.048735 |
| Adipor1 | -0.263588592 | 0.067086 |
| Scp2 | -0.255811389 | 0.046265 |
| Calr | -0.249805095 | 0.096231 |
| Dock8 | -0.249287293 | 0.014866 |
| Ywhaq | -0.237697673 | 0.039179 |
| Anxa6 | -0.234425807 | 0.023904 |
| H3f3a | 0.168258929 | 0.095692 |
| Rpsa | 0.178007311 | 0.058075 |
| Rps8 | 0.182856927 | 0.066881 |
| Rpl10a | 0.192097572 | 0.04144 |
| Slc25a3 | 0.193517238 | 0.066035 |
| Smarca4 | 0.196356642 | 0.075243 |
| Uba52 | 0.200139206 | 0.068912 |
| Pebp1 | 0.207813451 | 0.083186 |
| Flna | 0.209900895 | 0.031815 |
| Eef2 | 0.213347429 | 0.059846 |

| | | |
|---|---|---|
| Rps14 | 0.216559447 | 0.085329 |
| Prpf19 | 0.219204038 | 0.057084 |
| Rps19 | 0.220588416 | 0.090889 |
| Tubb5 | 0.220918684 | 0.072142 |
| Serinc3 | 0.227691815 | 0.084451 |
| Rps12 | 0.232808509 | 0.012245 |
| Mcm5 | 0.233692473 | 0.024833 |
| Rcc2 | 0.237057187 | 0.024915 |
| Rpl8 | 0.239145473 | 0.077464 |
| Rad23b | 0.24040263 | 0.092663 |
| Oaz1 | 0.243390302 | 0.043089 |
| Traf7 | 0.243863702 | 0.081286 |
| Rplp0 | 0.246021044 | 0.012711 |
| Dgcr2 | 0.248879963 | 0.083551 |
| Rnf187 | 0.251652456 | 0.045121 |
| Dot1l | 0.252459196 | 0.078037 |
| Wbp11 | 0.254299955 | 0.07578 |
| U2af2 | 0.256943717 | 0.020008 |
| Coro1c | 0.257389704 | 0.031279 |
| Cox4i1 | 0.260017585 | 0.055021 |
| Hint1 | 0.264112489 | 0.084584 |
| Txnrd1 | 0.264504755 | 0.07529 |
| Ptbp1 | 0.269758833 | 0.054559 |
| Wdr26 | 0.271881583 | 0.078674 |
| Ehd4 | 0.271890907 | 0.089578 |
| Ddx19a | 0.272125305 | 0.086912 |
| Ggta1 | 0.273836837 | 0.091022 |
| Rexo1 | 0.274580123 | 0.088432 |
| Ppan | 0.275062495 | 0.075243 |
| Park7 | 0.276025056 | 0.047328 |
| Mllt1 | 0.276671749 | 0.064349 |
| Parp1 | 0.279797436 | 0.064688 |
| Noc2l | 0.280855881 | 0.067869 |
| Ermp1 | 0.283130327 | 0.086744 |
| Mafg | 0.28342193 | 0.028637 |
| Phgdh | 0.283555878 | 0.073865 |
| Rps26 | 0.285060994 | 0.025958 |
| Clptm1 | 0.285618254 | 0.065348 |
| Fasn | 0.286388069 | 0.00394 |
| Cd44 | 0.286417258 | 0.089613 |
| Srcap | 0.287052291 | 0.045856 |
| Cops4 | 0.287168369 | 0.059335 |
| Abcf2 | 0.287736611 | 0.031661 |
| Zmiz1 | 0.288006389 | 0.09392 |

| | | |
|---|---|---|
| Npm3 | 0.288225848 | 0.038112 |
| Zfp710 | 0.288962208 | 0.065348 |
| Maz | 0.289592918 | 0.05835 |
| Pabpc4 | 0.292578334 | 0.030381 |
| Pom121 | 0.293423477 | 0.069144 |
| Cyfip1 | 0.294499656 | 0.017822 |
| Anxa3 | 0.294536067 | 0.095831 |
| Unc93b1 | 0.295588082 | 0.065838 |
| Setd1a | 0.296461168 | 0.019974 |
| Lrrc41 | 0.297769176 | 0.088432 |
| Kat2a | 0.298337555 | 0.028627 |
| E2f6 | 0.298876073 | 0.092873 |
| Afg3l2 | 0.298971039 | 0.045372 |
| Pisd | 0.299218488 | 0.047302 |
| Zeb2 | 0.299550923 | 0.091958 |
| Dnajc10 | 0.300310615 | 0.095678 |
| Tet3 | 0.301174861 | 0.03788 |
| Prpf6 | 0.301485268 | 0.098224 |
| Ybx3 | 0.303733772 | 0.026356 |
| Tbrg4 | 0.305117057 | 0.061936 |
| Zfyve26 | 0.305518823 | 0.045121 |
| Dnajc11 | 0.305734724 | 0.070994 |
| Pacs2 | 0.306374599 | 0.032512 |
| Vav3 | 0.307176852 | 0.097278 |
| Ms4a6c | 0.307340733 | 0.008272 |
| Gtf3c1 | 0.308368287 | 0.018608 |
| Bid | 0.308374588 | 0.024083 |
| Gusb | 0.308550834 | 0.004159 |
| Vps11 | 0.308916005 | 0.043749 |
| Tspan14 | 0.309459844 | 0.021984 |
| Nosip | 0.309608189 | 0.06151 |
| Gpx1 | 0.309884147 | 0.049169 |
| Rpl13 | 0.310235421 | 0.010408 |
| Pold1 | 0.311380267 | 0.010408 |
| Fam168a | 0.311588943 | 0.085337 |
| Cfp | 0.312213651 | 0.089714 |
| Pkd1 | 0.313596825 | 0.032085 |
| Zfp106 | 0.313642341 | 0.055136 |
| Ctbp2 | 0.314666481 | 0.070726 |
| Hira | 0.316594046 | 0.065061 |
| Slc35b4 | 0.316655305 | 0.051362 |
| Mrps7 | 0.317071686 | 0.073175 |
| Srsf9 | 0.317837751 | 0.084584 |
| Uvrag | 0.318159899 | 0.037218 |

| | | |
|---|---|---|
| Cactin | 0.320181432 | 0.072685 |
| L3mbtl2 | 0.320228653 | 0.034965 |
| Mgat5 | 0.320517543 | 0.01715 |
| Prdx4 | 0.32098143 | 0.021476 |
| Ccdc115 | 0.321262212 | 0.065394 |
| Atp1a1 | 0.322136043 | 0.004665 |
| Heatr1 | 0.323497693 | 0.035207 |
| Trim27 | 0.324051903 | 0.088432 |
| Tm7sf3 | 0.324831658 | 0.01912 |
| Coq2 | 0.329062638 | 0.094754 |
| Sdhc | 0.330392326 | 0.045227 |
| Nolc1 | 0.331758105 | 0.075933 |
| Gnptab | 0.33191114 | 0.003239 |
| Zfp282 | 0.332581082 | 0.073022 |
| BC005537 | 0.332836036 | 0.017829 |
| Prim2 | 0.334018848 | 0.070994 |
| Atad3a | 0.334607686 | 0.026391 |
| Micos10 | 0.335013711 | 0.052307 |
| Smpd4 | 0.335538826 | 0.07578 |
| Mpeg1 | 0.335820285 | 0.020266 |
| Tuba4a | 0.335906362 | 0.003094 |
| Rnf169 | 0.336065215 | 0.0298 |
| Pkm | 0.336678405 | 0.030257 |
| Pgd | 0.337637092 | 0.09484 |
| Myo5a | 0.337874594 | 0.062791 |
| Snx12 | 0.338410112 | 0.099256 |
| Slc25a39 | 0.339705735 | 0.016573 |
| Polr3e | 0.341293586 | 0.098224 |
| Ankrd52 | 0.341720167 | 0.064688 |
| Igsf8 | 0.342430823 | 0.045358 |
| Pld4 | 0.343086673 | 0.097004 |
| Timm13 | 0.343345278 | 0.066035 |
| Pik3r5 | 0.344374887 | 0.070394 |
| Kdm2b | 0.344464407 | 0.006036 |
| Lpcat1 | 0.3452248 | 0.056861 |
| Sun1 | 0.34635881 | 0.01991 |
| Tns3 | 0.346679366 | 0.008366 |
| Rnf126 | 0.34671466 | 0.059505 |
| Larp1 | 0.348108563 | 0.049511 |
| Epn1 | 0.349693675 | 0.023563 |
| Mad1l1 | 0.349779265 | 0.098264 |
| Tysnd1 | 0.350293255 | 0.09388 |
| Mast2 | 0.350416844 | 0.06968 |
| Espl1 | 0.350473548 | 0.016186 |

| | | |
|---|---|---|
| Glrx5 | 0.350617727 | 0.05326 |
| Mapkapk2 | 0.351440492 | 0.039869 |
| Cxcr4 | 0.35202472 | 0.028163 |
| Gcn1 | 0.353440279 | 0.015113 |
| Hexa | 0.354387272 | 0.011109 |
| Tgfbrap1 | 0.354458585 | 0.039747 |
| Mybbp1a | 0.355565647 | 0.001572 |
| Sympk | 0.356059054 | 0.000825 |
| Rere | 0.356093169 | 0.001084 |
| C1qbp | 0.356256812 | 0.088432 |
| Iars2 | 0.357433344 | 0.05265 |
| Llgl1 | 0.357541273 | 0.066335 |
| Uqcr10 | 0.359758599 | 0.066454 |
| Slc25a4 | 0.3631285 | 0.09484 |
| Wdr74 | 0.36320293 | 0.048738 |
| Polr1b | 0.363543864 | 0.025826 |
| Fbxo22 | 0.364139189 | 0.021476 |
| Prep | 0.364771806 | 0.025782 |
| Pan2 | 0.367839209 | 0.043614 |
| Cst3 | 0.368017808 | 0.043635 |
| Dhx38 | 0.368416095 | 0.01217 |
| Slc7a1 | 0.369097704 | 0.042177 |
| Got2 | 0.369543392 | 0.008241 |
| Ramp1 | 0.37030437 | 0.002998 |
| Dctpp1 | 0.370365834 | 0.027272 |
| Sh2b3 | 0.370385099 | 0.023686 |
| Dohh | 0.371746684 | 0.086656 |
| Rassf4 | 0.372106249 | 6.56E-05 |
| Map3k3 | 0.37298905 | 0.026069 |
| Pwp2 | 0.374245325 | 0.041093 |
| Dennd1a | 0.376372057 | 0.040199 |
| Bak1 | 0.376469993 | 0.090112 |
| Qser1 | 0.3773416 | 0.050893 |
| Cluh | 0.377587762 | 0.049194 |
| G6pdx | 0.377780852 | 0.017829 |
| Abhd17a | 0.378185348 | 0.065727 |
| Rcc1 | 0.379740746 | 0.038031 |
| Eif1ax | 0.381516985 | 0.076243 |
| Atxn7l1 | 0.382584194 | 0.011077 |
| Rusf1 | 0.383093396 | 0.033742 |
| Rrp12 | 0.383184074 | 0.027409 |
| Sipa1l3 | 0.383206327 | 0.014833 |
| Rrp1b | 0.383817485 | 0.020478 |
| Pim3 | 0.384587906 | 0.064276 |

| | | |
|---|---|---|
| Dgkd | 0.384625961 | 0.002274 |
| Mfge8 | 0.384794208 | 0.048735 |
| Wasf2 | 0.385590226 | 0.012205 |
| Katnip | 0.386030201 | 0.089578 |
| Mrpl12 | 0.386214114 | 0.092228 |
| Nutf2 | 0.387775096 | 0.025907 |
| Gde1 | 0.38788201 | 0.069891 |
| Scrib | 0.388320069 | 0.009285 |
| Abcb8 | 0.38930498 | 0.044923 |
| Fech | 0.391735337 | 0.063738 |
| Hnrnpll | 0.392390238 | 0.043089 |
| Cd93 | 0.394854069 | 0.000484 |
| Naa25 | 0.395940941 | 0.008272 |
| Thoc6 | 0.397071385 | 0.098147 |
| Snrpd3 | 0.399884998 | 0.072142 |
| Gnaq | 0.401346943 | 0.065394 |
| Wdr77 | 0.40150377 | 0.011477 |
| Rnpep | 0.402527863 | 0.020573 |
| Srgap2 | 0.403485604 | 0.010063 |
| Mthfd1 | 0.403548105 | 0.004651 |
| Heatr6 | 0.406069531 | 0.029659 |
| Tcof1 | 0.406354026 | 0.021476 |
| Gorasp1 | 0.406471043 | 0.093781 |
| Mrpl4 | 0.407160775 | 0.041912 |
| Fbl | 0.407320154 | 0.008761 |
| Lmnb2 | 0.407488517 | 0.021476 |
| Mrps26 | 0.408329865 | 0.054437 |
| Piezo1 | 0.409420165 | 0.000446 |
| Wdr46 | 0.410491583 | 0.070338 |
| Ncor2 | 0.413404678 | 0.027578 |
| H2ax | 0.414130611 | 0.098224 |
| Bmyc | 0.414602125 | 0.059846 |
| Pkig | 0.414770475 | 0.044845 |
| Tspan5 | 0.415146678 | 0.048358 |
| Rab32 | 0.415538995 | 0.065871 |
| Sephs2 | 0.415904874 | 0.029429 |
| Stard3nl | 0.415963808 | 0.083262 |
| Hsf1 | 0.416917184 | 0.038098 |
| Wdr18 | 0.417462302 | 0.072099 |
| Gigyf1 | 0.417959631 | 0.020249 |
| Vac14 | 0.418376052 | 0.016062 |
| Rad18 | 0.418694942 | 0.077464 |
| Adss1 | 0.419371662 | 0.054666 |
| Acer3 | 0.419414752 | 0.048593 |

| | | |
|---|---|---|
| Mettl13 | 0.419913519 | 0.040363 |
| Foxo4 | 0.421008051 | 0.023686 |
| Thop1 | 0.421358718 | 0.041632 |
| Phlpp2 | 0.421606274 | 0.011668 |
| Tspan4 | 0.421829704 | 0.023525 |
| Bahcc1 | 0.422090165 | 0.004115 |
| Ccne1 | 0.423740738 | 0.073189 |
| Rcc1l | 0.424198949 | 0.069144 |
| Nagpa | 0.424738286 | 0.073175 |
| Polr1c | 0.424929316 | 0.042779 |
| Tmem109 | 0.425743438 | 0.013607 |
| Man2a1 | 0.427008916 | 0.092355 |
| Nol6 | 0.428480361 | 0.086463 |
| Sppl2b | 0.428495871 | 0.031574 |
| Tomm40l | 0.429274669 | 0.048736 |
| Nlrp3 | 0.429406228 | 0.038318 |
| Itpripl1 | 0.433757266 | 0.087222 |
| Xrcc3 | 0.434282019 | 0.085722 |
| Arhgap31 | 0.436361076 | 0.054755 |
| Mybl2 | 0.436624402 | 0.00162 |
| Ift140 | 0.437456357 | 0.016149 |
| Capg | 0.437907766 | 0.028627 |
| Lyl1 | 0.439052553 | 0.00024 |
| Tmem94 | 0.439693312 | 0.004037 |
| Atp2b1 | 0.440102349 | 0.035506 |
| Cdk5rap2 | 0.440380075 | 0.018695 |
| Csf1r | 0.440397254 | 3.96E-05 |
| Slc25a22 | 0.440525369 | 0.075143 |
| Kti12 | 0.440720115 | 0.021759 |
| Parl | 0.441926973 | 0.009168 |
| Neurl4 | 0.442601423 | 0.051629 |
| Plec | 0.443597193 | 0.000737 |
| Ahsa2 | 0.445922038 | 0.024833 |
| Apc | 0.446231732 | 0.007152 |
| Slc36a1 | 0.446252929 | 0.005438 |
| Tnfrsf1a | 0.447363669 | 0.005599 |
| Frat2 | 0.449240125 | 0.092599 |
| Klrb1f | 0.449254816 | 0.018812 |
| Sars2 | 0.449600618 | 0.063484 |
| Irf5 | 0.450502708 | 0.086901 |
| Plpbp | 0.452179051 | 0.061617 |
| Rpap1 | 0.452682065 | 0.008179 |
| Scarb1 | 0.454147516 | 0.000369 |
| Cad | 0.454531245 | 0.03788 |

| | | |
|---|---|---|
| Cds1 | 0.454552164 | 0.06193 |
| Slc16a10 | 0.454919259 | 0.009082 |
| Glg1 | 0.454944993 | 0.00295 |
| Slx4 | 0.455103942 | 0.004551 |
| Scap | 0.456532732 | 0.001501 |
| Cybb | 0.457202683 | 0.065716 |
| Dr1 | 0.459304351 | 0.04303 |
| Recql4 | 0.460814666 | 0.050678 |
| Mettl1 | 0.46141116 | 0.037947 |
| Niban2 | 0.463538096 | 0.020739 |
| Tnfrsf1b | 0.464040516 | 0.03324 |
| Ntpcr | 0.465794505 | 0.091939 |
| Ifngr1 | 0.465912383 | 0.089921 |
| Tomm20 | 0.466554781 | 0.03124 |
| Osbpl3 | 0.466579838 | 0.011109 |
| Pcyox1l | 0.467393331 | 0.006072 |
| Pxylp1 | 0.467999231 | 0.008464 |
| Cox10 | 0.46816317 | 0.091958 |
| Map3k20 | 0.468422737 | 0.012386 |
| Utp20 | 0.468612764 | 0.00546 |
| Macir | 0.469072375 | 0.002111 |
| Abl2 | 0.469635419 | 0.051923 |
| Inf2 | 0.470063634 | 0.010408 |
| Socs7 | 0.470632253 | 0.007671 |
| Stx6 | 0.470658465 | 0.053411 |
| Cenpb | 0.473445488 | 0.005836 |
| Wdfy2 | 0.473453336 | 0.014224 |
| Ncln | 0.476354683 | 0.000125 |
| Dstyk | 0.477003488 | 0.002223 |
| Atxn1l | 0.477952567 | 0.019974 |
| Atp6v0c | 0.478923762 | 0.00638 |
| Ppm1h | 0.482117879 | 0.03997 |
| Ppp1r21 | 0.482623435 | 0.000193 |
| Mfsd1 | 0.483226119 | 0.015105 |
| Polg | 0.484001145 | 0.000218 |
| Acp6 | 0.484557284 | 0.016062 |
| Csf2ra | 0.484780359 | 0.01569 |
| Mpp1 | 0.484787643 | 0.063605 |
| Lpin2 | 0.486162371 | 0.026223 |
| Ptpro | 0.487715129 | 0.015686 |
| Phka2 | 0.488202748 | 0.00136 |
| Spag5 | 0.491655644 | 0.028528 |
| Ap2a2 | 0.49192113 | 0.002895 |
| Pdf | 0.49334169 | 0.066132 |

| | | |
|---|---|---|
| Stom | 0.497688126 | 0.005707 |
| Dtx4 | 0.498606976 | 0.054666 |
| Cd68 | 0.498814258 | 0.016817 |
| Usp49 | 0.498984516 | 0.049202 |
| Noa1 | 0.499510294 | 0.059667 |
| Itga5 | 0.501496377 | 0.005935 |
| Mrps28 | 0.508694105 | 0.080512 |
| Usp36 | 0.509591914 | 0.000456 |
| Coq6 | 0.512973522 | 0.099047 |
| Cdc42ep4 | 0.513201015 | 0.082871 |
| Mosmo | 0.515586256 | 0.042327 |
| Cryl1 | 0.520691088 | 0.021476 |
| Ercc2 | 0.521051925 | 0.000247 |
| Erf | 0.52338973 | 0.03019 |
| Lrp5 | 0.526263143 | 0.000206 |
| Atrnl1 | 0.531567481 | 0.025815 |
| Tmem185b | 0.538648423 | 0.009285 |
| Arhgap12 | 0.539189769 | 0.064039 |
| Cit | 0.539957045 | 0.047018 |
| Nrp1 | 0.54079415 | 2.56E-05 |
| Rap2a | 0.54160888 | 0.022293 |
| Sell | 0.542655826 | 1.32E-08 |
| Ece2 | 0.543652779 | 0.069851 |
| Prkar2a | 0.544679576 | 0.001066 |
| Rassf8 | 0.545837007 | 0.067123 |
| Gga3 | 0.547510509 | 0.076581 |
| Iba57 | 0.547966613 | 0.093781 |
| Spopl | 0.548533349 | 0.059338 |
| Tlr2 | 0.549183364 | 0.055396 |
| Slc43a2 | 0.550132456 | 2.89E-06 |
| Sestd1 | 0.550164277 | 0.081781 |
| Sapcd2 | 0.551001254 | 0.095826 |
| Iqgap3 | 0.554076952 | 0.030499 |
| Tomm40 | 0.555095765 | 0.004037 |
| Flt3 | 0.556474447 | 0.010063 |
| Trmt61a | 0.558084235 | 0.040528 |
| Clec5a | 0.558226269 | 0.030625 |
| Rbsn | 0.558291556 | 0.057574 |
| Metrn | 0.558734162 | 0.018775 |
| Emc8 | 0.561640272 | 0.000321 |
| Nckipsd | 0.564016656 | 0.0729 |
| Tom1 | 0.564367176 | 0.030493 |
| Etv5 | 0.565738657 | 0.010255 |
| Pgp | 0.567891754 | 0.015059 |

| | | |
|---|---|---|
| Zc3h12c | 0.570370525 | 0.090959 |
| Pou2f2 | 0.572774993 | 0.008272 |
| Rhob | 0.574938909 | 0.042394 |
| Ddi2 | 0.58466939 | 5.12E-05 |
| Dhx37 | 0.585921275 | 0.012237 |
| Mpo | 0.592007868 | 0.025293 |
| Aoah | 0.593860298 | 0.05265 |
| Slc49a4 | 0.59779079 | 9.93E-05 |
| Axl | 0.598691902 | 0.066035 |
| Anxa2 | 0.604489661 | 9.67E-09 |
| Snapc4 | 0.604580003 | 0.035506 |
| Nsl1 | 0.609159671 | 0.000534 |
| Klf13 | 0.609392266 | 2.08E-06 |
| Cntrob | 0.60940769 | 0.014611 |
| Adam15 | 0.610118636 | 0.003465 |
| Ring1 | 0.610478605 | 0.055139 |
| Met | 0.610865446 | 0.001682 |
| Kctd17 | 0.61198104 | 0.059689 |
| Alg1 | 0.614532441 | 0.00224 |
| Tbc1d16 | 0.616278071 | 0.037142 |
| Bola2 | 0.61849069 | 0.022181 |
| Pik3ip1 | 0.621264464 | 0.010063 |
| Itga1 | 0.625598507 | 0.064039 |
| Btla | 0.626311071 | 0.006471 |
| Zfp568 | 0.629474432 | 0.002607 |
| Cd81 | 0.635001894 | 0.0457 |
| Dse | 0.635853003 | 0.043319 |
| Caskin2 | 0.638223673 | 0.020249 |
| App | 0.640607059 | 0.010711 |
| Urb1 | 0.644752888 | 8.22E-06 |
| Tsen54 | 0.647747119 | 0.001901 |
| B3gnt5 | 0.647967493 | 0.019974 |
| Cebpd | 0.648171527 | 0.004949 |
| Gan | 0.649838213 | 0.001193 |
| Mtus1 | 0.654878626 | 0.018695 |
| Plod1 | 0.655546825 | 0.027837 |
| Cyp4f16 | 0.658713308 | 0.000726 |
| Ydjc | 0.659174468 | 0.003724 |
| Cdc42bpb | 0.659605501 | 0.002098 |
| Tnfaip2 | 0.659703872 | 2.36E-08 |
| Mmp19 | 0.660025938 | 0.077429 |
| Lrfn4 | 0.661450524 | 0.092784 |
| Sla | 0.663629537 | 2.63E-05 |
| Apba1 | 0.66610256 | 0.00136 |

| | | |
|---|---|---|
| Sik2 | 0.667926415 | 0.001029 |
| Il13ra1 | 0.6703657 | 0.099805 |
| B4galt6 | 0.671254055 | 0.02324 |
| Clock | 0.67282999 | 0.011061 |
| Mul1 | 0.67556856 | 0.019391 |
| Wwc2 | 0.678263573 | 0.001612 |
| Cd320 | 0.68351533 | 0.061956 |
| Them6 | 0.686114911 | 0.017293 |
| Qtrt1 | 0.689450192 | 0.021912 |
| Dusp3 | 0.689896991 | 0.09484 |
| Aars2 | 0.694930069 | 0.003675 |
| Cd14 | 0.699554117 | 0.040998 |
| Fam210a | 0.708899652 | 0.000303 |
| Dusp7 | 0.713648017 | 2.30E-06 |
| Slc19a2 | 0.717760778 | 0.006183 |
| Ctnnd1 | 0.718857602 | 0.001426 |
| Spns1 | 0.719513227 | 0.001682 |
| Ahnak | 0.722661346 | 0.005853 |
| Spp1 | 0.727941725 | 0.002 |
| Ldlrad3 | 0.729610756 | 0.005211 |
| Slc25a15 | 0.72983409 | 0.024896 |
| Bmf | 0.73038192 | 0.066367 |
| Glul | 0.731550946 | 0.001321 |
| Sort1 | 0.732464758 | 3.89E-05 |
| Atg9b | 0.73632483 | 0.076467 |
| Nt5dc2 | 0.739575617 | 0.001469 |
| Bcl2l13 | 0.741275822 | 0.011319 |
| Comtd1 | 0.744600759 | 0.062578 |
| Usp45 | 0.75074951 | 4.58E-05 |
| Mycbp | 0.752690657 | 0.008328 |
| Lifr | 0.757667177 | 0.075034 |
| Thbs1 | 0.760564224 | 0.004099 |
| Pkn3 | 0.760566917 | 0.018387 |
| Sh3pxd2b | 0.761525566 | 0.002433 |
| Slc12a4 | 0.76687601 | 0.000646 |
| Klf4 | 0.768608472 | 0.00175 |
| Tbkbp1 | 0.770216386 | 0.000859 |
| Ralb | 0.777294801 | 0.000119 |
| Abcb6 | 0.778508466 | 0.001865 |
| Kcnk12 | 0.781252904 | 0.015463 |
| Thbd | 0.785914773 | 0.000394 |
| Endod1 | 0.794429736 | 0.000119 |
| Lrp1 | 0.800676493 | 0.000681 |
| Elk3 | 0.811860381 | 0.006851 |

| | | |
|---|---|---|
| Pira11 | 0.821395959 | 0.013644 |
| Tigar | 0.825975964 | 0.028528 |
| Sema6b | 0.833011665 | 0.043618 |
| Lmo1 | 0.834416967 | 3.10E-05 |
| Galc | 0.836780568 | 0.042061 |
| Syne3 | 0.846060527 | 2.97E-05 |
| Man1a2 | 0.847332213 | 0.015181 |
| Il6ra | 0.849275834 | 2.28E-11 |
| Jun | 0.857202176 | 0.010408 |
| Tmem38b | 0.857461909 | 0.099805 |
| Angptl2 | 0.858041231 | 0.024915 |
| Gpr65 | 0.868284562 | 0.000137 |
| Ptgr1 | 0.8696032 | 0.059667 |
| Gda | 0.880749634 | 0.001772 |
| Slc16a13 | 0.882535384 | 0.0078 |
| Tlr13 | 0.888506445 | 0.002166 |
| Nrg2 | 0.894262856 | 0.00027 |
| Per1 | 0.904684038 | 0.03613 |
| Tgfbi | 0.906064058 | 0.000405 |
| Gspt2 | 0.913561131 | 0.041695 |
| Scnn1a | 0.915294514 | 0.045121 |
| Clec4a2 | 0.943048249 | 0.091022 |
| Gpnmb | 0.95163159 | 0.094309 |
| Cd99l2 | 0.953687588 | 0.077464 |
| Sh3pxd2a | 0.958135731 | 0.000807 |
| Zfp775 | 0.96621024 | 0.088432 |
| Fkbp5 | 0.968092943 | 3.71E-13 |
| Coa7 | 0.982342923 | 0.005509 |
| Tfec | 1.004107381 | 0.000506 |
| Slc46a3 | 1.004833796 | 0.082644 |
| Lonrf3 | 1.011236585 | 0.002166 |
| Acot1 | 1.025997805 | 0.098876 |
| Gatb | 1.031885104 | 3.47E-06 |
| Slc35e4 | 1.034994878 | 0.000914 |
| Prss16 | 1.03621594 | 6.76E-06 |
| Abcd2 | 1.048893035 | 0.001107 |
| Gm16867 | 1.064860471 | 0.002999 |
| Maged1 | 1.081891981 | 0.08196 |
| Pcdhgc3 | 1.088626667 | 0.009399 |
| Hmox1 | 1.093349308 | 0.00571 |
| Eps8 | 1.117144774 | 0.005725 |
| Arhgef10l | 1.13236845 | 0.000255 |
| Hip1 | 1.156057704 | 6.30E-10 |
| Zfyve9 | 1.194518878 | 0.00971 |

| | | |
|---|---|---|
| Deptor | 1.194868552 | 0.001425 |
| Nav2 | 1.203824739 | 0.001264 |
| Ms4a6d | 1.227873938 | 3.66E-07 |
| Mmp28 | 1.235697261 | 0.096395 |
| Ece1 | 1.241808806 | 2.46E-05 |
| Etv4 | 1.287507668 | 0.001572 |
| Tsc22d3 | 1.289860739 | 2.35E-11 |
| Map3k6 | 1.318156392 | 0.002969 |
| Celsr3 | 1.342712214 | 0.052918 |
| F13a1 | 1.356977954 | 2.84E-10 |
| Mgst3 | 1.361825842 | 0.000369 |
| Prune2 | 1.371308732 | 0.066717 |
| Afap1l1 | 1.381733577 | 2.38E-05 |
| Stxbp6 | 1.440414284 | 0.018695 |
| Sult1a1 | 1.455323764 | 0.075143 |
| Tnks1bp1 | 1.483370627 | 0.011922 |
| Trem2 | 1.535226463 | 0.00971 |
| Nrp2 | 1.536678829 | 0.000909 |
| Trpv4 | 1.540472612 | 0.051176 |
| Arhgef37 | 1.553493495 | 0.002401 |
| Setd4 | 1.672802014 | 0.022833 |
| Gpx3 | 1.723226751 | 0.002033 |
| Lhfpl2 | 1.742495989 | 0.027409 |
| Tymp | 1.796405503 | 0.08503 |
| Gm5960 | 1.813705007 | 3.54E-08 |
| Kcng2 | 1.843783516 | 0.059338 |
| Plagl1 | 1.858178555 | 0.0952 |
| Tlr8 | 1.881180991 | 4.22E-09 |
| Lpl | 1.906169423 | 1.37E-14 |
| Klf9 | 1.932362521 | 0.00065 |
| Plxna1 | 2.042993441 | 6.80E-05 |
| Fsd1l | 2.15312308 | 0.032643 |
| Slc27a2 | 2.195158978 | 0.016115 |
| Hal | 2.238605758 | 0.086463 |
| Kcnk13 | 2.263870713 | 0.031112 |
| Ric3 | 2.291748255 | 0.06968 |
| Vcan | 2.301018767 | 0.025596 |
| Zswim9 | 2.305733198 | 0.026778 |
| Kcnn1 | 2.472370477 | 0.062667 |
| Cdh17 | 2.665681986 | 0.00065 |
| Chil3 | 3.128798246 | 0.000511 |
| Id3 | 3.878293418 | 0.000119 |
| Fn1 | 4.033684156 | 8.32E-21 |
| Myo1b | 4.044002259 | 0.045534 |

| | | |
|---|---|---|
| Tubb2b | 4.246724464 | 0.070994 |
| Iqck | 4.387126991 | 0.071336 |
| Prss53 | 4.847987533 | 0.095551 |
| Rnase2a | 5.062756672 | 0.002025 |
| H3c4 | 5.206832571 | 0.0473 |
| Spa17 | 5.245526275 | 0.056181 |
| H3c15 | 5.297869805 | 0.056973 |
| Gm12537 | 5.575994363 | 0.094996 |
| Cd5l | 5.648408475 | 0.027299 |
| Upp2 | 6.568890315 | 0.012547 |
| Fabp4 | 6.725009098 | 0.019974 |
